# The ontogeny of immune tolerance: a model of early-life secretory IgA - gut microbiome interactions

**DOI:** 10.1101/2024.05.20.594845

**Authors:** Burcu Tepekule, Ai Ing Lim, C. Jessica E. Metcalf

## Abstract

To achieve immune and microbial homeostasis during adulthood, the developing immune system must learn to identify which microbes to tolerate and which to defend against. How such ‘immune education’ unfolds remains a major knowledge gap. We address this gap by synthesizing existing literature to develop a mechanistic mathematical model representing the interplay between gut ecology and adaptive immunity in early life. Our results indicate that the inflammatory tone of the microenvironment is the mediator of information flow from pre- to post-weaning periods. We evaluate the power of postnatal fecal samples for predicting immunological trajectories and explore breastfeeding scenarios when maternal immunological conditions affect breastmilk composition. Our work establishes a quantitative basis for ’immune education’, yielding insights into questions of applied relevance.

## Introduction

The global burden of immune-mediated disease is rapidly growing [1]. Epidemiological data indicate that early life exposures are key determinants of immune-mediated diseases later in life [2], such as allergies [3], asthma [4], type 1 diabetes [5], and inflammatory bowel disease (IBD) [6]. This impact of early life is primarily attributed to interactions between the microbiome and the immune system during a critical developmental window, which enables hosts to establish tolerance to commensal bacteria [7], ensuring the maintenance of immune and microbial homeostasis into adulthood, while appropriately defending against pathogens [8,9]. When this crosstalk is perturbed, pathological imprinting may develop, characterized by excessive immune reactivity and increased susceptibility to inflammatory diseases in adulthood [10]. Both inherent microbial factors and maternal cues determine how microbe-immune interactions unfold during early life: symbiotic commensals provide metabolic products that establish regulatory pathways facilitating a balanced immune response, whereas pathogens stimulate the immune system to develop defense mechanisms. Breastmilk delivers microbes and nutrients, but also antibodies that dictate the timing and nature of bacterial antigen presentation to the infant’s developing immune system, establishing a transgenerational cycle of immune priming [11,12]. The collective influence of these processes on immune ontogeny and maturation is encapsulated by the term ’immune education’ [13].

The information available to tackle the establishment of ‘immune education’ is now considerable [14]. To date, thousands of papers have been published, ranging from observational data from human populations to experimental perturbations in animal models. However, experimental methods inevitably focus on a limited set of mechanisms, while larger scale descriptive analysis rooted in observational data may illuminate patterns, but ultimately yield verbal descriptions lacking mathematical characterization. The time is ripe to integrate these layers of evidence into a systems biology framework that formally accounts for the multiple potential interacting components. Such a foundation will open the way to generating testable hypotheses for empirical investigation based on experimental and clinical data, and further mechanistic modeling.

To this end, we introduce a mathematical framework describing the reciprocal imprinting of the human gut microbiome and the antigen-specific endogenous mucosal Secretory Immunoglobulin A (SIgA) response during the first two years of life. While immune outcomes are undoubtedly multifaceted and influenced by numerous factors, many lines of evidence indicate the importance of the SIgA response, as its reactivity to microbial antigens plays a crucial role in mediating gut immune responses and their dysregulation [15,16]. The SIgA response also integrates both the B-cell and T-cell arms of immune ontogeny, and manifests the dual functionality of the gut mucosal immune system by selectively neutralizing pathogens while tolerating commensal bacteria beneficial for immune homeostasis [17,18]. Furthermore, SIgA represents the transgenerational aspect of immune priming: maternal SIgA modulates antigen presentation while simultaneously regulating the gut community composition [11,19].

Our mechanistic modeling strategy (Fig. 1) blends flexibility with tractability in reflecting the effects of maternal factors, feeding practices, consumer resource dynamics within microbial communities, and multifunctionality of SIgA (Fig. 1B); and introduces a quasi-stochastic mathematical model of germinal center reactions embedded in a combination of ordinary and delay differential equations (Fig. 1C). We parameterize our framework using data from infant fecal samples [20–23], formally mapping microbial abundance measurements into expectations for the dynamics of the lumen and the germinal centers of the gut, contexts traditionally obscured by the impracticality of direct sampling [24] (Fig. 1A). The mechanistic model structure entails a high dimensional parameter space, but qualitative results are consistent across parameter combinations (Supplementary Materials, Global and Local Sensitivity Analyses).

**Fig 1.**
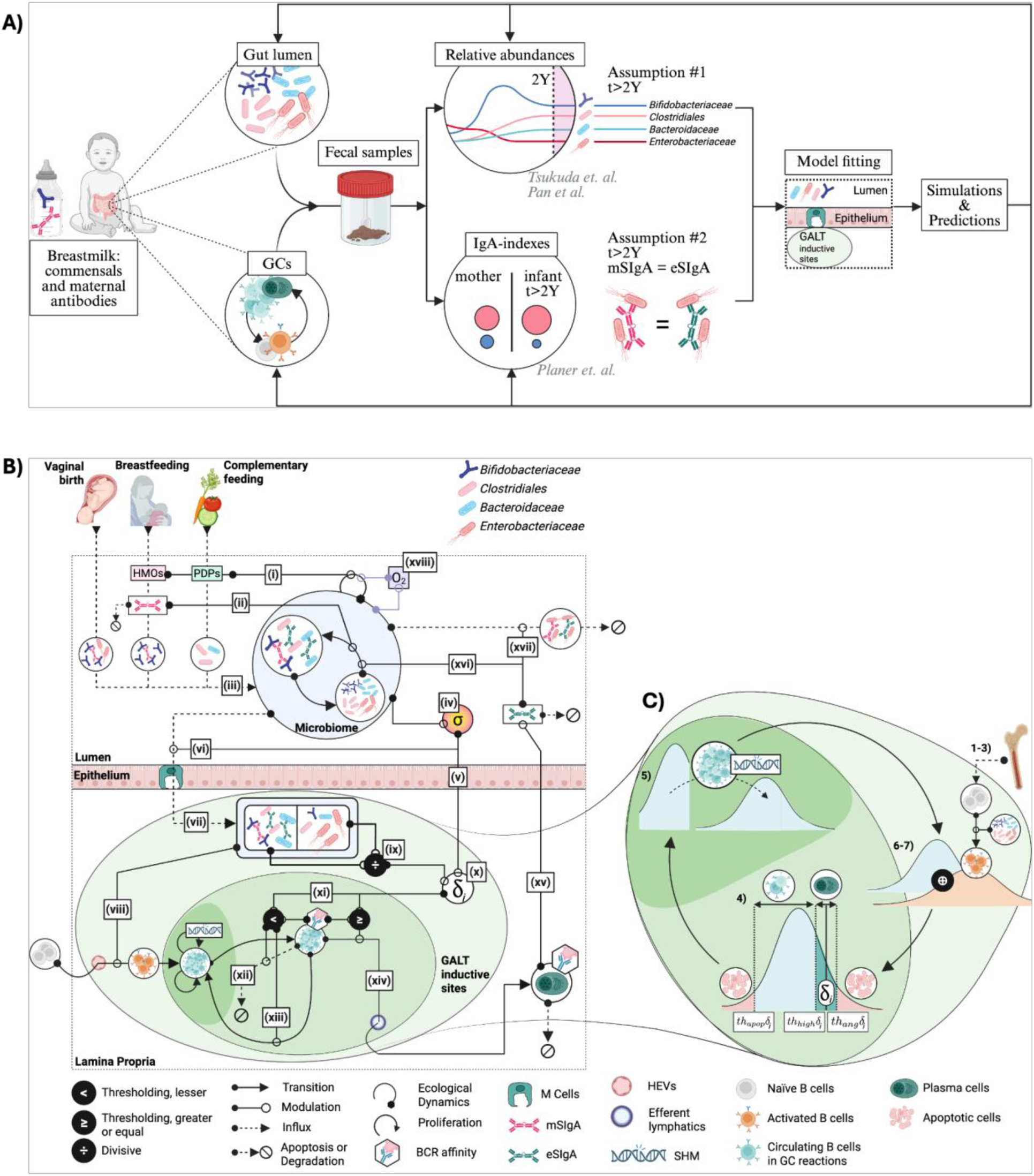
Modeling framework. **A)** Key data sources and assumptions used during model fitting, including relative abundance (line plot) and IgA-index (bubble plot) data, each colored by taxon. Model predictions reconstruct ecological dynamics in the gut lumen and the germinal center (GC) reactions. 2Y: 2 years; mSIgA and eSIgA: maternal and endogenous secretory Immunoglobulin A. **B)** Model dynamics. **(i)** Breastfeeding and complementary feeding introduces human milk oligosaccharides (HMOs), plant-derived polysaccharides (PDPs), **(ii)** mSIgA, and **(iii)** commensal bacteria to the gut lumen as SIgA-bacteria complexes or bacteria alone, alongside with the bacteria seeding the infant’s gut during vaginal birth. The microbiome within the gut lumen (large light blue circle), contains subpopulations of commensals that exist either in an SIgA-bound (white circle, left) or unbound (white circle, right) state, and transition between the two is driven by SIgA coating or degradation. **(iv)** Bacterial populations modulate the inflammatory tone of the microenvironment of the gut lumen (𝜎), **(v)** influencing epithelial cell-mediated communication with the organized gut-associated lymphoid tissue (GALT) inductive sites, and **(vi)** the antigenic sampling through barrier integrity. **(vii)** SIgA-bound and unbound antigens sampled by Microfold (M) cells **(viii)** activate the naïve B cells, and **(ix)** drive the balance between antigen-specific regulatory versus helper T cells (indicated by the divide sign). **(x)** This balance and 𝜎 shape the selection threshold 𝛿_𝑖_ for taxon 𝑖, **(xi)** which determines the BCR affinity ranges that lead to different B cell fates: **(xii)** apoptosis, **(xiii)** continued circulation, or **(xiv)** terminal differentiation into IgA-secreting plasma cells. **(xv)** IgA secreted by the plasma cells regulates the microbiome via **(xvi)** masking and **(xvii)** neutralization functions. Alongside the HMOs and PDPs, **(xviii)** the oxygen concentration (O_2_) also influences the ecological dynamics based on commensals’ metabolism. **C)** GC reactions (steps 1-7, materials and methods, corresponding to **(x)-(xiv)** on the previous panel). Naïve B cells with a wide BCR affinity distribution migrate from the bone marrow to the GALT inductive sites, and their affinity distribution is combined with that of circulating B-cells. B-cells activate upon encountering bacterial antigens. Based on how close their BCR affinities are to the selection threshold 𝛿_𝑖_, cells undergo positive selection, receiving adequate T cell help to continue circulating or differentiate to IgA-secreting plasma cells; and if not, become apoptotic. Circulating cells undergo somatic hypermutation (SHM), increasing the standard deviation of their BCR affinity distribution, continuing the cycle.

Overall, our model brings mathematical formalism to the concept of ’immune education’. This framework enables us to explore three questions with translational implications. First, it supports the quantification of potentially key diagnostic markers; second, the simulation of clinical intervention strategies, and third, identification of preventive measures before pathological trajectories are imprinted. Overall, it allows us to systematically explore the emergent properties of this complex system in a controlled, interpretable, and reproducible setting. In doing so, it opens the way to investigating persistent questions in early-life immunology with both fundamental and applied relevance.

## Results

### Model outcomes unveil the dynamics in gut lumen, successfully predicting out-of-sample data

Capturing immune education requires encompassing exogenous inputs into the gut lumen alongside endogenous dynamics in the gut lumen and organized gut-associated lymphoid tissue (GALT) inductive sites such as Peyer’s patches and colonic patches (Fig. 1A, see Table S4 for anatomical distinctions relevant to the model). Exogenous inputs quantified based on different feeding practices of infants [25–27] (Eqns. S1.1.18-S1.1.21), include caloric intake from human milk oligosaccharides (HMOs) and plant-derived polysaccharides (PDPs), and maternal secretory Immunoglobulin A (mSIgA) concentrations. HMOs and PDPs differentially modulate growth rates of different bacterial taxa based on their metabolism [28–32] (Eqn. S1.1.4). Timing of endogenous immune system activation is dictated by the maturation of Microfold (M) cells, which serve as the primary route for antigen transport to GALT inductive sites (Fig. 1B), the key site for the development of the gut adaptive immune response [33,34]. M cells are shown to appear shortly before weaning in mice models, primarily due to decreasing concentrations of maternal steroids in breastmilk [35], for which we use mSIgA as a proxy, given a lack of quantitative data. While the timing of M cell maturation in humans remains unclear, we address this uncertainty through sensitivity analysis (Supplementary Materials, Local Sensitivity Analyses). M cell maturation translates into antigenic sampling — and thus the start of immune response maturation (Fig. 2A).

**Fig 2.**
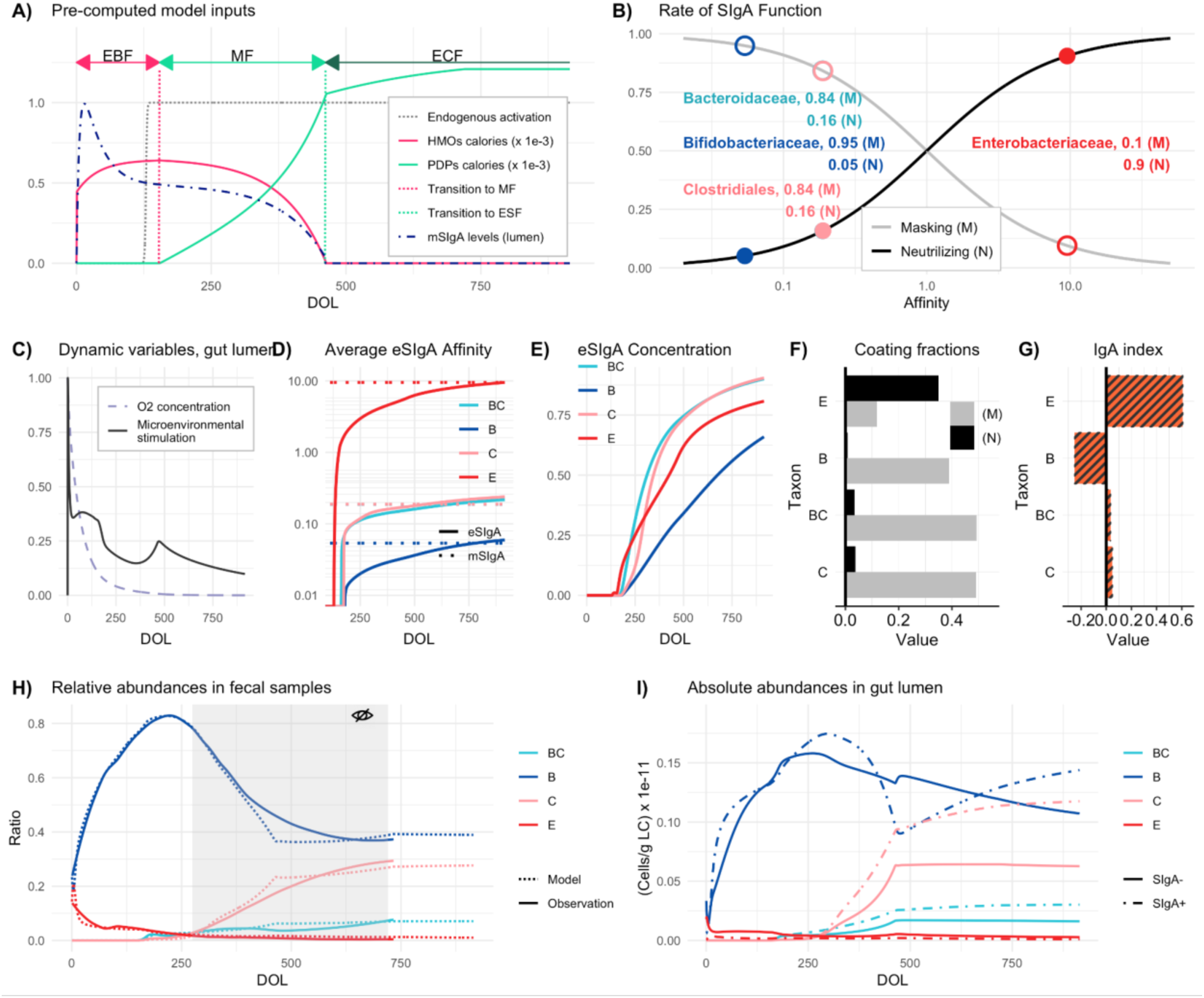
Illustration of Core Mechanisms and Model Fit. (**A**) Model inputs including normalized maternal secretory immunoglobulin A (mSIgA) concentration, human milk oligosaccharide (HMOs) and plant-derived polysaccharides (PDPs) caloric inputs, and timing of the endogenous immune system activation over time, noting that the duration of the different forms of feeding can be altered. EBM: exclusive breastfeeding; MF: mixed feeding; ECF: exclusive complementary feeding. **(B**) Rates and types of SIgA function relative to the level of antibody affinity, where mSIgA values for different taxa are highlighted. Filled and unfilled circles represent the rates of neutralizing and masking, respectively. (**C**) Normalized oxygen concentration and microenvironmental stimulation (normalized by its peak value, reflecting the inflammatory tone of the environment) over time. (**D**) Temporal progression of average endogenous SIgA (eSIgA) affinities (solid lines) gradually approaching the mSIgA affinities (dashed lines), (**E**) and the normalized antibody concentrations in the gut lumen for each key taxon. (**F**) Fractions of masked (M) and neutralized (N) bacteria and (**G**) IgA indexes for each taxon at DOL 735. (**H**) Relative abundances in fecal samples aligned with observational data, where the gray area highlights the out-of-sample data during parameter inference. (**I**) Absolute abundances in the gut lumen for key taxa distinguished by coating status. Model fit estimates for all parameters with their respective description, unit, and prior distribution are provided in Table S3. DOL: Day of life; E: *Enterobacteriaceae*; B: *Bifidobacteriaceae*; BC: *Bacteroidaceae*; C: *Clostridiales*; (M): Masking; (N): Neutralizing.

To fit our model to fecal microbiome data [20–23] and the established timeline of immune ontogeny [14,35], we leverage two observations: i) the infant develops an SIgA profile resembling their nursing mother’s [21], and ii) the infant’s immune response and microbial community structure stabilize after two years [33] (Fig. 1A). Specifically, the matured offspring secretes endogenous antibodies with quantitatively similar affinities to bacterial groups as their nursing mother; and relative abundances of bacterial taxa converge to their equilibrium values in host’s fecal samples from day 720. We estimate a single set of affinity values shared between the mother and the matured offspring, reflecting maternal antibodies in the gut lumen before activation and after the stabilization of the host’s immune response. For this step of inference, we define a composite optimization criterion that integrates the relative and absolute abundances of key taxonomic groups observed in fecal samples of infants and their IgA indexes (indicating enrichment in the SIgA coated (SIgA+) fractions). This optimization step provides estimates that remain fixed during the inference for the interim phase of endogenous immune system maturation. A second composite optimization criterion is applied to match the average SIgA coating ratio [36] in the gut lumen and the matured endogenous affinity values to the maternal ones.

To capture immunological and competitive interactions across ontogeny, we selected four bacterial taxonomic groups to reflect broad categories of host-bacterial interactions: *Enterobacteriaceae* encompasses the *Escherichia-Shigella* genus with potentially pathogenic [37,38] bacteria encountered pre-weaning when the endogenous system is more vulnerable to enteric infections. *Bifidobacteriaceae* represents early colonizers with anti-inflammatory properties that regulate the microenvironment and provide colonization resistance. *Bacteroidaceae* and *Clostridiales* are post-weaning bacteria with potentially anti-inflammatory properties that contribute to a balanced gut environment when properly regulated by SIgA (Table S2). Taxa are differentiated by two parameters: *i)* their invasiveness [39], defined as their potential to penetrate intestinal epithelial cells, and *ii)* their immunostimulatory potential, defined as their capacity to modulate the inflammatory tone of the microenvironment [40,41] via their metabolic products and epithelial conditioning [42]. While the relative ranking of these parameters across taxa is established *a priori* based on literature – for example, *Escherichia-Shigella* is assumed to have higher invasiveness and immunostimulatory potential than the rest of the taxa given its pathogenic potential – their specific values are inferred during model fitting (Table S3). Although bacterial niche (such as proximity to the epithelium [43]) could serve as an additional parameter for differentiating taxa in relation to SIgA responses, it has not been included in the model due to lack of existing quantitative data.

There is substantial evidence that the transfer of maternal antibodies to the infant gut through breastmilk plays a crucial role in shaping the neonatal gut microbiome and the immune system’s development [19,44,45]. We characterize SIgA’s functionality into two non-exclusive categories: masking (M), which represents SIgA’s non-neutralizing functionality, promoting the intestinal residency of the symbiotic commensals by limiting their interactions with the immune system [46,47]; and neutralizing (N), contributing to the expulsion of bacteria from the gut [43]. The masking (M) and neutralizing (N) rates per one unit of SIgA concentration are both modeled as a graduated response to affinity levels [48] (Eqn. S1.1.1-S1.1.2) based on the observation that SIgA targeting pathogens or pathobionts generally exhibits higher affinity compared to SIgA targeting symbiotic commensals, which often neutralizes rather than promotes their presence [49]. These rates are scaled by the binding ability of the antibodies, which we define as the strength of association between an antibody and its target antigen based on its affinity (Eqn. S1.1.3). This dual function of SIgA creates three subpopulations in the gut lumen — uncoated (SIgA-), masked (SIgA+ (M)), and neutralized (SIgA+ (N)) — of which only the uncoated and masked are metabolically active. The neutralized subpopulation neither interacts with other subpopulations nor stimulates the immune system. Because both masked and neutralized populations are coated with SIgA, techniques like IgA-Seq or BugFACS would detect binding in both cases, and the resulting SIgA coating ratio would reflect contributions from both types of bacteria. Furthermore, the functional dichotomy between high-affinity neutralizing and low-affinity masking SIgA is not absolute, as SIgA binding can influence bacterial growth or colonization in species-specific ways, involving mechanisms such as enchained growth, biofilm formation, or altered gene expression [17]. Given the scale of data available, our model does not attempt to capture these nuances (see supplementary material, section *Model Limitations*), but instead focuses on a fundamental, bacteria-agnostic mechanism, aiming to capture the core features of SIgA-microbiome interactions. Rather than relying on species-specific rules, our framework explores how affinity and functionality shift dynamically under different physiological conditions such as homeostasis and inflammation.

Our model assumes a common set of mechanisms for endogenous SIgA maturation across all commensals, allowing rates of antibody masking and neutralization to emerge from maturing affinity levels, as shaped by a backdrop of time-varying processes including developmental stages and dietary patterns (Fig. 2A). Our quasi-stochastic formulation of germinal center reactions assumes a temporally dynamic distribution of B-cell receptor (BCR) binding variability that dictates B cell fates; mean and standard deviation of binding variability are adjusted at each proliferation, somatic hypermutation, and selection (P-SHM-S) cycle (Fig. 1C). Naïve B cells migrate to the GALT inductive sites, are activated depending on the antigen uptake by M cells (Eqns. S1.1.5-S1.1.7), and start participating in GC reactions [50]. Each P-SHM-S cycle is informed by the microenvironmental stimulation in the PPs, and the selection pressure imposed by an implicit model of antigen-specific follicular helper and regulatory T cells (Materials and Methods, *Modeling the role of Germinal Centres*, and Eqn. S1.2.8), eventually determining the size and average affinity of the plasma cell pool. Microenvironmental stimulation is modeled as a composite variable (Eqn. S1.1.8) reflecting the cumulative effects of cytokines and microbial products; akin to the inflammatory tone of the local environment [51]. We infer mSIgA affinities (Fig. 2B) assuming the nursing mother with a balanced microbiome-immune response transfers masking antibodies for the symbiotic commensals (*Bifidobacteriaceae*, *Bacteroidaceae* and *Clostridiales*); and neutralizing antibodies for the pathogenic taxon *Enterobacteriaceae* [48,52–56]. Model fit estimates illustrate the trajectory of microenvironmental stimulation (Fig. 2C), average affinity of the endogenous antibodies converging toward maternal values (Fig. 2D), along with the normalized concentrations of antibody levels (Fig. 2E) proportional to the maturation of plasma cells being imprinted as the host ages.

Bacterial colonization is also affected by the oxygen concentration (O_2_) in the gut lumen (Eqn. S1.1.4). O_2_ inhibits the growth of obligate anaerobic bacteria and is consumed by facultative anaerobes (Eqn. S1.2.3) [57]. From an initially normalized level of 1, O_2_ degrades at a rate inferred during the model fitting process (Fig. 2C).

To parametrize our model, we use relative bacterial abundance and IgA-Seq data derived from infants’ fecal samples [20–23] as a proxy for the ecological and immunological dynamics within the gut lumen [58]; however, equating fecal samples with the gut lumen overlooks SIgA’s role in microbial turnover [46,47]. We reconstruct the relative abundances per fecal content by calculating the weighted sum of neutralized, coated, and uncoated subpopulations in the gut lumen. This calculation incorporates subpopulation-specific observation rates (Eqns. S1.1.14-S1.1.17), which represent the proportion of each bacterial subpopulation that successfully transfers from the gut lumen to fecal samples. These observation rates are inferred during the model fitting process and differ between neutralized, coated, and uncoated bacteria (Table S3), accounting for their varying propensities to appear in fecal samples relative to their actual abundance in the gut lumen. This approach translates the true microbial abundances in the gut lumen to the measurable abundances detected in fecal samples (Fig. 1A), allowing us to align the model output with the 16S rRNA sequencing analysis provided in *Tsukuda et al* [20]. We apply the same reconstruction using SIgA−, SIgA+(C), and SIgA+(N) subpopulations to compute the IgA indexes for each taxon, allowing us to include the IgA-seq profiles of *Enterobacteriaceae* and *Bifidobacteriaceae* provided in *Planer et al.* [21]. Our estimates indicate an average coating ratio of 48.8%, with a negative IgA index for *Bifidobacteriaceae*, slightly positive IgA indexes for *Bacteroidaceae* and *Clostridiales*, and a positive IgA index for *Enterobacteriaceae* (Fig. 2F).

Relative abundance estimates for periods of inactivity (neonatal, early postnatal) or stability (after 2 years) of the endogenous immune system demonstrate the performance of our model fitting procedure (Fig. 2G). Furthermore, out-of-sample data corresponding to the period of maturation— not used for parameter inference (Fig. 2G, gray area)—was also accurately predicted by the model. This alignment provides indirect support of the underlying assumptions regarding the immune response’s impact on microbial community dynamics, lending credibility to the model estimates of the gut lumen (Fig. 2H) for which direct data is unavailable (Fig. 1A).

Relative abundance estimates aligned with data points are depicted in Fig. S1, and additional assumptions implicit to the model structure are provided in Table S3. A shiny app is available, to allow exploration of model parameters and emergent phenomena.

### IgA-bound Enterobacteriaceae abundance can predict immune phenotypes but only at specific ontogenetic phases

Our ability to develop effective intervention strategies for preventing pathological imprinting hinges on our capacity to accurately predict the trajectory of ’immune education’ as the infant gut matures. While there are studies correlating the time course of microbiome composition and IgA-binding patterns with eventual disease susceptibility [59–61], the predictive potential of such analyses at different stages of ontogeny remains underexplored. To evaluate the potential of using taxonomic and IgA-seq profiles of fecal samples to predict the trajectory of infant’s immune maturation, we systematically explored variation in ecological and immunological outcomes by simulating our model for a total duration of 735 days across a comprehensive range of exclusive breastfeeding (EBF) and mixed feeding (MF) durations. Across 75.2% of all feeding scenarios, the host’s gut mucosal immune response matured to a tolerant profile by simultaneously developing predominantly masking SIgA (rate of masking being above 0.5, higher than rate of neutralizing) against all symbiotic commensals (*Bifidobacteriaceae*, *Bacteroidaceae*; and *Clostridiales*) (Fig. 3A), although the underlying rate of masking varied (Fig. 3B). The infant immune response diverges to a hyperreactive profile—developing predominantly neutralizing antibodies against at least one of the symbiotic commensals across 24.8% of all possible feeding scenarios (Fig. 3A). These results suggest robust convergence towards a tolerogenic profile across a wide spectrum of feeding patterns, in line with previously reported stereotypical convergence of the systemic immune response [2].

**Fig 3.**
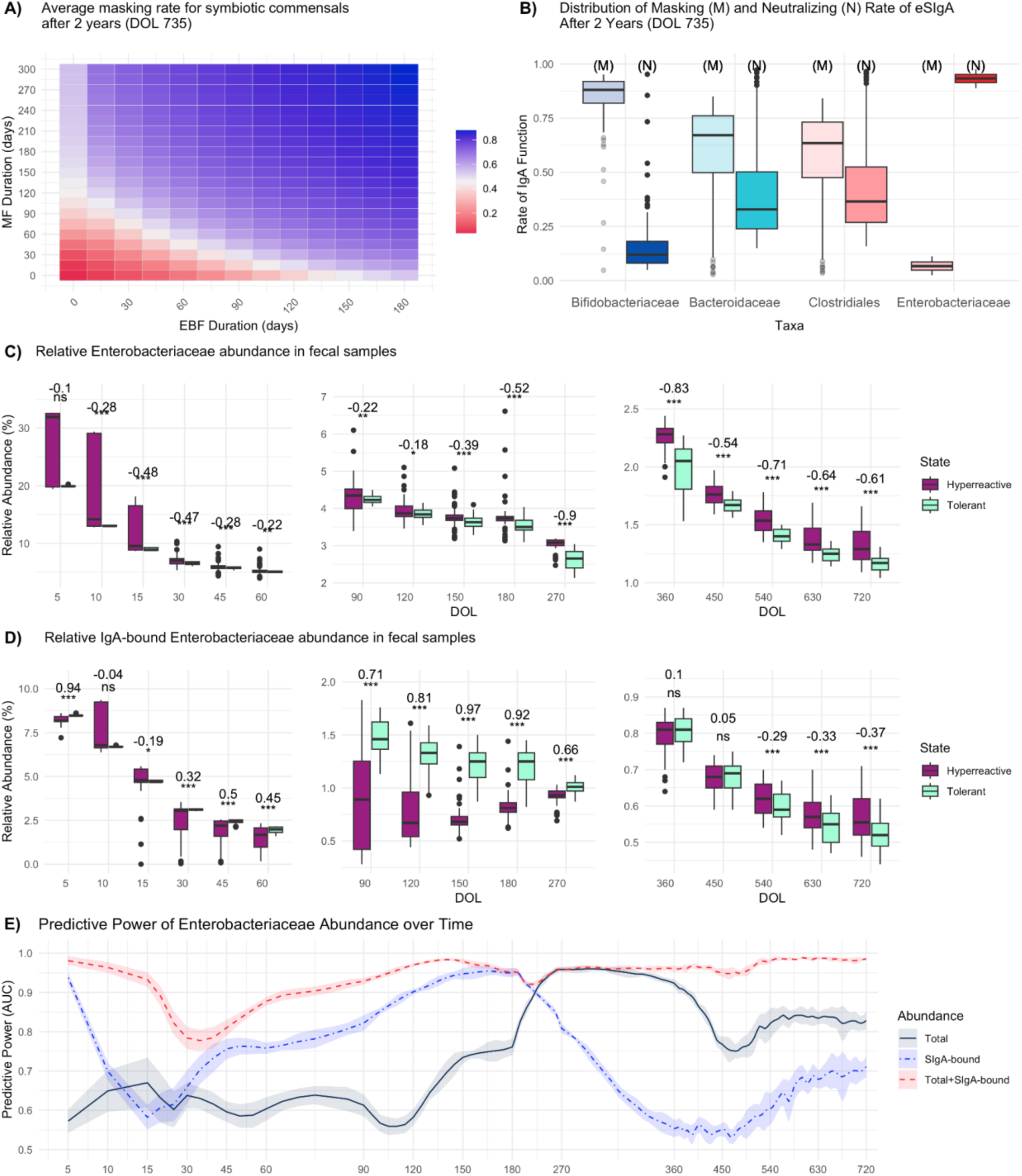
Immunological outcomes and *Enterobacteriaceae* abundances for the first 2 years of life for different combinations of exclusive breastfeeding (EBF) and mixed feeding (MF) durations. (**A**) Immune functionality across a spectrum of EBF and MF durations, where ‘tolerant’ corresponds to developing predominantly non-neutralizing but masking antibodies against all symbiotic commensals; and ’hyperreactive’ otherwise. EBF and MF durations vary from 0 to 180 and to 300 days respectively (**B**) Distribution of masking (M) and neutralizing (N) rates of endogenous SIgA at DOL 735. Distribution of (**C**) total and (**D**) SIgA-bound *Enterobacteriaceae* abundances over the first two years of life segregated by tolerant (predominantly masking SIgA for all symbiotic commensals simultaneously) and hyperreactive (predominantly neutralizing SIgA for at least one of the symbiotic commensals) states. The dataset is balanced for each time point to reflect the typical sample size of recent cohort studies [65]. A Wilcoxon rank-sum test is used to calculate the effect size (rank-biserial correlation from -1 to +1; values closer to ±1 indicating stronger associations) and the significance (*** for p < 0.001, ** for p < 0.01, * for p < 0.05, . for p < 0.1, and ns for p ≥ 0.1), displayed at the top of each boxplot. (**E**) Mean predictive power (AUC) of *Enterobacteriaceae* abundance segregated by data types. Predictive power of 0.5 indicates no better prediction of a tolerogenic/hyperreactive phenotype than random chance; 1 indicates perfect accuracy. Shaded areas represent ±1 standard error around the mean values. Values are smoothed using a local polynomial regression (LOESS) model. DOL: Day of life; AUC: Area under the receiver operating characteristic curve.

Using this synthetically generated data, we explored the diagnostic potential of early life fecal microbiome data in predicting emergent immunopathology. Specifically, we analyzed how effectively the total and SIgA-bound *Enterobacteriaceae* abundance from fecal samples collected at various developmental stages could discriminate between individuals who ultimately developed tolerance versus hyperreactivity after 2 years (at DOL 735). We focused on SIgA-bound Enterobacteriaceae as our primary marker based on the evidence that SIgA coating patterns can identify bacteria associated with immune dysregulation and inflammatory conditions [43]. We first compared the total and SIgA-bound *Enterobacteriaceae* abundance between tolerant and hyperreactive subjects at different time points throughout development. Higher levels of SIgA-bound *Enterobacteriaceae* abundance gradually flip from reflecting tolerant to reflecting hyperreactive immunological states over the time course of ontogeny (Fig. 3D). Lower SIgA-bound *Enterobacteriaceae* observed for hyperreactive phenotypes during early phases of ontogeny are indicative of insufficient *Enterobacteriaceae* neutralization by maternal antibodies. As the host matures, this pattern alters, with the hyperreactive phenotype exhibiting an increase in SIgA-bound *Enterobacteriaceae* abundance relative to the tolerant state, echoing the enrichment in SIgA-coated *Enterobacteriaceae* in adults with IBD-like phenotypes [62,63].

Building on these associations, we next asked how well total and SIgA-bound *Enterobacteriaceae* abundances at different stages of ontogeny could predict immunological phenotypes. We implemented a 5-fold cross-validation approach with stratified sampling to account for potential class imbalances in our binary outcome. For each time point, we fitted three logistic regression models including as covariates: i) total *Enterobacteriaceae* abundance, ii) SIgA-bound *Enterobacteriaceae* abundance, and iii) both total and SIgA-bound *Enterobacteriaceae* abundances. To assess and compare the predictive performance of each model, we calculated the area under the receiver operating characteristic curve (AUC) along with standard errors at each time point. This metric quantifies how well each model discriminates between the two outcome categories, with higher AUC values indicating better classification ability.

Total *Enterobacteriaceae* abundances had the lowest predictive capacity during the first 6 months of life (Fig. 3C, 2E), which can be explained by considering the two compensatory mechanisms regulating it: *i)* ecological competition preventing pathogen overgrowth and *ii)* early activation of the endogenous immune system (earliest at DOL 30) when maternal antibodies are not sufficiently neutralizing. In contrast, SIgA-bound *Enterobacteriaceae* abundance alone during the very early days of life provides valuable information regarding the immunological trajectory (Fig. 3D, 3E), as this quantity reflects the efficiency and/or presence of maternal antibodies [60]. Starting from month 6 (DOL 180), the predictive power of total *Enterobacteriaceae* abundance increases while that of SIgA-bound *Enterobacteriaceae* decreases. This shift indicates that endogenous immune responses against symbiotic commensals combined with ecological competition become the primary regulators of *Enterobacteriaceae* population, exerting stronger selection pressure than the endogenous SIgA response to *Enterobacteriaceae* itself (Fig S2). This shift in predictive powers also coincides with the transition from maternal to endogenous antibody responses. Notably, SIgA-bound *Enterobacteriaceae* samples collected during this transitional period from maternal to endogenous SIgA dominance in the gut lumen provided the least information about the host’s immunological trajectory, revealing non-monotonic temporal patterns in the predictive value of 16S rRNA and IgA-seq analyses. Comparison with an independent dataset provided in [64] showed that our model captured similar patterns in the relative importance of microbial and inflammatory predictors (Fig. S11), providing additional support to the plausibility of our results.

### Ecology versus the microenvironment: what carries the information?

An enduring mystery in mucosal immunology is how postnatal influences on the immune system persist into adulthood, even though many core cell types are not present when these initial influences occur [11]. One way to test whether this persistent influence is also reflected in our model is to investigate the impact of EBF duration on affinity maturation: antigenic sampling is delayed if M cells remain immature as a result of persistently high levels of mSIgA, and this postpones the activation of B and T cells until the end of this period. Thus, we expect the influence of the EBF period – if any – to rapidly fade unless other components of the system have enduring effects, since mSIgA has a strict half-life and the delivery of HMOs ceases instantaneously with the cessation of breastfeeding (Fig. S3A).

We first investigate the differential impacts of EBF and MF durations in determining endogenous SIgA (eSIgA) affinity across a comprehensive range of EBF and MF durations. Given the presence of pre-weaning commensals (*Bifidobacteriaceae*, *Enterobacteriaceae*) in the gut lumen during both EBF and MF, eSIgA affinity maturation towards these taxa will be influenced by the duration of both feeding practices. Conversely, a significant role for EBF duration in the affinity maturation against post-weaning commensals (*Bacteroidaceae*, *Clostridiales*) would align with the persistent postnatal influences described above. As hypothesized, immune responses against pre-weaning and post-weaning commensals are impacted by EBF and MF durations differently shown by a predictor importance analysis (Fig 4A), where we compare the relative importance of EBF and MF durations in explaining the variance in eSIgA affinities. Recapitulating the presented conundrum, the influence of EBF duration remains prominent in explaining the variance in eSIgA affinities against post-weaning commensals, despite MF duration being 1.5 times more influential. This influence is more prominent when EBF is followed by Exclusive Complementary Feeding (ECF) without any intervening MF period, which makes it challenging for the host to develop tolerance against post-weaning commensals (Fig. S3B).

**Fig 4.**
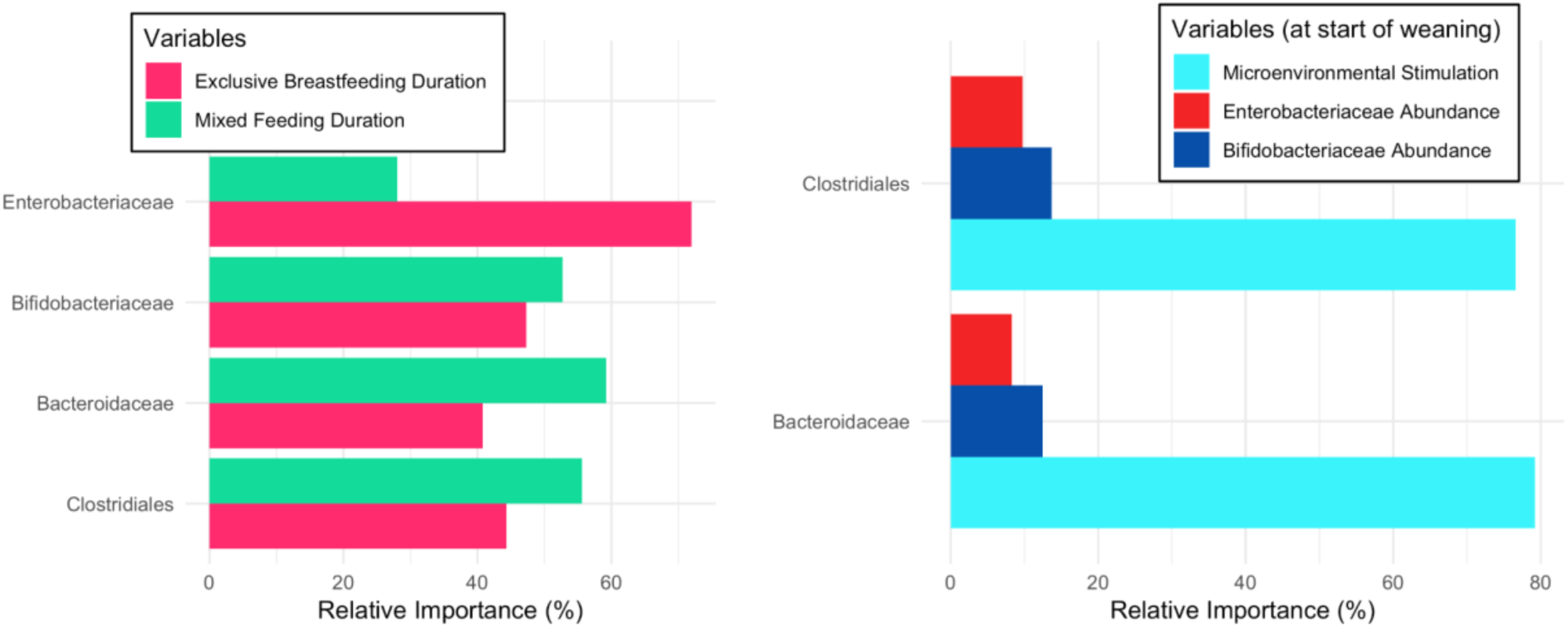
Impact of different feeding durations and environmental variables in determining the matured eSIgA values for key taxa. (**A**) Relative importance analysis in determining the eSIgA values for key taxa, comparing the impact of EBF and MF durations. (**B**) Relative importance analysis in determining the eSIgA values for post-weaning commensals, comparing the impact of the abundances of pre-weaning commensals and the microenvironmental stimulation.

Although mSIgA itself may not persist, its initial modifications may continue to shape both the ecology and the inflammatory tone of the microenvironment; similarly, HMOs may have shaped the ecology by selecting for bacteria based on their metabolic capacity, leading to a sustained temporal influence. Therefore, we compare the impact of pre-weaning commensal abundance and the microenvironmental stimulation on eSIgA affinities for post-weaning commensals to identify which components of our system carry information from the EBF period onward. We employed the Random Forests method to handle correlation between microbial abundances and microenvironmental stimulation. Intriguingly, our findings reveal a significantly greater influence of the microenvironmental stimulation over taxonomic composition (Fig. 4B). These findings suggest that the conditions and signals within the gut microenvironment—such as epithelial permeability and the tolerogenic bias of follicular T cells which are directly modulated by the microenvironmental stimulation in our model—play a pivotal role in shaping the infant’s immune response relative to the presence or abundance of specific bacterial taxa. Moreover, these results indicate that breastfeeding’s influence extends beyond immediate transmission of HMOs, antibodies, and microbes, conditioning the gut microenvironment in ways that produce effects outlasting the transient presence of such factors. While inflammatory tone and immune maturation undoubtedly have bidirectional interactions in our model making causality difficult to disentangle, our analysis provides evidence for directionality through temporal precedence: early microenvironmental conditions at the start of weaning predict subsequent immune responses against taxa that only appear post-weaning. This temporal sequence helps untangle the cyclic relationship and supports the directionality from early inflammatory tone to downstream immune maturation.

### Maternal protection and immunological imprinting in the offspring depends on mother’s immunological profile

Despite the well-established health benefits of breastfeeding, it presents a complex decision-making process for mothers experiencing immune dysregulation [66]. While epigenetic regulation of various immune cells, including dendritic cells, stromal cells, T cells, alveolar macrophages, and epithelial cells, has been implicated in the persistence of transgenerational immune priming [12,59], other mechanisms may also play a role. By meticulously eliminating other possibilities such as genetic and epigenetic factors, microbiota variation, and differences in milk-derived metabolites, recent work by Ramanan *et al.* demonstrated that the primary maternal factor allowing for transgenerational immune priming in mice is vertical transmission of maternal IgA [11]. This observation opens up the possibility of transmission of inflammatory phenotypes via maternal IgA, as well as tolerogenic ones. Indeed, evidence suggests that Inflammatory Bowel Disease (IBD) is significantly more prevalent in the offspring of IBD mothers, who tend to produce breastmilk with higher inflammatory potential and lower antibody levels [67–69]; and inadequate levels of maternal SIgA to food allergens have been linked to infantile allergic diseases in breastfed infants, showing a stronger correlation with symptoms than parental atopic history [70]. However, breastmilk’s influence goes beyond immune priming and antibody transfer, also enhancing colonization resistance against pathogens through nutritional and microbial support. Therefore, it is not immediately apparent under what circumstances the benefits of breastfeeding outweigh its risk of inflammatory priming.

To address this question, we compare five different feeding scenarios with varying maternal immunological profiles : *i)* control group recapitulating the scenario in Fig. 2, *ii)* excessive targeting of commensal bacteria by maternal antibodies (high affinity mSIgA against *Bifidobacteriaceae*, *Bacteroidaceae*, and *Clostridiales*), mimicking an IBD-like phenotype [71], *iii)* breastmilk devoid of mIgA but supplying HMOs and *Bifidobacteriaceae*, indicative of a maternal SIgA deficiency, *iv)* exclusive complementary feeding (ECF), with *Bifidobacteriaceae* introduced at a lower inoculum size and delayed introduction of post-weaning taxa (*Bacteroidaceae* and *Clostridiales*), mimicking a scenario with no breastfeeding, and *v)* ECF with *Bacteroidaceae* and *Clostridiales* transfer from the start, mimicking additional probiotic supplementation. In all scenarios, *Enterobacteriaceae* are introduced at the same inoculum size to assess the balance between pathogenic and symbiotic commensals (*Bifidobacteriaceae*, *Bacteroidaceae*, and *Clostridiales*) during the first 30 days of life, a critical period when the infant is particularly vulnerable to pathogen overgrowth.

Scenarios differed in degree of pathogen control, demonstrating two opposite trends (Fig. 5). ECF with probiotic supplementation (Fig. 5A, 5C; dashed dark purple line) and SIgA-deficient breastfeeding (Fig. 5A, 5C; dashed yellow line) initially showed an increase in the pathogenic-to-symbiotic commensal ratio followed by a decrease that brought them to a similar level by day 10, echoing their similarity in leading to Necrotizing enterocolitis (NEC) [60] and sepsis susceptibility [72]. SIgA-deficient breastfeeding had an initial advantage suggesting that HMOs and *Bifidobacteriaceae* can partially compensate for the lack of mSIgA by directly modulating the gut microbiome composition and thus the colonization resistance against *Enterobacteriaceae*. In contrast, ECF without probiotic supplementation (Fig. 5A, 5C; light green line) resulted in the highest pathogenic-to-symbiotic commensal ratio across all scenarios, emphasizing the importance of key symbiotic commensals in providing defense against pathogens. Interestingly, the second worst outcome in terms of pathogen control was observed with the IBD-like phenotype (Fig. 5 dotted magenta line), even though it provides neutralizing antibodies against the pathogenic commensal *Enterobacteriaceae*. The transfer of hyperreactive mSIgA against symbiotic commensals allowed for excessive *Enterobacteriaceae* overgrowth by neutralizing *Bifidobacteriaceae*, leading to significantly higher relative abundances of SIgA-bound *Enterobacteriaceae* in the fecal samples compared to the control group (Fig. 5B).

**Fig 5.**
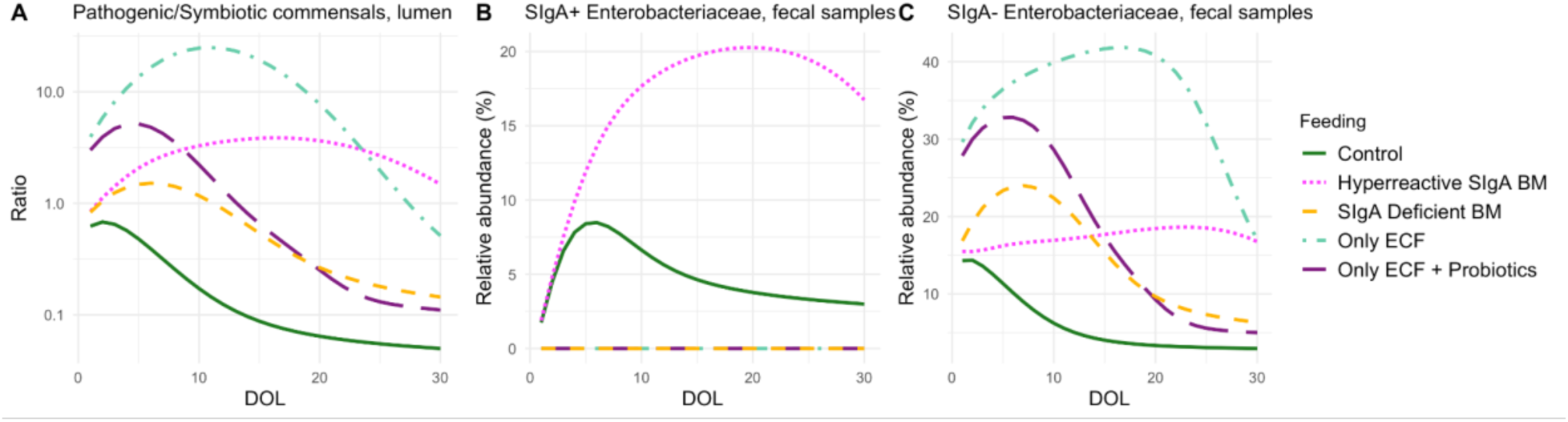
Comparison of various breastfeeding scenarios and their impact on *Enterobacteriaceae* growth. Temporal progression of (**A**) Pathogenic-to-symbiotic commensal ratio in the gut lumen, (**B**) relative SIgA-bound (SIgA+) *Enterobacteriaceae* abundance in fecal samples, and (**C**) relative SIgA-free (SIgA-) *Enterobacteriaceae* abundance in fecal samples of the infant for the first 30 days of life (DOL) for four different breastfeeding scenarios : Control, breastmilk (BM) with hyperreactive mSIgA, mSIgA deficient BM, and only exclusive complementary feeding (ECF).

In all scenarios where breastmilk lacked mSIgA, we observed hyperreactive eSIgA maturation in the infant, highlighting the influence of breastmilk antibody levels in immune imprinting (Table 1, Fig. S4), consistent with findings in the literature [67–70]. In terms of the transgenerational transmission of IgA affinity profiles, our model demonstrates a modulation of hyperreactivity against symbiotic commensals rather than complete transfer of maternal profiles (Table 1, Fig. S4). Offspring hyperreactivity against symbiotic commensals is attenuated relative to maternal hyperreactivity through both antigen-specific and antigen-agnostic mechanisms. In the antigen-specific pathway, the invasiveness of individual bacterial taxa influences offspring developing immune responses, modulated by the limited masking capacity of high-affinity mSIgA in breastmilk (Fig S7). This limited mSIgA masking capacity, combined with the M cells’ selective bias for sampling IgA-bacteria complexes [73,74] promotes the accumulation of IgA-coated bacterial antigens in the GALT inductive sites, which activates tolerogenic dendritic cells [75] and dampens the development of high-affinity neutralizing antibodies against symbiotic commensals (Fig 6B, C and D). Simultaneously, an antigen-agnostic pathway operates through microenvironmental stimulation, driven by the cumulative inflammatory capacity of unmasked bacterial subpopulations, with mSIgA’s limited masking capacity contributing to a minor reduction in inflammation. Unlike symbiotic commensals, pathogenic bacteria are inherently more invasive, making the mSIgA affinity and the IgA-mediated M cell sampling bias less influential for the taxonomic group *Enterobacteriaceae* in our model (Fig. 6A). The complete transmission of tolerance and the attenuated transmission of hyperreactivity suggest an overall beneficial effect of breastfeeding for developing tolerance against symbiotic commensals, even when the mother has hyperreactive mSIgA in her breastmilk which leads to a suboptimal endogenous antibody development (Table 1, row “Hyperreactive SIgA BM”). This attenuated transmission likely provides a mechanism for gradual normalization of immune responses across generations. However, it is important to note that this result of attenuated transmission of hyperreactivity does not account for potential contributions from maternally derived cytokines (e.g., IL-1β, IL-6, TNF-α, TGF-β) or other mediators transferred through the placenta or breastmilk [76].

**Fig 6.**
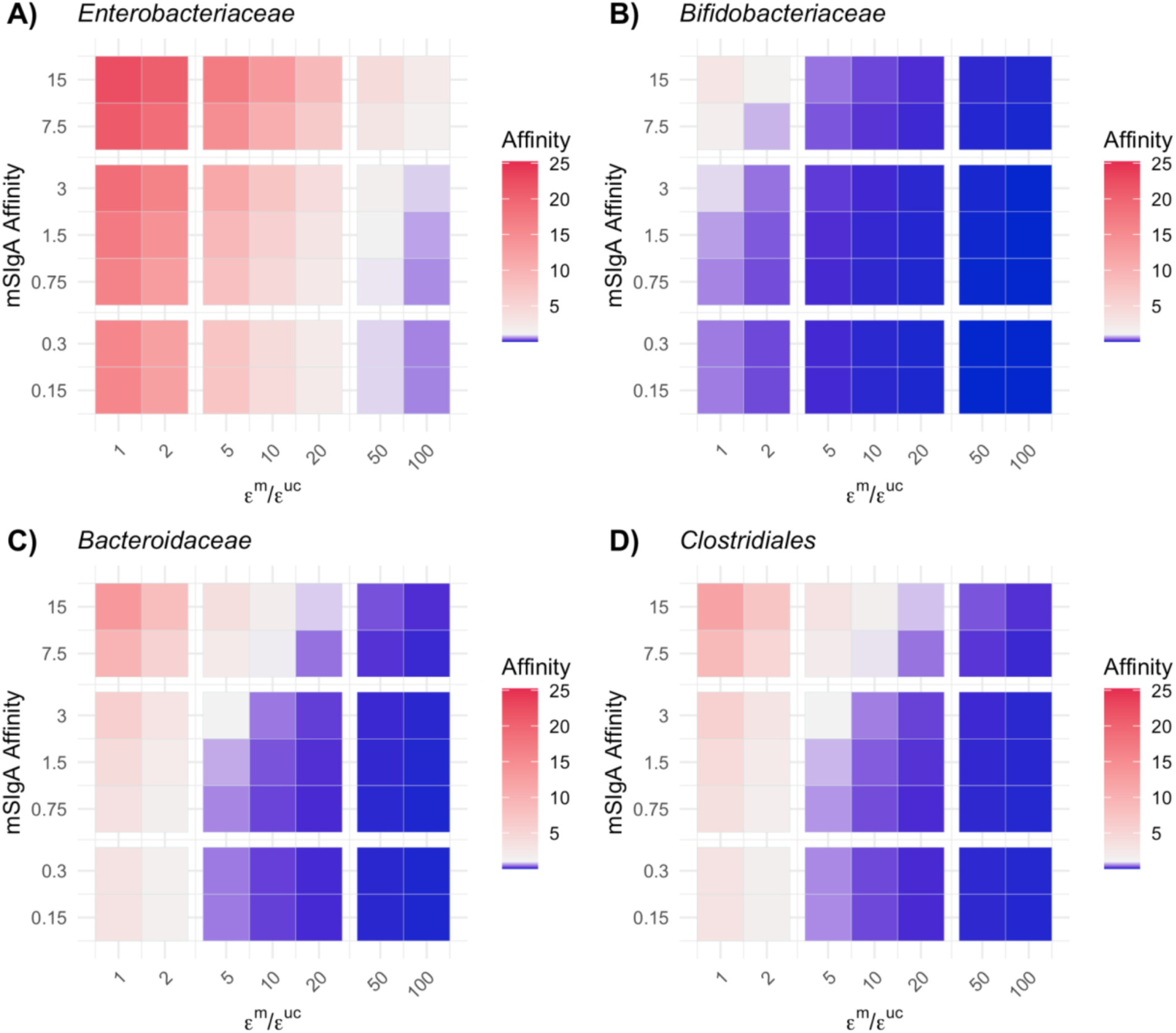
Combined influence of maternal SIgA affinity and antigenic sampling rates on the endogenous SIgA affinities. Heatmaps show endogenous SIgA (eSIgA) affinity levels at two years (DOL 735) across combinations of maternal SIgA (mSIgA) affinities and the ratio of M-cell antigenic sampling rates for masked versus uncoated bacterial antigens (represented by 𝜖^𝑚^/𝜖^𝑢𝑐^, where 𝜖^𝑚^ and 𝜖^𝑢𝑐^ are the antigenic-sampling rate of masked and uncoated bacterial antigens by M cells, respectively) for **A)** *Enterobacteriaceae*, **B)** *Bifidobacteriaceae*, **C)** *Bacteroidaceae*, and **D)** *Clostridiales*. Color gradients reflect the magnitude of eSIgA affinity: red indicates values greater than 1 (predominantly neutralizing eSIgA), white corresponds to 1, and blue indicates values below 1 (predominantly masking eSIgA).

**Table 1.** SIgA profiles in the mother and the offspring for various breastfeeding scenarios. Each column shows the neutralizing (N) rates of SIgA for each taxon, scenario, and host (mother or offspring) combination, as analyzed in Fig. 5. Cell colors indicate the magnitude of the neutralizing (N) rates, with red representing higher values, blue representing lower values, and white indicating zero – aligned with the color scheme in Fig. 6. Note that entries are left blank for scenarios in which the mother either does not breastfeed or has IgA deficiency. Rows for the scenario “Hyperreactive SIgA BM” demonstrate the attenuated transmission of SIgA hyperreactivity from the mother to the offspring. The level of maternal SIgA hyperreactivity is attenuated 82% for *Bifidobacteriaceae* (maternal neutralizing rate of 0.9 translates to a neutralizing rate of 0.16 in the offspring), and 40% for *Bacteroidaceae* and *Clostridiales* (maternal neutralizing rate of 0.9 translates to a neutralizing rates of 0.54 and 0.53 in the offspring, respectively).

|  |  | Neutralizing (N) rates of SIgA for each taxon |  |  |  |
| --- | --- | --- | --- | --- | --- |
|  |  | *Enterobacteriaceae* | *Bifidobacteriaceae* | *Bacteroidaceae* | *Clostridiales* |
| Scenario | Host |  |  |  |  |
| Control | Mother | 0.90 | 0.05 | 0.16 | 0.16 |
| Control | Offspring | 0.89 | 0.05 | 0.17 | 0.18 |
| Hyperreactive SIgA BM | Mother | 0.90 | 0.90 | 0.90 | 0.90 |
| Hyperreactive SIgA BM | Offspring | 0.92 | 0.16 | 0.54 | 0.53 |
| SIgA Deficient BM | Mother | – | – | – | – |
| SIgA Deficient BM | Offspring | 0.98 | 0.97 | 0.97 | 0.97 |
| Only ECF | Mother | – | – | – | – |
| Only ECF | Offspring | 0.98 | 0.98 | 0.96 | 0.88 |
| Only ECF + Probiotics | Mother | – | – | – | – |
| Only ECF + Probiotics | Offspring | 0.98 | 0.95 | 0.97 | 0.97 |

## Discussion

‘Immune education’ has proved a powerful concept in understanding the long shadow of early life on health outcomes. Yet, to date, the concept has remained largely qualitative, informally encompassing a wide range of individual components. This diversity and complexity of interactions make integrating these effects by experimental methods alone intractable, quite apart from the question of the degree to which animal models translate to humans. We take the first steps towards formalizing the concept of immune education in the context of gut mucosal SIgA -microbiome interactions, leveraging the power of mathematical frameworks to robustly integrate known feedback loops connecting the ecology and immunology of the gut, and inferring core parameters using data on early life immunity and microbial community dynamics.

Our analysis offers novel qualitative insights that, while grounded in our mechanistic formulations and model assumptions, may inform clinically relevant hypotheses. First, it adds a predictive dimension to the interpretation of gut microbial composition data. Traditional studies, employing 16S rRNA and IgA-seq techniques on fecal samples, have primarily focused on identifying associations between early-life microbiome compositions and later immune-mediated health outcomes. We add mechanistic understanding as to why and how the IgA coating patterns vary over the course of ontogeny by disentangling the influences of maternal and endogenous antibodies. We show that IgA-seq analysis of fecal samples collected during the transitional period—from the gradual decline of maternal antibodies to the rise of endogenous SIgA—has limited predictive value for immunological outcomes. We also reveal opposing temporal trends: reduced SIgA-bound *Enterobacteriaceae* in early ontogeny indicates insufficient maternal neutralization and predicts hyperreactivity, while the same pattern later suggests tolerogenic imprinting. These non-monotonic temporal patterns suggest that the timing of sample collection critically shapes the ability of 16S rRNA and IgA-seq data to predict long-term immune phenotypes. Moreover, they emphasize the importance of timing in study design and clinical sampling strategies, particularly when sample availability or resources are limited. These findings can be empirically tested through longitudinal studies that assess whether the predictive power of IgA-seq varies systematically across ontogeny, potentially revealing optimal sampling windows for forecasting immune-mediated conditions.

Mixed feeding duration emerges as a critical determinant of the infant’s immune trajectory: the sustained presence of maternal antibodies after weaning enables the introduction of novel antigens to the immune system in a non-inflammatory manner, promoting the development of tolerance. Exclusive breastfeeding continues to shape the immune response against microbes introduced even after its cessation by its remnant effects on the inflammatory tone within the gut lumen and GALT inductive sites. This emergent phenomenon suggests a resolution to the perennial question of how information from the early postnatal phase propagates to further stages of immune development: our model indicates that the information is not specifically carried by either symbiotic or pathogenic taxa, but, rather, results from the cumulative capacity of their SIgA-coated and uncoated subpopulations to modulate the inflammatory tone of the microenvironment.

Beyond these fundamental insights, our analysis systematically explores the applied question of how to approach breastfeeding when maternal immunological conditions such as allergies, IBD, or SIgA deficiency affect breastmilk composition. Both lack of SIgA in breastmilk, or excessive targeting of symbiotic commensals by SIgA can lead to suboptimal immune imprinting in the infant, indicating that maternal SIgA is one of the key mechanisms driving the transgenerational transmission of immunological phenotypes. The absence of breastmilk SIgA leads to a fully hyperreactive endogenous SIgA maturation, while hyperreactive maternal SIgA results in an attenuated rather than complete transmission of the inflammatory phenotype. This attenuation occurs through the combination of limited masking by high-affinity maternal SIgA, selective bias of M cells for sampling IgA-bacteria complexes, and the inherently low invasiveness of commensal microbes. Although transfer of *Bifidobacteriaceae* and HMOs cannot entirely compensate for maternal SIgA’s role in tolerogenic imprinting, it can partially compensate for the absence of SIgA in pathogen control.

Our findings further provide an evolutionary perspective on the maternal contribution to offspring immune development through breastfeeding. While the absence of breastmilk SIgA leads to significantly impaired immune regulation, our model demonstrates that even “suboptimal” maternal SIgA profiles confer substantial benefits compared to no breastfeeding. This highlights the dual functionality of SIgA: first as an antigen-specific mediator of tolerance, and second as an antigen-agnostic molecule that promotes M cell-mediated antigen sampling [73,74] and programs dendritic cells toward tolerogenic phenotypes [75]. When both functions operate optimally, complete transgenerational transfer of tolerance occurs. However, even when antigen-specific tolerance is impaired (as in maternal inflammatory conditions), the antigen-agnostic properties remain effective, resulting in attenuated rather than complete transmission of maternal hyperreactivity. This mechanism allows for the gradual dilution of inflammatory phenotypes across generations, potentially explaining why some inflammatory conditions do not follow simple vertical transmission patterns despite breastfeeding. Such a system provides evolutionary resilience, allowing for recovery from dysregulated immune states over generational timescales rather than perpetuating inflammatory phenotypes indefinitely.

These findings, combined with the influence of excess microenvironmental stimulation on endogenous SIgA affinity maturation suggests the possibility of alternative therapeutic approaches for promoting tolerogenic imprinting when maternal immunity is hyperreactive or insufficient. Interventions could focus on strategies to modulate gut inflammation beyond probiotic supplementation, for example targeting Toll-like receptors such as TLR4, which has been proposed to tackle preterm birth and fetal inflammatory injury [77]. Preliminary results from our model indicate the potential effectiveness of this intervention (Fig. S5), suggesting that tolerance could be induced by reducing the inflammatory potential of Gram-negative commensals, similar to the effects observed with TLR4 antagonists. However, a complete exploration of such a scenario would necessitate expanding our model to explicitly incorporate cellular immunity, and to account for the distinct immunological effects of structurally diverse LPS molecules on early-life immune education [78] (Supplementary material, section *Model Limitations*).

Dysregulated immune development in children has repeatedly been shown to mediate multiple pathologies, ranging from developmental deficits in low-resource settings to immune-mediated disorders in developed countries [79]. Given the widespread burden of these conditions, developing a holistic mechanistic understanding of the concept of ‘immune education’ holds tremendous potential for clinical translation. Therefore, the primary motivation and contribution of this work is to establish the foundations of a mathematical framework for mechanistic thinking, generating clear and testable hypotheses for empirical investigation. Given the immense complexity and dynamism of early life, our model is necessarily a simplification, tailored to the range of data currently available, and excluding factors such as innate immune mechanisms, changes in gut permeability, and strain-specific shifts in the gut microbiome. Although quantitative predictions may change, qualitative conclusions are likely to be robust to these features(Supplementary material, section *Model Limitations*). While we included an illustrative validation using an external dataset (Supplementary Materials, section *Illustrative validation using an external dataset*), this serves primarily as a proof of concept. Future clinical, experimental, and measurement approaches are likely to open the way to evaluating our underlying mechanistic assumptions (Supplementary materials, section *Experimental Model Validation*). Ultimately, our quantitative analysis offers theoretical insights into pressing questions in the field, identifies the gaps in data collection needed to advance mechanistic understanding, and provides a systematic starting point for future data-driven mathematical models of early-life immune development.

## Acknowledgments

We are grateful to Prof. Dr. Emma Slack and Prof. Dr. M.D. Mathias Hornef for their valuable time and insightful discussions, which significantly improved our work.

## Authors contributions

Conceptualization: BT, AIL, CJEM

Data curation: BT

Methodology: BT, CJEM

Formal Analysis: BT, CJEM

Visualization: BT

Software: BT

Supervision: AIL, CJEM

Writing – original draft: BT, CJEM

Writing – review & editing: BT, AIL, CJEM

## Materials & Correspondence

Correspondence and material requests should be addressed to Burcu Tepekule.

## Data and materials availability

All data and code used in this work is available at https://github.com/burcutepekule/ONIMTOL under the GNU General Public License v3.0.

## Supplementary Materials

Materials and Methods

Supplementary Text

Figs. S1 to S11

Tables S1 to S6

References (80–180)

