## Supplementary Materials for "The ontogeny of immune tolerance: a model of early-life secretory IgA - gut microbiome interactions"

**The PDF file includes:**

Materials and Methods
Supplementary Text
Figs. S1 to S11
Tables S1 to S6
References (80–180)

### Materials and Methods

#### *Model Description*

Our model describes the reciprocal ontogeny of the gut microbiome and the gut mucosal secretory IgA (SIgA) response during the first two years of life, combining the ecology of the gut microbiome with the kinetics of the B cell responses in the gut-associated lymphoid tissues (GALT) and their bidirectional relationship. This period of early life is particularly dynamic, impacted by various external and internal factors such as feeding mode, microbial exposures, and the developmental clock of the host [80]. Given the varying influence of these factors at different stages, we categorize the developmental timeline into three distinct phases:

Maternal Phase: This phase represents the time frame from birth to transition to mixed feeding (MF), during which the infant is exclusively breastfed and thus the infant gut is primarily influenced by maternal factors. In breastfed infants, maternal secretory IgA (mSIgA) from breastmilk is active in the gut lumen, and nourishment primarily comes from human milk oligosaccharides (HMOs). Key immune structures like germinal centers (GCs) and Microfold (M) cells in the GALT inductive sites (Peyer's Patches (PPs) in the small intestine and the colonic patches in the colon) are not fully developed until decreasing concentrations of the maternal factors in the breastmilk favor M cell maturation, delaying the onset of an endogenous secretory IgA (eSIgA) response [35]. Naïve B cells migrate to gut lymphoid tissues but remain inactive until M-cell mediated antigenic sampling begins [81].

Developmental Phase: This phase encompasses the transition from exclusive breastfeeding (EBF) to MF and continues until the microbiome composition and eSIgA response stabilize, including the shift to exclusive solid feeding (ESF). As the influence of maternal antibodies diminishes, the infant's diet increasingly includes plant-derived polysaccharides (PDPs) instead of HMOs, meeting the total caloric need of the infant. Introduction of PDPs increases the overall taxonomic diversity of the gut microbiome by introducing novel antigens. Concurrently, GCs and M cells mature, fostering the development of the eSIgA response, a phenomenon often referred to as the "weaning reaction" [10]. Naïve immune cells become activated, participate in germinal center reactions, and eventually produce plasma cells secreting IgA.

Steady Phase: This is the final phase where the host achieves a stable, complex microbiome and a fully matured endogenous adaptive immune system which, crucially, resembles their mother's SIgA response [21]. The average affinity of the plasma cell pool and the relative abundance of key microbial populations reaches a steady state, indicative of a homeostatic gut ecosystem. During this phase, the host's diet is based solely on PDPs.

While the same equations apply across all phases, separating them into maternal, developmental, and the steady phases is crucial for parameter inference, which will be elucidated in the subsequent sections.

Our mathematical model consists of a system of ordinary and delay differential equations tracking bacterial abundances in the gut lumen and the fecal samples of the host, as well as the immune response focusing on antigen-specific SIgA to gut commensals and their affinity maturation process. A compartmental diagram illustrating the dynamics of the model is given in Fig. 1 (see introduction), model parameters with their corresponding notations, descriptions, units, and numerical ranges are given in Table S3, and the set of differential equations governing our system are provided via Eqns. S1.2.1-S1.2.14. Additional assumptions implicit to the model structure are given in Table S4.

***Formalizing secretory IgA (SIgA) action on microbial taxa***

Our model capitalizes on SIgA's multifunctionality within the gut lumen, categorizing its actions based on its affinity level [48]. For consistency, we denote both the affinity of the antigen-binding site of a B-cell receptor (BCR) on the B cells and the affinity of the SIgA produced by the plasma cells by $\rho_i$, where $i$ represents the taxon index for $1 \leq i \leq N$, and $N$ represents the number of taxonomic groups. SIgA may exert non-neutralizing but masking (M) actions, reflecting low affinity antibodies, facilitating immune exclusion (preventing commensal bacteria from interacting with the immune system) and enhancing adherence to the gut lumen; and neutralizing (N) actions, reflecting high affinity antibodies, contributing to the expulsion of bacteria from the gut lumen [17]. The activities of "masking" and "killing" are not mutually exclusive (Fig. 2B), with their rates determined by the affinity levels:

$$\mu^{m}{}_{i} = \frac{1}{1+\rho_i^{-p}}, \textbf{(S1.1.1)}$$

$$\mu^{n}{}_{i} = 1 - \frac{1}{1+\rho_i^{-p}}, \textbf{(S1.1.2)}$$

where, $\mu^{m}{}_{i}$ and $\mu_i{}^{n}$ represent the masking and neutralizing rates of SIgA with affinity $\rho_i$, respectively. To differentiate the impact of maternal SIgA (mSIgA) from endogenous SIgA (eSIgA), we use 4 different parameters - $\mu_i{}^{m,n}$, $\mu_i{}^{m,m}$, $\mu_i{}^{e,n}$ and $\mu_i{}^{e,m}$ - describing the neutralizing and masking rates of mSIgA and eSIgA for each taxon $i$.

Similar to their rate of action, the binding ability $\omega_i$ of the antibodies is determined by their respective affinity levels,

$$\omega_i = 1 - \frac{r_d}{1+\rho_i}, \textbf{(S1.1.3)}$$

where $r_d$ denotes the dissociation rate of SIgA, describing the rate at which the antibody dissociates from its target [82]. Masking and neutralizing rates are scaled by the binding ability for each taxon, which modulates their overall effectiveness. This dual function of SIgA creates three subpopulations for each microbial taxon $i$: uncoated (${y_i}^{L,uc}$), masked (${y_i}^{L,m}$), and neutralized, where the total population size that is metabolically active in the gut lumen is ${y_i}^{L} = {y_i}^{L,uc} + {y_i}^{L,m}$.

The exchange rate between ${y_i}^{L,m}$ and ${y_i}^{L,uc}$ depends on the level of mSIgA and eSIgA in the gut lumen, the half-life of SIgA ($I_{hl}$), the binding ability ($\omega_i$), and masking and killing rates (${\mu_i}^{m,m}$, and ${\mu_i}^{e,m}$ ) (Eqns. S1.2.5-S1.2.6). Throughout the remainder of this text, SIgA-, SIgA+ (M), SIgA+ (N) will denote bacteria and/or antigens that are free of SIgA, forming complexes masked with SIgA, and neutralized by SIgA, respectively. Both masked and neutralized populations are coated with SIgA, meaning that techniques like IgA-Seq or BugFACS would detect SIgA binding in both cases. The SIgA coating ratio, which measures the proportion of bacteria coated with SIgA, would reflect contributions from both masked and neutralized bacteria.

***Characterizing the ecological interactions within the microbial community***

The absolute abundance of each taxon (${y_i}^{L}$ ) is governed by a generalized Lotka–Volterra (gLV) model [83], incorporating a net growth rate $\lambda_i$ and an interaction matrix $\beta$, where $\beta_{i,j}$ denotes the impact of the abundance of taxon $j$ on the abundance of taxon $i$. We assume that ${y_i}^{L,uc}$ and ${y_i}^{L,m}$ interact identically with their counterparts, therefore the net change for each taxon is calculated based on the total abundance ${y_i}^{L}$ (Eqn. S1.2.4), which is then divided into two subpopulations ${y_i}^{L,uc}$ and ${y_i}^{L,m}$ depending on the processes described above. While the GLV model simplifies higher-order dynamics by considering only pairwise interactions, this simplification is a well-established method that balances model complexity with the feasibility of parameter inference and has been successfully employed in several quantitative studies of gut microbiota dynamics [84–86].

In addition to gLV dynamics, factors in the gut lumen such as the oxygen concentration ($O_2$), caloric intake from HMOs, and caloric intake from PDPs directly modulate the growth rate of bacteria depending on their capabilities to proliferate in aerobic conditions (${\phi_i}^{O2}$) and their efficiency in metabolizing different energy sources (${\phi_i}^{HMOs}, {\phi_i}^{PDPs}$ ) as follows,

$$\lambda_i^{adj}(t) = \left(1 + O_2(t){\phi_i}^{O_2} + HMOs(t){\phi_i}^{HMOs} + PDPs(t){\phi_i}^{PDPs}\right)\lambda_i, \textbf{(S1.1.4)}$$

where $\lambda_i^{adj}$ denotes the adjusted growth rate for taxon $i$. As a result of these dynamics, $O_2$ - which is assumed to be present in the gut lumen at its maximum value after birth - is depleted at a constant rate ($\mu_{O2}$) by facultative anaerobes (${y_{fn}}^{L}$) over time (Eqn. S1.2.3). Caloric intake from HMOs and PDPs as well as the mSIgA levels depend on the Exclusive Breastfeeding (EBF) and Mixed

Feeding (MF) durations, and the details of their parameterization will be discussed in the following sections.

#### *Modeling B-cell ontogeny*

To characterize the dynamics underlying 'immune education', our model focuses on the ontogeny of the B-cell response of the infant, while implicitly integrating the role of the T cell arm of humoral immunity. This requires formalizing an affinity maturation and plasma cell differentiation process that reflects BCR affinity on a continuous scale. To this end, we developed a set of differential equations that track the average affinity of the circulating B cells within the GCs and plasma cells in the Lamina Propria (LP). Our approach, inspired by previous work such as Molari *et al.* [87], adapts and extends their quantitative techniques to the specific context of gut mucosal immunity, incorporating continuous BCR affinity distributions and multiple thresholds for B cell fate decisions. Although previous stochastic models have explored germinal center dynamics [88–93], they primarily focus on vaccine development, systemic immunity, or model antibody affinity in a discretized manner, which limits their direct integration to our framework.

Circulating B cells originate from the naïve B cells migrating from bone marrow to the PPs, where they get activated upon antigen encounter. This activation rate depends on the linear combination of IgA-antigen immune complexes (${z_i}^m$) and IgA-free antigens (${z_i}^{uc}$) transported across M cells [33] on the PPs,

$$\psi_i = \psi^m {z_i}^m + \psi^{uc} {z_i}^{uc} \text{ , } \mathbf{(S1.1.5)}$$

where the activation rates per sampled antigen for ${z_i}^m$ and ${z_i}^{uc}$ are denoted by $\psi^m$ and $\psi^{uc}$ , respectively. $\psi_i$ represents the total activation rate dedicated to taxon $i$, and we assume that $\psi^{uc} > \psi^m$ due to the enhanced anti-inflammatory effects of ${z_i}^m$ compared to ${z_i}^{uc}$ [94,95] and thus the promotion of increased B cell recruitment and activation per ${z_i}^{uc}$ [96].

Five factors influence antigenic sampling by M-cells: *i)* the developmental clock of the host, and associated functionality of GALT inductive sites, GCs, and M cells (which gradually increase over ontogeny) [14]; ii*)* factors in the breastmilk that suppress the functionality of GALT inductive sites, GCs, and M cells [35]; *iii)* microenvironmental stimulation ($\sigma$) representing the anti- or pro-inflammatory tone of the microenvironment, influencing barrier integrity [97] and M cell function [98]; *iv)* the sampling rates of SIgA- and SIgA+ (M) bacterial antigens; and v) invasiveness of each taxon ($\alpha_i$), which measures the bacteria's capacity to adhere to and penetrate intestinal epithelial cells [99]. After the host reaches the age at which their endogenous system becomes functional, ${z_i}^{uc}$ and ${z_i}^m$ can be expressed as,

$$ {z_i}^{uc} = \sigma\, \alpha_i \epsilon^{uc} {y_i}^{L,uc} \text{ , } \mathbf{(S1.1.6)}$$
$$ {z_i}^{m} = \epsilon^{m} {y_i}^{L,m} \text{, } \mathbf{(S1.1.7)}$$

where $\epsilon^{uc}$ and $\epsilon^m$ denote the sampling rates of ${y_i}^{L,uc}$ and of ${y_i}^{L,m}$ for any taxon $i$, respectively. Given the M cells' binding capacity to SIgA [18,100,101], we assume that $\epsilon^m > \epsilon^{uc}$. Considering the role of SIgA binding in encouraging biofilm formation and thereby enhancing commensals' adherence to the mucosal surface in a beneficial manner [102–104], we assume that the sampling of $y_i^{L,uc}$ is influenced by both $\sigma$ and $\alpha_i$; whereas for $y_i^{L,m}$ , sampling occurs more consistently, determined solely by the sampling rate $\epsilon^m$.

***Role of the microenvironment***

Microenvironment in the gut lumen and within the GALT inductive sites has a significant role in modulating the antigenic sampling and the affinity maturation process, respectively. Here, we define microenvironmental stimulation ($\sigma$) as a composite variable that encapsulates the cumulative effects of cytokines, microbial products, and host-derived signals on the local tissue milieu. The intestinal epithelial cells (IECs) that line the gut lumen possess a variety of receptors, including pattern recognition receptors (PRRs) like Toll-like receptors (TLRs) and NOD-like receptors (NLRs), which can detect microbial-associated molecular patterns (MAMPs) from commensal or pathogenic microbes [105,106]. Upon sensing these signals, IECs produce and release a range of signaling molecules, including cytokines and chemokines that modulate immune responses [107]. Considering this signal transmission pathway, we define microenvironmental stimulation as,

$$\sigma = \sum_i \kappa_i y_i^{L,uc} - \sum_i I_i y_i^{L}, \quad \textbf{(S1.1.8)}$$

where $\kappa_i$ represents the relative immunostimulatory capacity of the SIgA- members of taxon $i$ compared to their SIgA+ (C) counterparts. $I_i$ is a binary variable representing the inherent anti-inflammatory potential of taxon $i$, including mechanisms such as short-chain fatty acid production, modulation of immune cell activity, or enhancement of epithelial barrier function [108–111]. Assigning a binary value to $I_i$ and continuous value to $\kappa_i$ allows us to focus solely on quantifying the *relative* immunostimulatory capacity of unmasked commensals [40,41], simplifying the parameter inference procedure discussed in the subsequent sections. Our framing captures how commensals can either reduce inflammation or, conversely, display pathogenic-like behavior when gut homeostasis is disrupted and the regulatory effect of SIgA masking on bacterial populations is compromised. For commensals without any anti-inflammatory capacity, $I_i$ would be zero, meaning that their masked counterparts are irrelevant in modulating the microenvironment. Thus, in the extremes, exclusively probiotic bacteria would be assigned $I_i$ = 1 and $\kappa_i$ = 1, meaning that their unmasked counterparts do not cause any increase in the inflammatory tone, whereas their masked counterparts decrease inflammation proportional to their abundance in the gut lumen ($\sigma = \sum_i \kappa_i y_i^{L,uc} - \sum_i I_i y_i^{L} = \sum_i (\kappa_i - I_i) y_i^{L,uc} - \sum_i I_i y_i^{L,m} = -\sum_i y_i^{L,m} \mid \kappa_i = I_i = 1$ ). In contrast, exclusively pathogenic bacteria would have $I_i$ = 0 and a high $\kappa_i$ underscoring their substantial capacity to induce inflammation. To reduce the impact of identifiability problems during inference, we fixed $\kappa_i = 1$ for all the symbiotic commensals (*Bifidobacteriaceae*, *Bacteroidaceae* and *Clostridiales*) in our model, and only inferred $\kappa_E$ for *Enterobacteriaceae*.

Optimal regulation of $\sigma$ is essential for maintaining gut homeostasis; however, when excessive, it can compromise the integrity of the intestinal barrier by disrupting tight junctions between epithelial cells, leading to altered M cell function either by direct damage or through influencing the M cell differentiation and function [98]. Therefore, $\sigma$ is used as an effect modulator in Eqn. S1.1.6.

Elevated levels of stimulation in the lumen can trigger IECs to produce pro-inflammatory signals directed toward essential immune cells involved in B cell maturation, including dendritic cells

(DCs) and T cells located in the subepithelial dome (SED) of the GALT inductive sites. While recognizing that the lumen and the distinct zones within the GALT inductive sites exhibit specific microenvironmental characteristics, we assume that the microenvironmental tone in the lumen is mediated through the epithelium [42,112] and influences the tone in the GALT inductive sites in a similar fashion. Thus, we employ $\sigma$ to simultaneously describe the microenvironmental stimulation in the gut lumen and within the GALT inductive sites.

***Modeling the role of Germinal Centres***

The microenvironment within the GALT inductive sites plays a crucial role in steering the balance between T follicular helper (Tfh) and follicular regulatory (Tfr) cells [113,114]. This balance is particularly important to select the pool of circulating B cells with the appropriate BCR affinity distribution to continue through each proliferation, somatic hypermutation, and selection (P-SHM-S) cycle. However, this conventional view of affinity maturation does not allow for antigen-specific low-affinity clones to survive the competition for T cell help in the light zone. An alternative view suggests that the continually available antigen and sufficient T-cell help (possibly of a range of types) cooperating with commensal-derived signals acting via pattern recognition receptors (PRRs) supports the survival and differentiation of low-affinity cells as well as the high affinity ones [115]. Therefore, assuming that the circulating B cells constitute a normally distributed set of BCR affinities [87], we propose a variable reflecting the Tfh:Tfr ratio which determines the BCR affinity *range* that will receive adequate T cell help to continue with the next SHM cycle, and eventually the average affinity of the plasma cells leaving the GC. We refer to this quantity as the selection threshold, denoted by $\delta_i$.

In addition to the microenvironment, antigen presentation is pivotal in determining the Tfr versus Tfh differentiation via conditioning of mucosal dendritic cells [116,117]. SIgA-coated bacterial antigens promote the tolerogenic programming of dendritic cells via the C-type lectin receptor SIGNR1 [75]. Given that the sampling of SIgA-antigen immune complexes will skew the bias in differentiation toward tolerogenic profiles compared to SIgA-free antigens [94,118], we model the rate of change in $\delta_i$ as a function of the ratio of cumulative uncoated $({z_i}^{\Sigma uc}(t) = \sum_t \quad {z_i}^{uc}(t))$ to cumulative total sampled antigens $({z_i}^{\Sigma}(t) = \sum_t \quad ({z_i}^{uc}(t) + {z_i}^{c}(t)))$, as well as the microenvironmental stimulation $\sigma$ (Eqn. S1.2.7), as a proxy to reflect cumulative imprinting of the Tfh:Tfr ratio. This formulation allows us to implicitly model the impact of T cells on GC reactions, although we do not explicitly include them in the differential equation system.

We model one cycle of GC reactions as follows,

1. At each timestep $t$, a population of naïve B cells $({B_i}^n)$ in the number of $\Delta {B_i}^n$ migrate from bone marrow to the dark zone PPs. As for T cells, the rate of migration is modeled as an exponentially decreasing function of time $t$ $(C_n e^{-c_n t})$, reflecting the diminishing pool of naïve cells as the host ages (Eqn. S1.2.8) [14,119]. Although naïve cells are not antigen-specific, for the sake of completeness, we use the taxon subscript $i$ to track the number of B cells activated in an antigen-specific fashion. These cells lay dormant in the GALT inductive sites until the M cells and GCs mature at time $t^m$.

2. At time $t^m$, $B_i^{\,n}$ starts getting activated at the rate of $\psi_i$(Eqn. S1.1.5) and join the pool of circulating B cells ($B_i^{\,c}$) participating in the GC reactions. These newly activated cells are calculated as $B_i^{\,c,new} = \psi_i B^n{}_i$.

3. Newly activated population of circulating B cells have a normal BCR affinity distribution with mean 0 and a standard deviation proportional to their population size, providing enough BCR variability for the selection process in the light zone.

4. Given the value of $\delta_i$, selection pressure imposed by T cells segregates the circulating cells in the light zone into 4 different groups,

    4.1. Cells with BCR affinity < $th_{apop}\delta_i$ fail to compete for T cell help and die through apoptosis. The number of these cells is equal to $B_i^{\,c}\Phi((th_{apop}\delta_i - \bar{\rho}_i^{\,c})/\sigma_i^{\,c})$, where $\Phi$, $\bar{\rho}_i^{\,c}$, and $\sigma_i^{\,c}$ denote the cumulative distribution function for normal distribution, the mean, and the standard deviation of the $B_i^{\,c}$ BCR affinity distribution, respectively.

    4.2. Cells with BCR affinity in the range of $[th_{apop}\delta_i,\ th_{high}\delta_i]$ will receive enough T cell help to continue circulating and will move to the dark zone for the next round of proliferation and SHM cycle in the dark zone. The number of these cells is equal to $B_i^{\,c}(\Phi((th_{high}\delta_i - \bar{\rho}_i^{\,c})/\sigma_i^{\,c}) - \Phi((th_{apop}\delta_i - \bar{\rho}_i^{\,c})/\sigma_i^{\,c}))$.

    4.3. Cells with BCR in the range of $[th_{high}\delta_i,\ th_{ang}\delta_i]$ will receive enough T cell help to terminally differentiate to plasma cells with the average affinity of $0.5(th_{ang} + th_{high})\delta_i$. The number of these cells is equal to $B_i^{\,c}(\Phi((th_{ang}\delta_i - \bar{\rho}_i^{\,c})/\sigma_i^{\,c}) - \Phi((th_{high}\delta_i - \bar{\rho}_i^{\,c})/\sigma_i^{\,c}))$.

    4.4. Cells with BCR affinity above $th_{ang}\delta_i$ will be rendered anergic through additional regulatory mechanisms, mirroring the principles of negative selection that eliminate self-reactive B cells [120]. The number of these cells is equal to $1 - B_i^{\,c}\Phi((th_{ang}\delta_i - \bar{\rho}_i^{\,c})/\sigma_i^{\,c})$.

This selection process confines the BCR affinity range of circulating cells to $[th_{apop}\delta_i,\ th_{high}\delta_i]$ prior to the next SHM cycle. Although the process involves truncating a Normal distribution, for the sake of analytical computability, we assume that the resulting population maintains a normal distribution with mean $0.5(th_{apop}\delta_i + th_{high}\delta_i)$ and standard deviation $(1/6)(th_{high}\delta_i - th_{apop}\delta_i)$, indicating that 99.7% of the affinity levels lie within the $[th_{apop}\delta_i,\ th_{high}\delta_i]$ range.

5. For $t > t^m$, circulating cells that received enough T cell help at the previous time step $(t - \Delta t)$ will migrate back to the dark zone, proliferate with rate $r_p$, and go through another round of SHM. We assume that the proliferation phase only changes the size of the $B_i^{\,c}$ pool, whereas SHM increases variability in the circulating BCR affinity pool. Each round of SHM increases the standard deviation of the circulating BCR affinity distribution proportional to $\tau^c|\delta_i(t+1) - \bar{\rho}_i^{\,c}(t)|$ to ensure that there is enough variation

in the circulating BCR affinities, allowing the model to mimic the ranking-based selection imposed by the T cells in the light zone of the GCs [88].

6. Proliferated and mutated cells will be pooled together with the newly activated cells ${B_i}^{c,new}$, which will modify the pool size as well as the mean and the standard deviation of the BCR affinity distribution. Since our model is based on differential equations, we keep track of the difference of the circulating pool size, the mean, and the standard deviation of the circulating BCR affinity pool between consecutive timesteps. We approximate the pooled mean and standard deviation as a weighted sum by the sample sizes of the already circulating and newly added cells, and calculate the differences as,

$$\Delta {B_i}^c(t+\Delta t) = {B_i}^c(t)\left[\Phi\left(\frac{th_{high}\delta_i(t)-\bar{\rho_i}^c(t)}{{\sigma_i}^c(t)}\right) - \Phi\left(\frac{th_{apop}\delta_i(t)-\bar{\rho_i}^c(t)}{{\sigma_i}^c(t)}\right)\right](1+r_p) + {B_i}^{c,new}(t) - {B_i}^c(t), \quad \textbf{(S1.1.9)}$$

$$\Delta \bar{\rho_i}^c(t)(t+\Delta t) = \frac{{B_i}^c(t)\left[\Phi\left(\frac{th_{high}\delta_i(t)-\bar{\rho_i}^c(t)}{{\sigma_i}^c(t)}\right)-\Phi\left(\frac{th_{apop}\delta_i(t)-\bar{\rho_i}^c(t)}{{\sigma_i}^c(t)}\right)\right]\int_{th_{apop}}^{th_{high}} \mathcal{N}\left(\bar{\rho_i}^c(t),{\sigma_i}^c(t)\right)\partial\rho}{{B_i}^c(t)\left[\Phi\left(\frac{th_{high}\delta_i(t)-\bar{\rho_i}^c(t)}{{\sigma_i}^c(t)}\right)-\Phi\left(\frac{th_{apop}\delta_i(t)-\bar{\rho_i}^c(t)}{{\sigma_i}^c(t)}\right)\right](1+r_p)+{B_i}^{c,new}(t)} - \bar{\rho_i}^c(t), \quad \textbf{(S1.1.10)}$$

$$\Delta {\sigma_i}^c(t+\Delta t) = \frac{{B_i}^c(t)\left[\Phi\left(\frac{th_{high}\delta_i(t)-\bar{\rho_i}^c(t)}{{\sigma_i}^c(t)}\right)-\Phi\left(\frac{th_{apop}\delta_i(t)-\bar{\rho_i}^c(t)}{{\sigma_i}^c(t)}\right)\right](1+r_p)\left(\frac{1}{6}\right)\left[th_{high}\delta_i - th_{apop}\delta_i\right] + {B_i}^{c,new}(t){\sigma_i}^{new}}{{B_i}^c(t)\left[\Phi\left(\frac{th_{high}\delta_i(t)-\bar{\rho_i}^c(t)}{{\sigma_i}^c(t)}\right)-\Phi\left(\frac{th_{apop}\delta_i(t)-\bar{\rho_i}^c(t)}{{\sigma_i}^c(t)}\right)\right](1+r_p)+{B_i}^{c,new}(t)} - {\sigma_i}^c(t), \quad \textbf{(S1.1.11)}$$

where $\mathcal{N}(x \mid \mu, \sigma)$ denotes the Normal probability distribution function with mean $\mu$ and standard deviation $\sigma$, and ${\sigma_i}^{new} = \tau^{new}\, {B_i}^{c,new}$ as discussed in step 3 above.

7. The same rationale applies when calculating the difference of the size of the plasma cell pool (${B_i}^p$) and the mean of the plasma BCR affinity pool ($\bar{\rho_i}^p$) between consecutive timesteps

$$\Delta {B_i}^p(t+\Delta t) = {B_i}^p(t)\left[\Phi\left(\frac{th_{ang}\delta_i(t)-\bar{\rho_i}^c(t)}{{\sigma_i}^c(t)}\right) - \Phi\left(\frac{th_{high}\delta_i(t)-\bar{\rho_i}^c(t)}{{\sigma_i}^c(t)}\right)\right](1 - {B_i}^p(t)), \quad \textbf{(S1.1.12)}$$

$$\Delta \bar{\rho_i}^p(t+\Delta t) = \frac{{B_i}^p(t)\left[\Phi\left(\frac{th_{ang}\delta_i(t)-\bar{\rho_i}^c(t)}{{\sigma_i}^c(t)}\right)-\Phi\left(\frac{th_{high}\delta_i(t)-\bar{\rho_i}^c(t)}{{\sigma_i}^c(t)}\right)\right](0.5(th_{ang}+th_{high})\delta_i(t))+{B_i}^p(t)\,\bar{\rho_i}^p(t)}{{B_i}^p(t\left[\Phi\left(\frac{th_{ang}\delta_i(t)-\bar{\rho_i}^c(t)}{{\sigma_i}^c(t)}\right)-\Phi\left(\frac{th_{high}\delta_i(t)-\bar{\rho_i}^c(t)}{{\sigma_i}^c(t)}\right)\right]{B_i}^p(t)} - \bar{\rho_i}^p(t). \quad \textbf{(S1.1.13)}$$

Note that $\Delta {B_i}^p(t+\Delta t)$ is multiplied by $(1 - {B_i}^p(t))$ to ensure that the plasma cell pool saturates at the carrying capacity of 1, reflecting the carrying capacity for plasma cells in the LP [121].

We assume that the longevity of plasma cells is primarily determined by the cumulative T cell help they have received at the time they left the GC reaction [122]. Since the affinity levels (or magnitude of $\delta_i$) do not necessarily equate to the cumulative T cell help (or the number of SHM-S-P cycles) itself, we employ delayed differential equations to track the relative progression of $\delta_i(t)$, and define the death rate of plasma cells as ${\mu^p}_i = \frac{\delta_i(t-1)}{\delta_i(t)}$ . By doing so, we introduce a higher death rate to the plasma cells that were produced at the early stages of immune ontogeny, and as the host matures and $\delta_i(t)$ approaches $\lim_{t\to\infty} \delta_i(t)$, ${\mu^p}_i$ approaches to 0.

In reality, plasma cells do not live infinitely long, therefore a ${\mu^p}_i$ of 0 would not be correct. However, plasma cells are continuously replaced by the memory cells, which we do not include in our model (see section *Model Limitations*). Therefore, ${\mu^p}_i$= 0 at the Steady Phase mimics the continuous replacement of plasma cells by the memory cells and thus the plasma cell population is kept at a steady level.

***Linking model outputs to data on microbial taxa***

Our model quantifies the absolute abundances of microbial populations within the gut lumen. However, for parameter inference, we relied on fecal samples from infants, analyzed using 16S rRNA sequencing techniques [20–22]. Fecal samples serve as a proxy for assessing the taxonomic distribution within the gut lumen. However, equating these two environments directly overlooks the distinct roles of SIgA—specifically, its ability to either neutralize bacteria by facilitating their expulsion from the gut lumen [43] or, conversely, enhancing their adherence through masking without neutralization [46,47]. To incorporate this dual functionality, we introduce 3 different observation rates $s^n, s^m,$ and $s^{uc}$ for neutralized, masked, and uncoated subpopulations in the fecal samples, assuming that the observed quantity is the sum of these subpopulations weighed by their respective observation rates. We keep track of the cumulative neutralized bacteria abundance (${dy_i}^{L,\Sigma n}$) to quantify the total bacterial population in the fecal samples, although they are assumed to remain inactive, neither interacting with other bacteria nor stimulating the immune system. During parameter inference, we reconstruct the absolute abundances in fecal samples as follows,

$$ {w^n}_i = s^n \frac{d{y_i}^{L,\Sigma n}}{dt}, \quad \textbf{(S1.1.14)} $$
$$ {w^m}_i = s^m {y_i}^{L,m}, \quad \textbf{(S1.1.15)} $$
$$ {w^{uc}}_i = s^{uc} {y_i}^{L,uc}, \quad \textbf{(S1.1.16)} $$
$$ w_i = {w^n}_i + {w^m}_i + {w^{uc}}_i, \quad \textbf{(S1.1.17)} $$

where ${w^n}_i$, ${w^m}_i$, ${w^{uc}}_i$, and $w_i$ denote the neutralized, masked, uncoated, and total absolute abundances observed per fecal content for taxon $i$, respectively.

***Informing time varying components of the model***

Our model mainly focuses on the mechanisms governing the GC reactions in the GALT inductive sites, and the main route of antigen transport to these sites is carried out by M cells which appear shortly before weaning primarily due to the decreasing concentrations of maternal steroids in the breastmilk [33,35]. In the absence of quantitative data, we propose using the concentration of mSIgA as an indirect measure of maternal steroid levels. We assume that antigenic sampling beings as $t^{50}$, representing the time point when mSIgA concentration falls below 50% of its peak value, favoring the qualitative observation regarding the close timing of M cell maturation to weaning (Fig. 1A). In the absence of breastfeeding, various factors—including the maturation of dendritic cells (DCs) and B cell receptors—may constrain the endogenous immune response, notably affecting the development of IgA-secreting plasma cells [123–125]. Observations in the literature suggest that these constraints are particularly pronounced during the first month of human life, and infants who do not receive human milk compensate by starting to produce SIgA in the intestine around 4 weeks of age [44]. Therefore, we assume that the endogenous immune system activation occurs at day $t^{m} = (t^{50}, 30)$, allowing us to compute $t^{m}$ for a generic combination of EBF and MF durations.

To simulate our system for any given feeding pattern, we parameterize mSIgA, HMOs, and PDPs as functions of EBF and MF durations denoted by $T_{EBF}$ and $T_{MF}$, respectively. To do so, we used the estimated calorie requirements of infants [25–27], and assumed that *i)* average calories in breastmilk per volume is constant through the period of breastfeeding and the caloric needs are met by modulating the volume ingested by the infant, *ii)* total calorie requirement of the infant for a given time point is always met regardless of the feeding pattern, and *iii)* when the infant starts MF, HMOs are reduced and PDPs are introduced gradually (Fig. 2A). Upon switching to MF at time point $t_{MF}$, this pattern resumes a gradual increase with a notable rise in the proportion of calories derived from PDPs. We formulate the function for HMOs caloric intake as,

$$HMOs(t) \stackrel{\text{def}}{=} f_{HMOs}(t, t_{MF}, T_{MF}) = \{K_{H1} + V_0(1 - e^{-c_H t}), K_{H2}(1 - e^{-c_H t_{mirror}}),\ t < t_{MF} \quad t \geq t_{MF}\ , \textbf{(S1.1.18)}$$

where $t_{mirror} = \frac{t_{MF}[T_{MF} - (t - t_{MF})]}{T_{MF}}$ represents the adjusted time points between $t_{MF}$ and the switch to ESF. $K_{H2} = \frac{K_{H1} + V_0(1 - e^{-c_H t_{MF}})}{1 - e^{-c_H t_{MF}}}$, ensuring a smooth transition at $t = t_{MF}$. To calculate the from PDPs, we first formulate the function for the total caloric intake $f_{TOT}$ as,

$$f_{TOT}(t, t^{*}_{MF}, T^{*}_{MF}) = \{f_{HMOs}(t, t^{*}_{MF}, T^{*}_{MF}), \frac{K_C}{1 + e^{-c_C(t - t^{*}_{MF})}}, \frac{K_C}{1 + e^{-c_C(t_C - t^{*}_{MF})}}, t < t^{*}_{MF} \quad t^{*}_{MF} \leq t < t_C \quad t \geq t_C\ , \textbf{(S1.1.19)}$$

where $t^{*}_{MF}$ and $T^{*}_{MF}$ denote the $t_{MF}$ and $T_{MF}$ values derived from the study that provided data for our model parameterization [18]. $K_c$ denotes the fixed caloric intake of the host when they reach the steady phase at $t = t_c$, and $c_c$ adjusts the rate of increase that intake until $t = t_c$. Eqn. S1.1.19 ensures that the total caloric intake of the infant for any given $\{t_{MF}, T_{MF}\}$ combination matches the total caloric intake in the study used for model parameterization. Given Eqn. S1.1.18 and S1.1.19, we can compute the caloric intake from PDPs as,

$$PDPs(t) \stackrel{\text{def}}{=} f_{PDPs}(t, t_{MF}, T_{MF}) = \{0, f_{TOT}(t, t^{*}_{MF}, T^{*}_{MF}) - f_{HMOs}(t, t_{MF}, T_{MF}),\ t < t_{MF} \quad t \geq t_{MF}\ , \textbf{(S1.1.20)}$$

Data in literature indicate that SIgA levels in breastmilk peak in the days immediately postpartum, followed by a decline over the first month, eventually stabilizing into a lower, steady state in mature milk [113–115]. Consequently, we formulate and parameterize the rate of change in mSIgA levels in the breast milk as,

$$\Delta f_{mSIgA}(t) = K_I + m_I e^{-c_I t}, \textbf{(S1.1.21)}$$

where $K_I$ represents the steady-state influx rate of mSIgA into the gut lumen as found in mature milk. Parameters $m_I$ and $c_I$ modulate the mSIgA level shortly after birth and its subsequent decline. This rate is modulated by $HMOs(t)$ as it is used as a proxy for the volume of breastmilk ingested by the infant (Eqn. S1.2.1).

***Differential Equation System***

*Microenvironmental Dynamics*

$$\frac{dmSIgA}{dt} = +HMOs(t)\Delta f_{mSIgA}(t) - I_{hl} mSIgA\ . \textbf{(S1.2.1)}$$

$$\frac{deSIgA}{dt} = +C_I B_i^{\,p}(1 - (1 - I_{hl}/C_I) eSIgA) - I_{hl} eSIgA\ . \textbf{(S1.2.2)}$$

$\frac{dO_2}{dt} = -\mu_{O2} O_2\, y_{fn}{}^L$ . **(S1.2.3)**

*Community Dynamics*

$\frac{dy_i{}^L}{dt} = y_i{}^L(\lambda_i{}^{adj} + \beta_i) - (\omega_i{}^{m,n}\mu_i{}^{m,n} + \omega_i{}^{e,n}\mu_i{}^{e,n})y_i{}^{L,uc}$ **(S1.2.4),** where $\beta_i = \sum_{j=1}^{N} \beta_{i,j}\, y_j{}^L$.

$\frac{dy_i{}^{L,uc}}{dt} = \frac{dy_i{}^L}{dt} + (1/I_{hl})y_i{}^{L,c} - (\omega_i{}^{m,c}\mu_i{}^{m,c} + \omega_i{}^{e,c}\mu_i{}^{e,c})y_i{}^{L,uc}$ . **(S1.2.5)**

$\frac{dy_i{}^{L,c}}{dt} = -(1/I_{hl})y_i{}^{L,c} + (\omega_i{}^{m,c}\mu_i{}^{m,c} + \omega_i{}^{e,c}\mu_i{}^{e,c})y_i{}^{L,uc}$ . **(S1.2.6)**

$\frac{dy_i{}^{L,\Sigma n}}{dt} = +(\omega_i{}^{m,n}\mu_i{}^{m,n} + \omega_i{}^{e,n}\mu_i{}^{e,n})y_i{}^{L,uc}$. **(S1.2.7)**

*Immune Dynamics*

To make our model neutral to the contribution of $\sigma$ vs $\frac{z_i{}^{\Sigma uc}(t)}{z_i{}^{\Sigma}(t)}$ in Eqn. S1.2.8 below, we introduce $\tau^\sigma$ as a multiplicative factor such that $max_t\, \tau^\sigma\sigma(t) = 1$. Due to the dynamic nature of our model, analytically calculating $\max_t \sigma(t)$ is not possible. However, the upper bound of this quantity can be calculated by using Eqns. S1.1.8 and S1.2.5 by assuming all bacteria in the lumen are uncoated and have reached their carrying capacity in the absence of any neutralization by antibodies. However, this calculation would require a numerical integration for a multi-taxa system. We can further approximate the upper bound by only using the steady state value of the pathogenic taxon in our model, which has the highest $\kappa_i$ that is orders of magnitude larger than the $\kappa_i$ for symbiotic commensals, and approximate $\tau^\sigma = 1/(\kappa_i\lambda_i/\beta_{ii})$.

$\frac{d\delta_i}{dt} = +\tau^\delta\, C_T e^{-c_T t}\, log\,(1 + \tau^\sigma\sigma)\, log\,(1 + \frac{z_i{}^{\Sigma uc}(t)}{z_i{}^{\Sigma}(t)})$. **(S1.2.8)**

$\frac{dB_i^n}{dt} = +C_B e^{-c_B t} - \psi_i B_i^n$. **(S1.2.9)**

$\frac{dB_i{}^c}{dt} = +\Delta B_i{}^c(t + \Delta t)$. **(S1.2.10)**

$\frac{dB_i{}^p}{dt} = +\Delta B_i{}^p(t + \Delta t) - \mu^p{}_i B_i{}^p$. **(S1.2.11)**

$\frac{d\bar{\rho}_i{}^c}{dt} = +\Delta\, \bar{\rho}_i{}^c(t + \Delta t)$. **(S1.2.12)**

$\frac{d\bar{\rho}_i{}^p}{dt} = +\Delta\, \bar{\rho}_i{}^p(t + \Delta t)$. **(S1.2.13)**

$\frac{d\sigma_i{}^c}{dt} = +\Delta\sigma_i{}^c(t + \Delta t)$. **(S1.2.14)**

Given the complexity of our model, we have included Table S

***Parameter Inference***

Given the complexity of our model, we incorporated as many publicly available datasets as possible to inform the inference of all model parameters and to minimize potential identifiability issues. This included integrating diverse datasets, including longitudinal data on relative abundances (ratios) [20], overall density of bacteria (cells/g of stool) [23], cross-sectional data on SIgA indexes [21], SIgA coating ratio [36], and the varying abundance of taxa with different feeding practices [22]. We have used the published abundance results on a taxonomic level. While we acknowledge that these data sources differ in sample origin, sequencing regions, and database references, they still represent the most comprehensive data available for our analysis. Table S1 summarizes the use of these data sources alongside their methods of taxonomic classification. Data used for model fitting can be found at https://github.com/burcutepekule/ONIMTOL/.

**Table S1. Taxonomic classification methods used across the studies for inference.** Summary of the taxonomic classification methods employed in studies used for inference, including details on the reference databases used, subjects, selection criteria, clustering methods, and bioinformatic tools.

| Study | Data used | Unit | Subjects | Selection | Taxonomic Classification |
|---|---|---|---|---|---|
| Tsukuda et al. [20] | Relative abundances during the first 2 years of life. | Ratios | Fecal samples of 12 human subjects during the first 2 years of life. | All subjects except Subject ID K (transition to solid food is not recorded.) were used since there were no subjects free of both antibiotic and probiotic exposure. Characteristics are provided in Table S1 of the respective study. | Taxonomy was assigned using the SILVA database (Release 138) with a 50% bootstrap threshold via the Qiime feature-classifier. Phylotypes were clustered using open-reference clustering against the NCBI 16S RefSeq records. |
| Palmer et al. [23] | Bacterial counts to scale the relative abundances to absolute abundances. | Cells/g of stool | Fecal samples of 14 healthy, full-term human infants during the first year of life. | Babies #7 and #9 who were delivered by vaginal birth, exclusively breastfed, and not exposed to antimicrobials. | Each sequence was taxonomically classified using the 2004 prokMSA taxonomy via BLAST alignment. This method compared sequences from 16S rRNA gene sequencing to the prokMSA taxonomy to assign taxonomic ranks. |
| Planer et al. [21] | IgA indexes for the Maternal Phase (days 60 and 120) and Steady Phase. | Unitless | Fecal samples of 40 healthy human twin pairs during the first 2 years of life. | Subjects are selected based on their 'Ratio of Breast milk: Formula' for the exclusive breastfeeding and mixed feeding periods. No information regarding antibiotic or probiotic exposure is provided. | Taxonomy was assigned using operational taxonomic units (OTUs) clustered with 97% identity against the GreenGenes 2013 reference database, analyzed with QIIME version 1.8. An abundance-filtered dataset was generated, only considering OTUs with relative abundances above 0.1% in at least 1% of the samples. |
| Pan et al. [22]. | Relative abundances for the case of no breastfeeding at day 30. | Ratios | Fecal samples of 100 healthy human newborns collected at day 3 and days 30-42 of life. | Analysis results based on all subjects are used. None of the subjects were on perinatal antibiotics. No probiotic data provided. | Sequences were clustered into OTUs using 16S rRNA sequencing, and classification was performed with Qiime 1.9.1. The reference database for genus-level classification was not specified. |
| van der Waaij et al. [36] | IgA coating ratio of the Steady Phase. | Ratio | Fecal samples of 15 healthy human adults (non-inflammatory controls in this study). | Analysis results based on non-inflammatory controls are used. Patients and controls had not used antibiotics within 2 weeks of sampling. No probiotic data provided. | This study did not use 16S rRNA sequencing. Instead, it relied on flow cytometry for measuring immunoglobulin-coated bacteria, without employing genus calling techniques. |

The complexity of our model relative to the volume of data posed challenges in simultaneously inferring all parameters. Therefore, we initially focused on the Maternal and Steady phases—time periods when the host's endogenous system is either not active or fully matured, respectively. This approach allowed us to ignore the parameters related to the developmental phase at this stage of inference and focus on a smaller set of shared parameters between the Maternal and Steady phases. Using the parameters estimated from this initial stage, we then proceeded to infer parameters relevant to the developmental phase.

*Joint parameter Inference of the Maternal and the Steady Phase*

The key observation which made this inference stage possible is the multigenerational transmission of the mucosal immune response [21], that the offspring converges to a similar SIgA response as the mother, i.e., $\mu_i{}^{e,n}(t) = \mu_i{}^{m,n}(t)$ and $\mu_i{}^{e,m}(t) = \mu_i{}^{m,m}(t)$ for $t \geq 720$Accordingly, we only need to infer one set of SIgA affinity values that are shared between the mother and the matured offspring. Since this stage of inference does not include the developmental phase, we cannot quantify the eSIgA levels as they are dynamically calculated via Eqn. S1.2.2., and thus assume that the eSIgA concentration reaches its carrying capacity of 1 for the matured offspring.

We employed Bayesian statistical analysis using the RStan software (Version 2.32.3; Stan Development Team) [126], and computed a composite likelihood function based on the simultaneous optimization of four different signals: *i)* relative abundances between days 3 and 274[1] representing the maternal phase, *ii)* relative abundances between days 721 and 735 representing the steady phase, *iii)* relative abundances at day 30 in case of no breastfeeding, and *iv)* SIgA indexes at the steady phase[2]. Error is assumed to be normally distributed for all signals. To manage signals of varying lengths and prevent overfitting to any particular signal, we established a precomputed upper bound for the likelihood of each signal, taking into account their length and the standard deviation applied in their likelihood calculations. Subsequently, we normalized the likelihood of each signal against this upper bound, capping it at 1 if it surpassed this limit, to ensure no single signal is disproportionately optimized. Parameters relevant to this inference stage are denoted by "Inferred-1" in Table S3 with their respective prior distributions.

*Parameter Inference of the Developmental Phase*

Using the parameters inferred from the previous stage, we move onto inferring parameters relevant for the developmental phase, denoted by "Inferred-2" in Table S3 with their respective prior distributions. We computed a composite likelihood function based on the SIgA affinities ($\rho^{p}{}_{\bar{i}}(t)$) and the total SIgA coating ratio of the microbiome observed in the fecal samples of the mature host [36], i.e., $\frac{w^{m}{}_{i}(t) + w^{n}{}_{i}(t)}{w\ {}_{i}(t)}$ for $t \geq 720$. Due to the non-smooth nature of the functions involved in this phase, we employ a grid-search algorithm [127] to infer the parameters to optimize the likelihood function.

---

[1]Given the metadata, the Maternal phase ends at day 172 when averaged over all subjects. After the data is interpolated and smoothed for inference, this time period was cut to 154 days. We then account for an additional 120 days adding up to 154+120 = 274 days for the symbiotic commensals including the post-weaning taxa (*Bifidobacteriaceae*, *Bacteroidaceae*, *Clostridiales*) to capture their interaction terms. However, we strictly used 172 days for the pathogenic taxon *Enterobacteriaceae* to avoid the impact of interaction to be mistaken for the neutralizing effects of the endogenous immune system starting to develop after day 154.

[2]To align with the feeding practices of the cohort *Tsuduka et al.*, we filtered the subjects in this dataset using the metadata provided on the ratio of breastmilk to formula feeding (B/F). We used a lower bound of 0.75 and an upper bound of 0.25 for the average B/F over the Maternal and the Developmental Phase, respectively, and picked the subjects who satisfied both criteria. This led to 154 days of exclusive breastfeeding and 308 days of mixed feeding on average.

*Data Curation and Pre-processing*

We used two datasets based on relative abundance data derived from stool samples during inference. First dataset is the 16S rRNA sequencing analysis combined with feeding metadata presented in *Tsukuda et al.* [20], which is based on longitudinally collected fecal samples from 12 subjects during the first 2 years of life. Based on their functional significance in determining the community profiles during early life, we have grouped our focal genera into four distinct taxonomic groups: *Enterobacteriaceae* ($E$), *Bifidobacteriaceae* ($B$)**,** *Bacteroidaceae* ($BC$)**,** and *Clostridiales* ($C$) (Table S2). Each group was selected for its unique role in influencing the model dynamics: $E$ encompasses potentially pathogenic bacteria (genera *Escherichia* and *Shigella*) encountered pre-weaning when the endogenous system is more vulnerable to enteric infections. $B$ represents the early colonizers with anti-inflammatory properties, regulating the microenvironment and providing colonization resistance. $BC$ and $C$ (*Anaerostipes*, *Blautia*, *[Eubacterium] hallii* group, *Faecalibacterium*, and *Ruminococcus*, all of which are non-pathogenic commensal bacteria contributing significantly to gut homeostasis) are the post-weaning bacteria with potential anti-inflammatory properties, which, unless regulated by SIgA, may stimulate the gut environment. We recognize that these taxonomic groups are not homogenous, and while some variability exists within each group—particularly under dysbiotic conditions—the classification reflects general trends observed in microbial behavior and immune modulation. Thus, although individual strains may diverge in their effects on gut homeostasis, this grouping provides a meaningful basis for modeling microbial-immune interactions.

Relative abundances provided in [20] are interpolated and smoothed using the loess function in R with a span parameter of 0.5, and weighed with a sigmoid function centered around the day of transition to MF (day 172 when averaged over all subjects based on the metadata) with a scale parameter of 5 to ensure a smooth transition from maternal to developmental phase (details can be found in our publicly available repository, https://github.com/burcutepekule/ONIMTOL/blob/main/misc/SMOOTH_DATA.R). These smoothed relative abundances are then converted to absolute abundances using data regarding fecal density counts (cells/g fecal content) [23], assuming that the total bacterial load stabilizes after 2 years (data extracted from Fig. 2 in [23]). Although $BC$ and $C$ were detected at low abundances in pre-weaning samples, we assume that their introduction primarily occurs with the transition to mixed feeding, thus, their abundances are considered negligible and set to zero in the maternal phase. We assume that the net growth rate encompasses the net influx of bacteria, and introduce the taxa once at their respective time point as modifications to the initial conditions of the differential equation system. Consequently, depending on the duration of EBF, MF, and the values of $HMOs(t)$ and $PDPs(t)$, we adjust the initial conditions for $B$ and $E$ at the start of the Maternal, and for $BC$ and $C$ at the start of the Developmental Phase. We assume that abundance of $B$ transferred from mother to offspring exponentially decreases to 25% of its maximum value as the EBF duration shortens from 3 to 0 days[3], mimicking the high bacterial counts in colostrum and its impact on initial microbial seeding [128]. $BC$ and $C$ abundances at the start of the

[3] To adjust the abundance of *Bifidobacteriaceae* for a given duration of EBF including the durations shorter than 3 days, we multiply the abundance that is transferred for the feeding patterns provided in the metadata with a multiplier of 0.25+0.75(1-exp(-2.207t)), ensuring a decreasing increase from 0.25 to 1 between days 0 and 3 and convergence to 1 for day 3 and onwards.

Developmental Phase are determined by the first non-zero value they take at time $t = t^*_{MF}$, and are scaled by $PDPs(t_{MF})/PDPs(t^*_{MF})$ to adjust the inoculum size introduced with MF accounting for the deviations from the dataset used for inference.

The second dataset by Pan *et al.* [22] consists of 16S rRNA sequencing data from infants categorized by feeding type: breastfed, partially breastfed, and formula-fed. Due to the lack of detailed information on the partially breastfed group, we focus on the cross-sectional abundance data from formula-fed infants to impose additional constraints on the inference procedure. Relative abundances of *E* (~32%), *B* (~21%), *BC* (~23%), and *C* (~12%) provided in [22] for formula fed infants at day 30 of life (https://github.com/burcutepekule/ONIMTOL/blob/main/PAN_DATA/Figure2a_quantified.png) are scaled according to the total relative abundance of *E*+*B*+*BC*+*C* (72%) at day 29 provided in [20] and incorporated into the composite likelihood function representing the scenario of no breastfeeding. Including this scenario provides valuable information to the inference process as to how different calorie sources (HMOs and PDPs) impact the growth rate of different taxa.

IgA indexes of human twins provided in [21] are filtered according to the 'Ratio of Breast milk: Formula' column in the dataset. For the exclusive breastfeeding period (until day 154, time of weaning in our model), 'Ratio of Breast milk: Formula'>0.75 is used. For the mixed feeding period (after day 154) this constraint is changed to 'Ratio of Breast milk: Formula'<0.5 to ensure that the subjects' diet timeline aligns with the ones in [20]. The code for processing the data provided in [21] is available in our repository, https://github.com/burcutepekule/ONIMTOL/blob/main/DATA_PLANER.R.

Differential growth capabilities and $O_2$ metabolism is provided in Table S2 [28–31,57]. We assumed a negative directionality for all intra-taxa competition terms and for the inter-taxa competition terms between the set $\{B, BC, C\}$ and $E$ [84] and informed the estimation of $\beta_{i,j}$ accordingly. However, determining the directionality of the remaining inter-taxa terms is not straightforward due to multiple mechanisms simultaneously dictating these interactions such as competition and cross-feeding [129–133], thus they were not informed *a priori*.

**Table S2. Key taxonomic groups and their corresponding inoculation time, oxygen ($O_2$), and carbohydrate metabolism.**

| Family / Order | Genus | Inoculation | $O_2$ metabolism | Carbohydrate metabolism |
|---|---|---|---|---|
| *Enterobacteriaceae* | *Escherichia-Shigella* | At birth. | Facultative anaerobes, $\phi_E^{O_2} = 0$. | Mostly PDPs, $\phi_E^{HMOs}$ and $\phi_E^{PDPs}$ are inferred from data. |
| *Bifidobacteriaceae* | *Bifidobacterium* | At birth and with milk within the first 3 days. | Strict anaerobes, $\phi_B^{O_2} = -1$. | Mostly HMOs, but also PDPs, $\phi_B^{HMOs}$=1 and $\phi_B^{PDPs}$ are inferred from data. |
| *Bacteroidaceae* | *Bacteroides* | With mixed feeding. | Strict anaerobes, $\phi_{BC}^{O_2} = -1$. | Mostly PDPs, but also HMOs, $\phi_{BC}^{HMOs}$ is inferred from data and $\phi_{BC}^{PDPs} = 1$. |

| | | | | |
|---|---|---|---|---|
| *Clostridiales* | *Anaerostipes, Blautia, [Eubacterium] hallii group, Faecalibacterium, Ruminococcus* | With mixed feeding. | Strict anaerobes, ${\phi_C}^{O_2} = -1.$ | Mostly PDPs, but also HMOs, ${\phi_C}^{HMOs}$ is inferred from data and ${\phi_C}^{PDPs} = 1.$ |

*Inference Results and Values of Model Parameters*

We use three different quantification types to indicate how parameters are quantified : *i)* "Inferred-1" indicates that the value of the parameter is estimated during the joint parameter inference of the Maternal and the Steady Phase, *i)* "Inferred-2" indicates that the value of the parameter is estimated during the joint parameter inference of the Developmental Phase, *iii)* "Dependent" indicates that the quantification of the parameter depends on the value of other parameters, *iv)* "Assumed Based on Literature" (ABL) refers to parameters for which quantitative values are derived from other quantitative or qualitative data sources and expert opinion available in the literature, and *v)* "Calibrated" refers to parameters that are tuned to achieve the values quantified via the former first four quantification methods. Similar quantification methods apply to parameterize the prior distributions used during inference, where they are either ABL or calibrated.

Given the absence of quantitative data for specific variables and parameters within our system, we employ normalization to assess their temporal influence on model dynamics. This approach applies to variables such as $O_2$, $mSIgA$, and $eSIgA$. Their impact on the system is maximized when their normalized value reaches 1 and minimized at the value of 0.

**Table S3. Parameters with their corresponding descriptions, units, ranges, prior distributions, and model fit estimates.**

| **Notation** | **Description** | **Units** | **Quantification** | **Range/ Constraints** | **Prior Dist./ Value** | **Model fit estimate** |
|---|---|---|---|---|---|---|
| **State Variables** | | | | | | |
| $mSIgA$ | Maternal SIgA concentration in the gut lumen. | Unitless | - | [0,1] | | |
| $eSIgA$ | Endogenous SIgA concentration in the gut lumen. | Unitless | - | [0,1] | | |
| $O_2$ | Oxygen concentration in the gut lumen. | Unitless | - | [0,1] | | |
| ${y_i}^L$ | Absolute abundance of taxon $i$ in the gut lumen. | Cells/gLC | - | (0, ∞) | | |
| ${y_i}^{L,uc}$ | Absolute abundance of SIgA- taxon $i$ in the gut lumen. | Cells/gLC | - | (0, ∞) | | |
| ${y_i}^{L,c}$ | Absolute abundance of SIgA+ (C) taxon $i$ in the gut lumen. | Cells/gLC | - | (0, ∞) | | |
| ${y_i}^{L,\Sigma n}$ | Cumulative absolute abundance of neutralized taxon $i$ in the gut lumen. | Cells/gLC | - | (0, ∞) | | |
| $\delta_i$ | Selection threshold for taxon $i$. | Unitless | - | (0, ∞) | | |
| ${B_i}^n$ | Number of naïve B cells available for entering circulation, dedicated to taxon $i$. | Unitless | - | (0, ∞) | | |

| | | | | | | |
|---|---|---|---|---|---|---|
| $B_i^c$ | Number of circulating B cells within the GCs, dedicated to taxon $i$. | Unitless | - | (0, ∞) | | |
| $B_i^p$ | Number of plasma cells in LP, dedicated to taxon $i$. | Unitless | - | [0,1] | | |
| $\bar{\rho}_i^c$ | Average BCR affinity of circulating B cells, dedicated to taxon $i$. | Unitless | - | [0, ∞) | | |
| $\bar{\rho}_i^p$ | Average BCR affinity of plasma B cells, dedicated to taxon $i$. | Unitless | - | [0, ∞) | | |
| $\sigma_i^c$ | Standard deviation of the BCR affinity distribution of circulating B cells, dedicated to taxon $i$. | Unitless | - | [0, ∞) | | |
| **Time-independent parameters** | | | | | | |
| $\mu_{O2}$ | Degradation rate of $O_2$ by facultative anaerobes. | 1/ (days x Cells/gLC) | Inferred-1 | (0,1) | $\Gamma(10,6)$[4] | 1.35 |
| $\phi_i^{O_2}$ | Impact of factor $O_2$ on the net growth rate of taxon $i$. | Unitless | ABL [45] | {0, 1} | - | $\phi_E^{O_2} = 0$ <br> $\phi_B^{O_2} = -1$ <br> $\phi_{BC}^{O_2} = -1$ <br> $\phi_C^{O_2} = -1$ |
| $\phi_i^{HMOs,PDPs}$ | Impact of factor HMOs and PDPs on the net growth rate of taxon $i$. | Unitless | Inferred-1 | $0 < \phi_E^{HMOs} < 0.9$, <br> $\phi_E^{HMOs} < \phi_C^{HMOs} < 0.9$, <br> $\phi_C^{HMOs} < \phi_{BC}^{HMOs} < 0.9$, <br> $0 < \phi_E^{PDPs} < 0.9$, <br> $\phi_E^{PDPs} < \phi_B^{PDPs} < 0.9$. | $\beta(5,5)$[5] | $\phi_E^{HMOs} = 0.31$ <br> $\phi_{BC}^{HMOs} = 0.61$ <br> $\phi_C^{HMOs} = 0.47$ <br> $\phi_E^{PDPs} = 0.48$ <br> $\phi_B^{PDPs} = 0.63$ |
| $\lambda_i$ | Net growth rate of taxon $i$. | 1/days | Inferred-1 | (0, ∞) | Γ (3,2) | $\lambda_E = 1.323$ <br> $\lambda_B = 1.19$ <br> $\lambda_{BC} = 1.00$ <br> $\lambda_C = 1.34$ |
| $\lvert\beta_{i,j}\rvert$ | Magnitude of impact of taxon $j$ abundance on taxon $i$ abundance for each pair of $\{i,j\}$, where the directionality of impact is known *a priori.* | 1/ (days x Cells/gLC) | Inferred-1 | (0, ∞) | Γ (2,0.1) | $\beta_{E,E} = -56.61$ <br> $\beta_{E,B} = -28.31$ <br> $\beta_{E,BC} = -14.94$ <br> $\beta_{E,C} = -24.40$ <br> $\beta_{B,E} = -2.43$ <br> $\beta_{B,B} = -5.92$ <br> $\beta_{BC,E} = -6.16$ <br> $\beta_{BC,BC} = -17.64$ <br> $\beta_{C,E} = -1.90$ <br> $\beta_{C,C} = -5.28$ |
| $\beta_{i,j}$ | Value of impact of taxon $j$ abundance on taxon $i$ abundance for each | 1/ (days x Cells/gLC) | Inferred-1 | (-∞, ∞) | Ɲ(0,10)[6] | $\beta_{B,BC} = -3.70$ <br> $\beta_{B,C} = -4.73$ |

[4] Γ(α,β) denotes the gamma distribution, where α and β denote the shape and the rate parameter, respectively.

[5] β(α,β) denotes the beta distribution, where α and β denote the shape parameters.

[6] Ɲ(μ,σ) denotes the normal distribution, where μ and σ denote the mean and the standard deviation, respectively.

| | | | | | | |
|---|---|---|---|---|---|---|
| | pair of $\{i,j\}$, where the directionality of impact is not known *a priori.* | | | | | $\beta_{BC,B} = 10.67$<br>$\beta_{BC,C} = -15.02$<br>$\beta_{C,B} = -5.49$<br>$\beta_{C,BC} = -2.14$ |
| $\rho_i$ | Average mSIgA affinity, specific to taxon $i$. | Unitless | Inferred-1 | [9,99] if $i \in \{E\}$,<br>(0.10,0.25] if $i \in \{B\}$,<br>(0,0.10] if $i \in \{BC, C\}$. | U (9,99)[7] if $i \in \{E\}$,<br>$\Gamma$(5,100) if $i \in \{B\}$,<br>$\Gamma$ (20,100) if $i \in \{BC, C\}$. | $\rho_E = 9.52$<br>$\rho_B = 0.05$<br>$\rho_{BC} = 0.18$<br>$\rho_C = 0.18$ |
| $\mu_i^{m,m}$ | Masking rate of mSIgA, specific to taxon $i$. | 1/days | Dependent | 0 if $i \in \{E\}$,<br>[0.8,1) if $i \notin \{E\}$. | - | $\mu_E^{m,m} = 0.10$<br>$\mu_B^{m,m} = 0.95$<br>$\mu_{BC}^{m,m} = 0.84$<br>$\mu_C^{m,m} = 0.84$ |
| $\mu_i^{m,n}$ | Neutralizing rate of mSIgA, specific to taxon $i$. | 1/days | Dependent | [0.8,0.95] if $i \in \{E\}$,<br>0 if $i \notin \{E\}$. | - | $\mu_E^{m,n} = 0.90$<br>$\mu_B^{m,n} = 0.05$<br>$\mu_{BC}^{m,n} = 0.16$<br>$\mu_C^{m,n} = 0.16$ |
| $r_d$ | Dissociation rate of SIgA | Unitless | Inferred-1 | (0, 1) | $\Gamma$ (10,20) | 0.602 |
| $\omega_i$ | Binding ability of mSIgA, specific to taxon $i$. | Unitless | Dependent | [0, 1] | - | $\omega_E = 0.94$<br>$\omega_B = 0.43$<br>$\omega_{BC} = 0.49$<br>$\omega_C = 0.49$ |
| $\epsilon^{uc}$ | Antigenic-sampling rate $y_i^{L,uc}$ for any taxon $i$. | Unitless | Calibrated | $\epsilon^c > \epsilon^{uc}$ | - | 0.005 |
| $\epsilon^c$ | Antigenic-sampling rate $y_i^{L,c}$ for any taxon $i$. | Unitless | Calibrated | $\epsilon^c > \epsilon^{uc}$ | - | 0.05 |
| $\alpha_i$ | Relative invasiveness of taxon $i$. | Unitless | Inferred-2 | $\alpha_E > max\{\alpha_{BC}, \alpha_C\} > \alpha_B$,[39] | $\Gamma$ (2,250) if $i \in \{B\}$,<br>$\Gamma$ (2,50) if $i \in \{BC, C\}$. | $\alpha_E = 1$ (fixed)<br>$\alpha_B = 0.008$<br>$\alpha_{BC} = 0.036$<br>$\alpha_C = 0.038$ |
| $\kappa_i$ | Relative immunostimulatory capacity of the SIgA- members of taxon $i$. | Unitless | Inferred-2 | A priori distribution is parameterized given the relative immunostimulatory capacities reported in the literature [36,37]. | $\Gamma$ (30,0.1) if $i \in \{E\}$ | $\kappa_E = 314.1$<br>$\kappa_B = 1$ (fixed)<br>$\kappa_{BC} = 1$ (fixed)<br>$\kappa_C = 1$ (fixed) |
| $I_i$ | Binary variable representing the inherent anti-inflammatory potential of taxon $i$. | Unitless | ABL [118–120] | 0 if $i \in \{E\}$,<br>1 if $i \notin \{E\}$. | - | $I_E = 0$<br>$I_B = 1$<br>$I_{BC} = 1$<br>$I_C = 1$ |
| $C_n$ | Amplitude of the exponential function describing the diminishing pool of naïve T and B cells. | Unitless | Calibrated | - | - | 0.03 |
| $c_n$ | Decay rate of the exponential function describing the diminishing pool of naïve T and B cells. | Unitless | Calibrated | - | - | 0.000634 |
| $th_{apop}$ | Selection threshold multiplier determining the | Unitless | Calibrated | - | - | 0.25 |

7 U(a,b) denotes the uniform distribution, where a and b denote the upper and lower bounds, respectively.

| | | | | | | |
|---|---|---|---|---|---|---|
| | minimum BCR affinity to get sufficient T cell help to avoid apoptosis. | | | | | |
| $th_{high}$ | Selection threshold multiplier determining the minimum BCR affinity to get sufficient T cell help to differentiate to plasma cells. | Unitless | Calibrated | - | - | 0.75 |
| $th_{ang}$ | Selection threshold multiplier determining the minimum BCR affinity for B cells to be rendered anergic through regulatory mechanisms. | Unitless | Calibrated | - | - | 1.25 |
| $r_p$ | Proliferation rate of circulating B cells in the dark zone of the Germinal Center. | 1/days | Calibrated | - | - | 1[8] |
| $\tau^c$ | Multiplier to calculate the additional standard deviation in BCR affinity distribution after somatic hypermutation. | Unitless | Inferred-2 | - | Γ (2,0.05) | 28.33 |
| $\tau^{new}$ | Multiplier to calculate the additional standard deviation in BCR affinity distribution in newly activated B cells. | 1/(Cells/gLC) | Inferred-2 | - | Γ (10,0.05)[9] | 180.21 |
| $\tau^{\delta}$ | Multiplier to adjust the incremental increase in the selection threshold calculated in Eqn. S1.2.8. | Unitless | Inferred-2 | - | Γ (2,5) | 0.42 |
| $t^m$ | Time of M cell activation and the start of antigenic sampling. | Days | - | - | - | $(t^{50}, 30)$ |
| $t_{MF}$ | Time of switching to MF from EBF. | Days | - | [0,180] | - | - |
| $t^*_{MF}$ | Average time of switching to MF from EBF in the dataset used for parameter inference after data pre-processing. | Days | - | - | - | 154 |
| $T_{EBF}$ | Duration of EBF. | Days | - | [0,180] | - | - |
| $T_{MF}$ | Duration of MF. | Days | - | [0,300] | - | - |

8 Rapidly proliferating B cells in the dark zone of the germinal centers have cell cycles typically between 6 and 12 h [121]. Here we assume a 12h cycle, meaning the cell count doubles within a day.

9 *A priori* distributions for $\tau^c$ and $\tau^{new}$ are parametrized to reflect the significant contribution of somatic hypermutation (SHM) relative to the contribution of newly activated B cells in increasing BCR diversity. While some degree of affinity maturation can occur in the absence of germinal centers, the extent and efficiency of this process are substantially greater with SHM. SHM targets the variable regions of BCR genes for high-rate mutations, leading to a dramatic increase in BCR diversity and specificity. This mechanism far surpasses the initial diversity provided by newly activated B cells through V(D)J recombination, crucially enhancing the immune system's ability to fine-tune and strengthen responses to specific antigens [122].

| | | | | | | |
|---|---|---|---|---|---|---|
| $T^*_{MF}$ | Average duration of MF in the dataset used for parameter inference after data pre-processing. | Days | - | - | - | 308 |
| $K_{H1}$ | $f_{HMOs}(t, t_{MF}, T_{MF})$ parameter (Eqn. S1.1.18) | Unitless | Calibrated | - | - | 0.44 |
| $V_0$ | $f_{HMOs}(t, t_{MF}, T_{MF})$ parameter (Eqn. S1.1.18) | Unitless | Calibrated | - | - | 0.2 |
| $c_H$ | $f_{HMOs}(t, t_{MF}, T_{MF})$ parameter (Eqn. S1.1.18) | 1/days | Calibrated | - | - | 40 |
| $K_C$ | $f_{TOT}(t, t^*_{MF}, T^*_{MF})$ parameter (Eqn. S1.1.19) | Unitless | Calibrated | - | - | 1.28 |
| $c_c$ | $f_{TOT}(t, t^*_{MF}, T^*_{MF})$ parameter (Eqn. 1.1.19) | 1/days | Calibrated | - | - | 0.005 |
| $t_C$ | $f_{TOT}(t, t^*_{MF}, T^*_{MF})$ parameter (Eqn. 1.1.19) | Days | Calibrated | - | - | 720 |
| $K_I$ | $\Delta f_{mSIgA}(t)$ parameter (Eqn. 1.1.21) | Unitless | Calibrated | - | - | 0.15 |
| $m_I$ | $\Delta f_{mSIgA}(t)$ parameter (Eqn. 1.1.21) | Unitless | Calibrated | - | - | 0.38 |
| $c_I$ | $\Delta f_{mSIgA}(t)$ parameter (Eqn. 1.1.21) | 1/days | Calibrated | - | - | 0.0316 |
| $C_I$ | Maximum secretion capacity of plasma cells. | Unitless | Inferred-2 | - | Γ (10,1) | 12.39 |
| $I_{hl}$ | Half Life of SIgA in the gut lumen. | Days | ABL [123] | - | - | 5 |
| $\psi^{c}$ | Activation rate of naïve B cells per unit of SIgA-antigen complex. | 1/(Cells/gLC) | Calibrated | - | - | 0.1 |
| $\psi^{uc}$ | Activation rate of naïve B cells per unit of SIgA-free antigen. | 1/(Cells/gLC) | Calibrated | - | - | 1 |
| $s^m$ | Observation rate of masked bacteria in fecal samples. | Unitless | Inferred-1 | [0,1] | $\beta$(5,4) | 0.48 |
| $s^{uc}$ | Observation rate of uncoated bacteria in fecal samples. | Unitless | Inferred-1 | [$s^c$, 1] | $\beta$(10,4) | 0.82 |
| $s^n$ | Observation rate of neutralized bacteria in fecal samples. | Unitless | Inferred-1 | [$s^{uc}$, 1] | $\beta$(20,4) | 0.85 |
| **Time-dependent parameters** | | | | | | |
| $\lambda_i^{adj}$ | Adjusted growth rate of taxon $i$ based on $\lambda_i$, $\phi_i^x$, $O_2(t)$, $HMOs(t)$, and $PDPs(t)$. | 1/days | Dependent | Determined by Eqn. S1.1.4. | - | - |
| $\mu_i^{e,c}$ | Coating rate of taxon $i$ induced by endogenous SIgA. | 1/days | Dependent | Determined by Eqn. S1.1.1. | - | - |
| $\mu_i^{e,n}$ | Neutralization rate of taxon $i$ induced by endogenous SIgA. | 1/days | Dependent | Determined by Eqn. S1.1.2. | - | - |

| | | | | | | |
|---|---|---|---|---|---|---|
| $\omega_i^{e,n}$ | Binding ability of neutralizing eSIgA, specific to taxon $i$. | Unitless | Dependent | Determined by Eqn. S1.1.3. | [0,1] | - |
| $\omega_i^{e,c}$ | Binding ability of coating eSIgA, specific to taxon $i$. | Unitless | Dependent | Determined by Eqn. S1.1.3. | [0,1] | - |
| $\sigma$ | Microenvironmental stimulation | Unitless | Dependent | Determined by Eqn. S1.1.8. | - | - |
| $t^{50}$ | Time point when mSIgA concentration falls below 50% of its peak value, representing the beginning of antigenic sampling and activation of the endogenous immune system. | Days | Dependent | - | - | - |
| $z_i^{uc}$ | IgA-free antigens transported to Peyer's Patches through M cells. | Cells/gLC | Dependent | Determined by Eqn. S1.1.6. | - | - |
| $z_i^{c}$ | IgA-antigen immune complexes transported to Peyer's Patches through M cells. | Cells/gLC | Dependent | Determined by Eqn. S1.1.7. | - | - |
| $y_{fn}^{L}$ | Abundance of facultative anaerobes in the gut lumen. | Cells/gLC | Dependent | - | - | - |
| $\psi_i$ | Total B cell activation rate dedicated to taxon $i$. | 1/Cells/gLC | Dependent | Determined by Eqn. S1.1.5. | - | - |
| $B_i^{c,new}$ | Newly activated B cells dedicated to taxon $i$. | Unitless | Dependent | $\psi_i B_i^{n}$ | - | - |
| $w^{n}{}_i$ | Absolute abundance of neutralized bacteria observed per fecal content for taxon $i$. | Cells/gFC | Dependent | Determined by Eqn. S1.1.14. | - | - |
| $w^{m}{}_i$ | Absolute abundance of masked bacteria observed per fecal content for taxon $i$. | Cells/gFC | Dependent | Determined by Eqn. S1.1.15. | - | - |
| $w^{uc}{}_i$ | Absolute abundance of uncoated bacteria observed per fecal content for taxon $i$. | Cells/gFC | Dependent | Determined by Eqn. S1.1.16. | - | - |
| $w_i$ | Absolute abundance of total bacteria observed per fecal content for taxon $i$. | Cells/gFC | Dependent | Determined by Eqn. S1.1.17. | - | - |

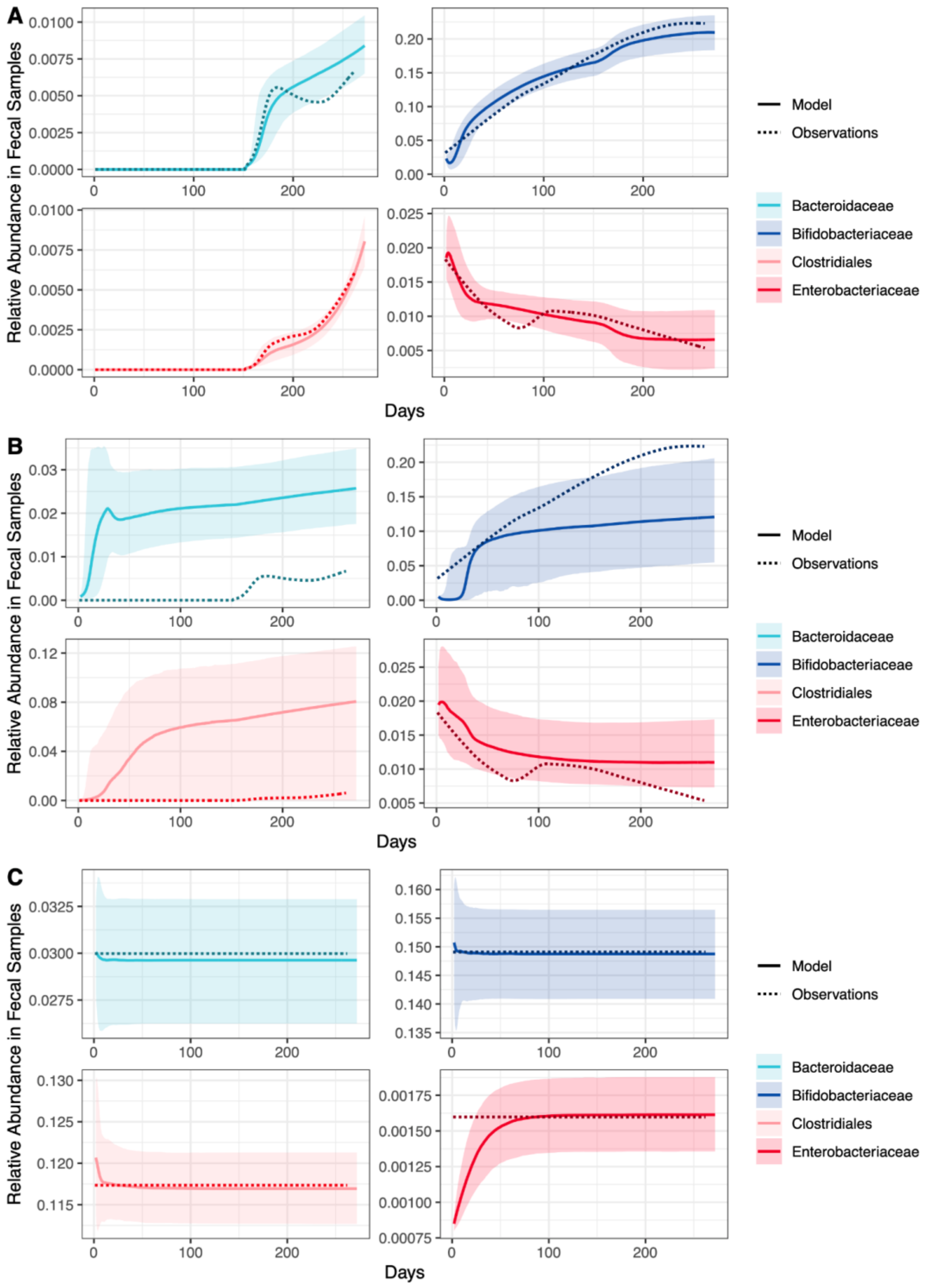

**Fig S1. Inference results of RStan.** 4 chains are used with 500 and 1000 for warm-up and total iterations, respectively. 122 of 2000 (6.0%) transitions ended with a divergence. (**A**) Relative abundance estimate results for the maternal phase, for 154 days of exclusive breastfeeding (EBF) and 308 days of mixed feeding (MF). (**B**) Relative abundance estimate results for the maternal phase, for 308 days of mixed feeding (MF) with no exclusive breastfeeding. (**C**) Relative abundance estimate results for the steady phase, for 154 days of exclusive breastfeeding (EBF) and 308 days of mixed feeding (MF). Shaded areas represent the 95% confidence intervals.

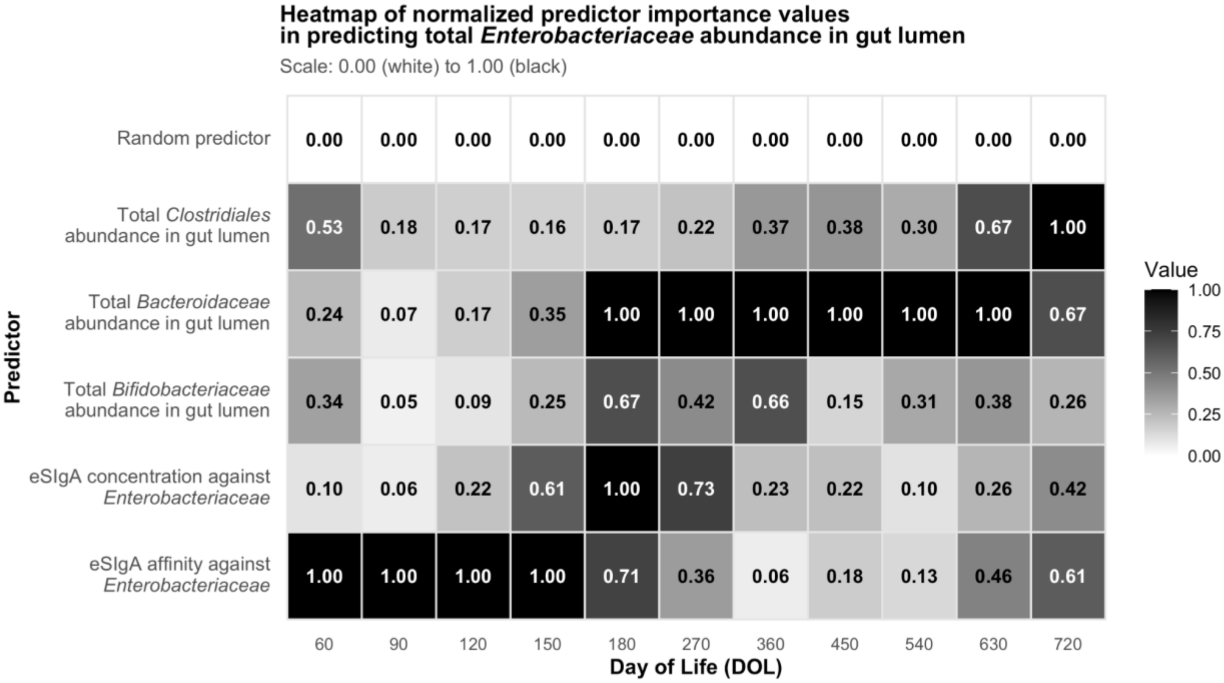

**Fig S2. Heatmap of normalized predictor importance values across time points from random forest models.** Each cell shows the normalized importance of a predictor in explaining total fecal abundance of *Enterobacteriaceae* at a given time point (in days), as determined by conditional permutation importance in a random-forest model. Predictors include eSIgA affinity and concentration against *Enterobacteriaceae*, and total abundance of *Bifidobacteriaceae*, *Bacteroidaceae*, and *Clostridiales* in the gut lumen. A random predictor was included as a negative control. Predictor importance values were normalized within each time point between 0 and 1. The most influential variable in any column is black (value = 1.00) and progressively lighter shades indicate lower relative importance. Numeric values are overlaid for clarity. Starting from month 6 (DOL 180), endogenous immune responses against symbiotic commensals combined with ecological competition become the primary regulators of *Enterobacteriaceae* population, exerting stronger selection pressure than the endogenous SIgA (eSIgA) responses to *Enterobacteriaceae* itself, as seen from the decreasing importance of both the affinity and the concentration of eSIgA against *Enterobacteriaceae*. By DOL 720, importance values are relatively evenly distributed across predictors, consistent with the similarity in predictive power between total and SIgA-bound *Enterobacteriaceae* in Fig. 3E.

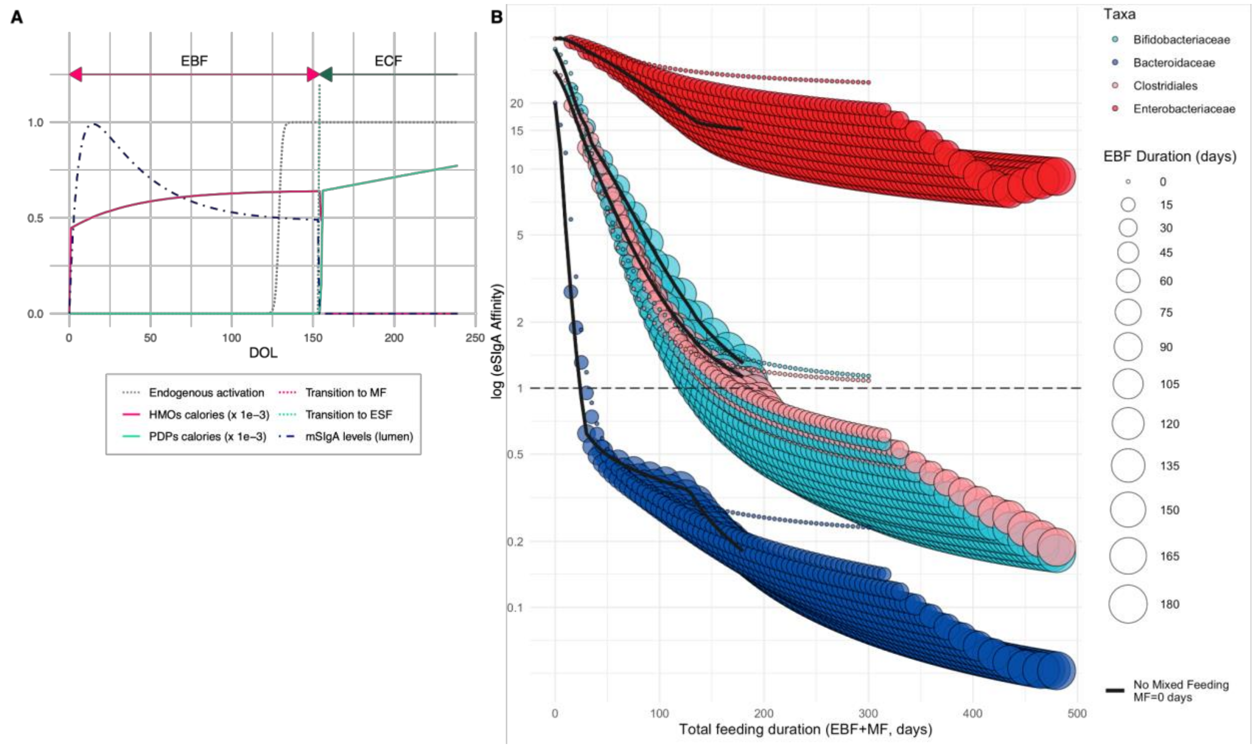

**Fig S3. Illustration of model inputs without mixed feeding and differential impacts of EBF and MF durations in determining eSIgA affinity across a comprehensive range of feeding durations.** (**A**) Model inputs demonstrating a sharp transition from exclusive breastfeeding (EBF) to exclusive solid feeding (ECF), with no mixed feeding (MF) period in between, including normalized maternal secretory immunoglobulin A (mSIgA) concentration, human milk oligosaccharide (HMOs) and plant-derived polysaccharides (PDPs) calorie inputs, and timing of the endogenous immune system activation over time. EBM: exclusive breastfeeding; MF: mixed feeding; ECF: exclusive complementary feeding. (**B**) log (eSIgA Affinity) values at steady state for different combinations of EBF and MF durations, where the solid black line demonstrates the case of no MF followed by EBF. DOL: Day of life; E: *Enterobacteriaceae*; B: *Bifidobacteriaceae*; BC: *Bacteroidaceae*; C: *Clostridiales*.

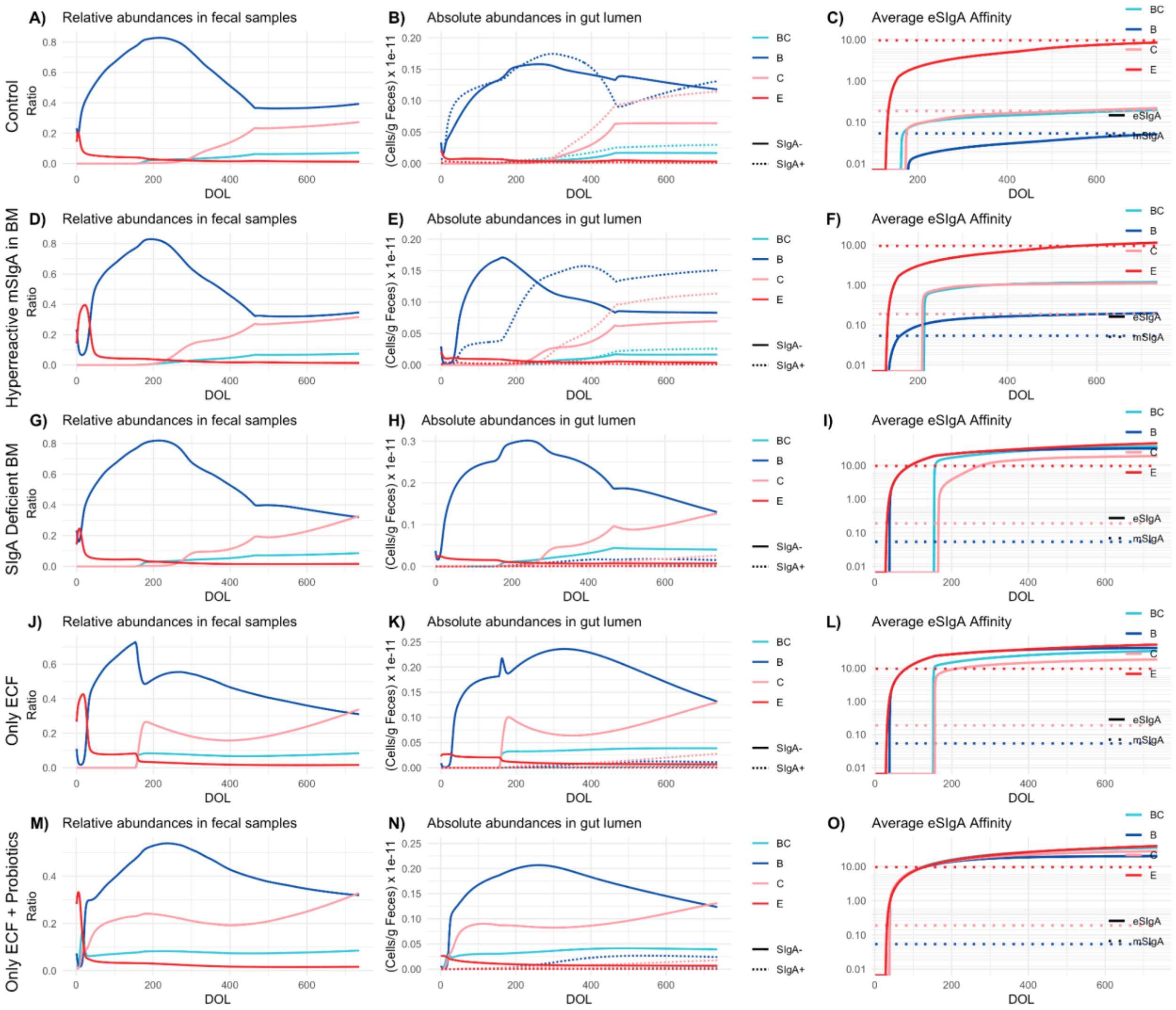

**Fig S4. Comparison of various breastfeeding scenarios and their impact on endogenous affinity maturation.** Relative abundances in fecal samples, absolute abundances in the gut lumen, and temporal progression of average endogenous SIgA (eSIgA) affinities for (**A**)-(**C**) control, (**D**)-(**F**) hyperreactive mSIgA in breastmilk (BM), (**G**)-(**I**) SIgA deficient BM, (**J**)-(**L**) only exclusive complementary feeding (ECF), and (**M**)-(**O**) ECF with probiotic (*Bacteroidaceae* and *Clostridiales*) supplementation. DOL: Day of life; E: *Enterobacteriaceae*; B: *Bifidobacteriaceae*; BC: *Bacteroidaceae*; C: *Clostridiales*.

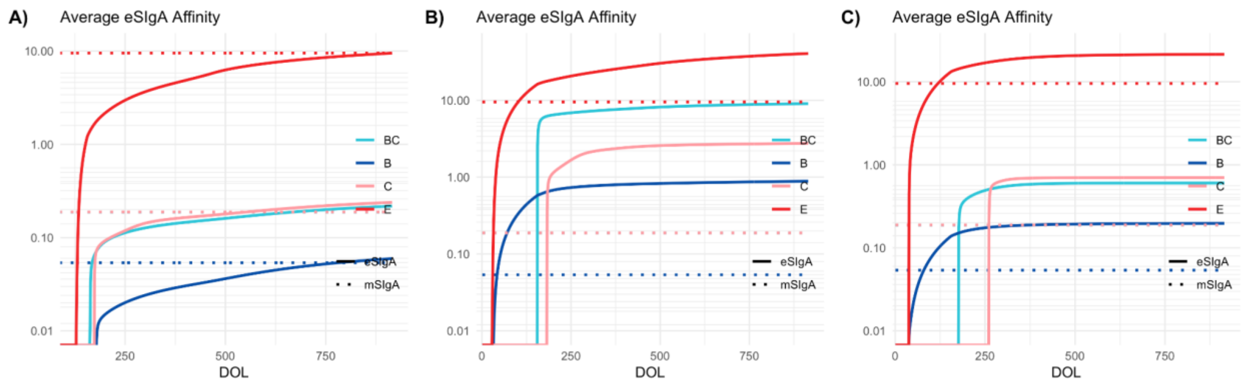

**Fig. S5. Hypothetical scenario demonstrating the administration of TLR4 Antagonists to prevent inflammatory imprinting when mSIgA in breastmilk is insufficient.** Temporal progression of average endogenous SIgA (eSIgA) affinities for (**A**) control**,** (**B**) when mSIgA levels are 85% reduced, (**C**) when mSIgA levels are 85% reduced with TLR4 antagonists' administration (90% reduction in TLR4 stimulation). DOL: Day of life; E: *Enterobacteriaceae*; B: *Bifidobacteriaceae*; BC: *Bacteroidaceae*; C: *Clostridiales*.

**Table S4. Additional assumptions implicit to the model structure.**

| Assumption | References |
|---|---|
| We assume that the initial exposure to microbes occurs at birth and that the intrauterine environment is sterile, although this traditional view has been recently challenged and it has been shown that bacteria existed as low-abundance, low-biomass and sparse populations, in utero bacterial colonization did occur during healthy pregnancy. | [134,135] |
| Although SIgA titers and the proportion of IgA-coated bacteria are generally higher in the small intestine, our analysis is based on fecal sample data, which predominantly reflects the microbial composition of the colon. We therefore assume that the IgA responses observed in our data represent a combined mucosal immune response, with a dominant contribution from colonic organized lymphoid tissues. Although some of the IgA coating may result from cross-reactivity—where antibodies generated in Peyer's patches of the small intestine in response to small-intestinal bacteria bind to structurally similar antigens on colonic bacteria—the observed taxon specificity of IgA binding in fecal samples suggests that many of these bacteria are likely true immunogens, eliciting local IgA responses within gut-associated inductive sites in the colon.<br><br>Colonic patches, like Peyer's patches, are organized inductive structures within the gut-associated lymphoid tissue (GALT). They exhibit key immunological features required for IgA induction, including M cell-containing follicle-associated epithelium, germinal centers with defined dark and light zones (as observed in canonical secondary lymphoid tissues and inducible mucosal lymphoid structures), high endothelial venules (HEVs), and segregated B and T cell areas. Given this structural and | [34,136–145] |

| | |
|---|---|
| functional similarity, our model focuses on the germinal center dynamics within such organized lymphoid tissues in the gut mucosa, without explicitly distinguishing between small-intestinal and colonic sites. Note that while scientific nomenclature sometimes includes mesenteric lymph nodes (MLNs) within the definition of GALT, we follow the classification presented by Mörbe *et al.* [34], which distinguishes between GALT structures (Peyer's patches and colonic patches) embedded within the intestinal wall and the separate intestine-draining MLNs. Thus, our definition of GALT excludes MLNs. | |
| Human SIgA is highly somatically hypermutated (SHM), and SHM is commonly linked to T cell-dependent (TD) germinal center (GC) reactions, although not completely excluding T-cell independent (TI) mechanisms. While the contribution of TI pathways is well established in mice, particularly in early life, significant differences exist between mice and human infants. In mice, mucosal immune structures like PPs develop postnatally, promoting stronger TI responses initially. In contrast, human PPs, colonic patches, and isolated lymphoid follicles (ILFs) are organized by 22 weeks of gestation, enabling earlier initiation of GC reactions and favoring TD pathways even in infancy. This greater maturity in human infants allows them to initiate germinal center (GC) reactions earlier than mice. Moreover, many bacterial outer membrane components, including carbohydrate structures like lipopolysaccharides (LPS), are often attached to surface proteins, which can trigger TD responses even in early life given that the model organism is mature enough to do so. In light of this evidence, we assume that all the IgA producing cells are the product TD GC reactions, and no class-switch-recombination occurs in the lamina propria. | [34,50,146–149] |
| In lymphoid organs associated with the gut, germinal centers (GCs) are chronically present. B-cell clones induced in early life are maintained through chronic GCs in the gut, which serve as the predominant reservoir responsible for SIgA-secreting plasma cell maintenance. | [150–152] |
| Germinal center-derived, affinity matured B cell responses towards defined antigens are important for the effects of SIgA on the microbiota. Therefore, SIgA responses to gut commensals are assumed to be antigen-specific. | [55] |
| We assume that different taxonomic groups induce SIgA with different affinities. It has been shown that different commensal bacterial species are coated by IgA to varying extents (IgA-Seq data from Planer *et al.* [21], used for model fitting). Although the differential coating suggests that IgA affinities are different for different bacterial taxa, it does not provide direct proof because the established techniques do not account for the biases that might arise from using relative abundances when sorting IgA-bound and - | [21,55,153] |

| |
|---|
| unbound fractions. However, recent work by Jackson *et al* [153] proposed a probabilistic scoring method designed to address these biases, including the use of relative abundances, by adjusting for the bacterial composition before sorting and quantifying the likelihood of IgA binding. By showing varying degrees of IgA binding to different commensal bacteria after adjusting for the biases described, this study indicates strong likelihood of differences in IgA affinities. Combined with strain-specific IgA responses in the gut [55], we believe that it is appropriate to assume the affinities would vary. |

#### *Global Sensitivity Analysis*

We implemented a global sensitivity analysis using the Morris method, which is a variance-based approach suitable for models with a large number of parameters [154]. The Morris method uses an OAT (One-At-a-Time) design to evaluate the impact of each parameter by varying them individually across a specified range. For this analysis, we considered 14 parameters: $t^m$, $\psi^m$, $C_I$, $C_n$, $c_n$, $th_{apop}$, $th_{range}$, $\tau^{new}$, $\tau^{\delta}$, $\tau^c$, $\alpha_B$, $\alpha_{BC}$, $\alpha_C$, $\kappa_E$ and 1 composite parameter: $\epsilon^m/\epsilon^{uc}$. We set up the Morris design with 500 trajectories, 10 levels per factor, and a grid jump of 2, resulting in 10000 total model evaluations (500 trajectories times 20 steps per trajectory). We used the average endogenous SIgA affinity at DOL 735 against symbiotic commensals (*Bifidobacteriaceae*, *Bacteroidaceae* and *Clostridiales*) as the output variable. For parameters with a prior distribution (see Table S3), lower and upper bounds of the sampling ranges are based on the 5th and 95th percentiles of their respective prior distributions. Results are provided in Table S5, with the parameters' respective descriptions, ranges of sampling, inferred/calibrated values used in our model, units, and sensitivity metrics including $\mu$ (mean elementary effect, indicating the overall influence of each parameter), $\mu^*$(mean absolute elementary effect, which captures the magnitude of the parameter's impact regardless of direction), and $\sigma$ (representing the variability or nonlinearity of the parameter's effect on the output).

**Table S5. Results of the global sensitivity analysis.** Results of the global sensitivity analysis using the Morris method. This table lists the parameters included in the analysis, their respective descriptions, the ranges used for sampling, their inferred or calibrated values in the model, units, and the sensitivity metrics: $\mu$ (mean elementary effect, representing the overall influence of each parameter), $\mu^*$ (mean absolute elementary effect, indicating the magnitude of the parameter's impact irrespective of direction), and σ (standard deviation of the elementary effects, representing the variability or nonlinearity in the parameter's effect on the output). The output variable considered in this analysis is the average endogenous SIgA affinity at DOL 735 against symbiotic commensals (*Bifidobacteriaceae*, *Bacteroidaceae*, and *Clostridiales*). The design includes 500 trajectories with 20 steps per trajectory, resulting in 10,000 total model evaluations.

| **Parameter (Fig. S6)** | **Description** | **Range of sampling** | **Inferred /calibrated value used in model** | **Unit** | **mu ($\mu$)** | **mu.star ($\mu^*$)** | **sigma ($\sigma$)** |
|---|---|---|---|---|---|---|---|

| | | | | | | | |
|---|---|---|---|---|---|---|---|
| $\epsilon^m/\epsilon^{uc}$ | Ratio of the antigenic-sampling rate of ${y_i}^{L,m}$ to ${y_i}^{L,uc}$ by M cells. Represents the selective bias of M cells for sampling IgA-bacteria complexes. | [1,100] | 10 | Unitless | -2.1382 | 2.1544 | 7.4271 |
| $\tau^\delta$ | Multiplier to adjust the incremental increase in the selection threshold calculated in Eqn. S1.2.8. | [0.069, 0.945] | 0.42 | Unitless | 0.5941 | 0.6282 | 2.2591 |
| $C_n$ | Amplitude of the exponential function describing the diminishing pool of naïve T and B cells. | [0.005, 0.1] | 0.03 | Unitless | -0.4487 | 0.4564 | 2.8103 |
| $c_n$ | Decay rate of the exponential function describing the diminishing pool of naïve T and B cells. | [2, 30] x 1e-4 | 6.34 x 1e-4 | Unitless | -0.1466 | 0.324 | 2.0277 |
| $th_{range}$ | The plasma cell differentiation range, where $th_{high} = 1 - th_{range}$ and $th_{ang} = 1 + th_{range}$. | [0.01, 0.50] | 0.25 | Unitless | 9E-04 | 0.2928 | 1.6526 |
| $\alpha_{BC}$ | Invasiveness of *BC*. | [0.0069, 0.0946] | 0.036 | Unitless | 0.2107 | 0.2118 | 0.7995 |
| $\alpha_C$ | Invasiveness of *C*. | [0.0069, 0.0946] | 0.038 | Unitless | 0.1844 | 0.1905 | 0.6545 |
| $\tau^c$ | Multiplier to calculate the additional standard deviation in BCR affinity distribution after somatic hypermutation. | [6.90, 94.6] | 28.33 | Unitless | 0.0095 | 0.1632 | 0.8823 |
| $\kappa_E$ | Relative immunostimulatory capacity of *E*. | [215.9, 395.4] | 358 | Unitless | 0.0793 | 0.1334 | 0.6623 |
| $\psi^m$ | Activation rate of naïve B cells per unit of SIgA-antigen complex. | [0.01, 1] | 0.1 | 1/(Cells/gLC) | -0.1028 | 0.1194 | 0.3938 |
| $th_{apop}$ | Selection threshold multiplier determining the minimum BCR affinity to get sufficient T cell help to avoid apoptosis. | [0.01,0.49] | 0.25 | Unitless | -0.0039 | 0.0616 | 0.3263 |
| $\alpha_B$ | Invasiveness of *B*. | [0.0014, 0.0189] | 0.008 | Unitless | 0.0508 | 0.0528 | 0.2423 |
| $C_I$ | Maximum secretion capacity of plasma cells. | [5.43, 15.7] | 12.39 | Unitless | -0.0451 | 0.0508 | 0.1541 |
| $t^m$ | Time of M cell activation and the start of antigenic sampling. | [30, 154] | 129 | Days | 0.036 | 0.0411 | 0.5526 |
| $\tau^{new}$ | Multiplier to calculate the additional standard deviation in BCR affinity distribution in newly activated B cells. | [108.5, 314.0] | 180.21 | 1/(Cells/gLC) | 0.0197 | 0.027 | 0.1761 |

The mean absolute elementary effect ($\mu^*$) values represent the overall sensitivity of each parameter, quantifying the average impact that changes in the parameter have on the model's output. Since the relative ranking of $\mu^*$ values is key for identifying the most sensitive parameters, we used the 50th percentile as a threshold to distinguish the most influential ones (Fig. S6). Parameters with a $\mu^*$ above this threshold, $\{\epsilon^m/\epsilon^{uc}, \tau^\delta, c_n, C_n, th_{range}, \alpha_{BC}, \alpha_C\}$, are considered to be in the top 50%

of sensitivity, representing the most influential factors driving model behavior. These parameters describe the ratio of the antigenic-sampling rate of masked and uncoated bacterial antigens by M cells ($\epsilon^{m}/\epsilon^{uc}$), the multiplier to adjust the incremental increase in the selection threshold during GC reactions ($\tau^{\delta}$), decay rate ($c_n$) and amplitude ($C_n$) of the exponential function describing the diminishing pool of naïve T and B cells, the plasma cell affinity differentiation range ($th_{range}$), and the invasiveness of *Bacteroidaceae*, and *Clostridiales* ($\alpha_{BC}, \alpha_{C}$). Table S6 provides detailed explanations of these highest-ranked parameters from our global sensitivity analysis and discusses their impact on model outcomes to improve the interpretability of the fundamental model dynamics.

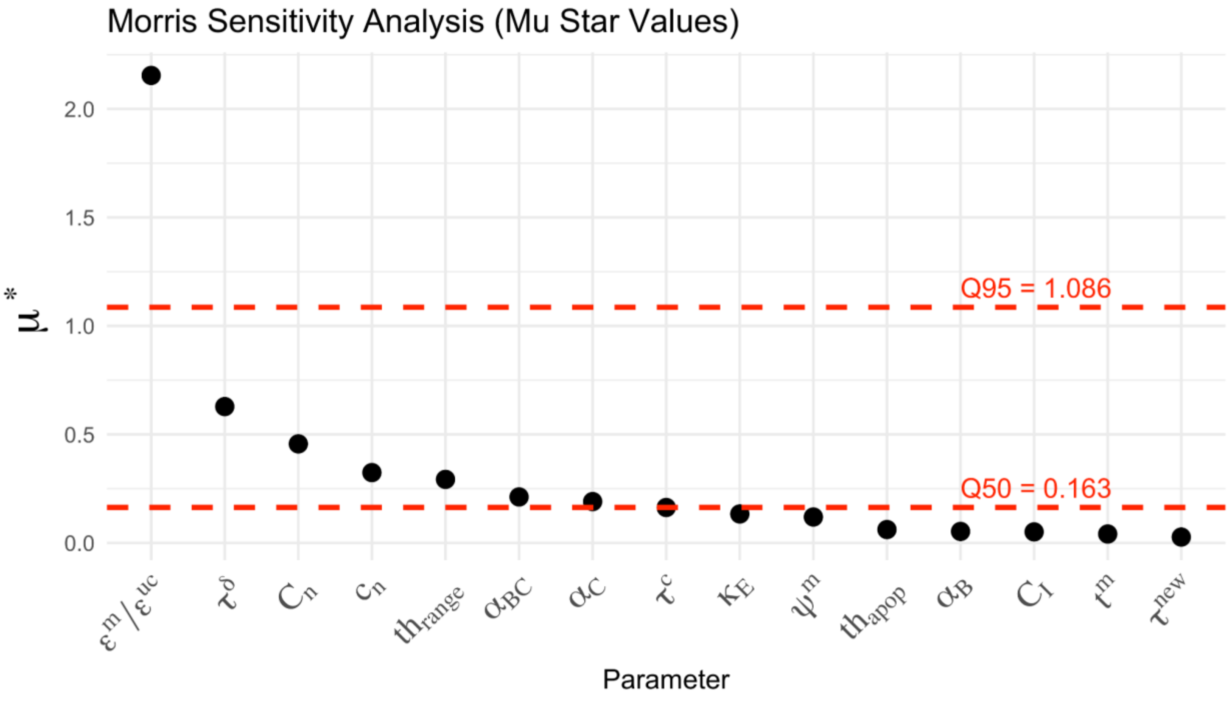

**Fig S6. Visualization of the global sensitivity analysis.** Visualization of the global sensitivity analysis presented in Table S5. Red dashed lines represent the 50th and 95th percentiles to distinguish the most influential parameters.

These parameters – $\epsilon^{m}/\epsilon^{uc}$, $\tau^{\delta}$, $C_n$, $c_n$, $th_{range}$, $\alpha_{BC}$ and $\alpha_{C}$ – are then examined in isolation through local sensitivity analyses (Figs. 6, S10, S8, and S7), alongside other model parameters. These analyses allow us to assess how localized variations in each parameter affect the model's output while holding the others constant, providing detailed insights into the individual impact of these parameters on endogenous affinity values.

**Table S6. Interpretation of the highest-ranked parameters in global sensitivity analysis and their impact on model dynamics.** This table presents parameters ranking above the 50th percentile (Q50) in our global sensitivity analysis (Fig. S6). For each parameter, we provide its biological interpretation, functional role within the model, and its quantitative influence on model outcomes.

| Notation | Description | Interpretation |
|---|---|---|
| $\epsilon^m/\epsilon^{uc}$ | The ratio of the antigenic-sampling rate of masked and uncoated bacterial antigens by M cells. | This ratio represents the selective bias of M cells for sampling IgA-bacteria complexes. As $\epsilon^m/\epsilon^{uc}$ increases, it promotes higher accumulation of IgA-coated bacterial antigens relative to uncoated ones in the GALT inductive sites. This leads to the relatively higher activation of tolerogenic dendritic cells compared to inflammatory dendritic cells, and consequently leading to a lower Tfh:Tfr ratio. This ratio, proxied by the ratio of uncoated to total antigens accumulating in the inductive sites in Eqn. S1.2.8, directly influences the selection threshold parameter $\delta_i$ that governs the scale of affinity maturation, thus directly influencing the development of endogenous affinity levels. Note that the effect of $\epsilon^m/\epsilon^{uc}$ is not taxon-specific. A detailed analysis of the impact of $\epsilon^m/\epsilon^{uc}$ is presented in Fig. 6. |
| $\tau^\delta$ | Multiplier to adjust the incremental increase in the selection threshold calculated in Eqn. S1.2.8. | This parameter directly influences the selection threshold value $\delta_i$ (Fig. 2C). Biologically, it represents how "stringent" the immunological signals (such as Tfh:Tfr cell ratio and the inflammation level) in the germinal center environment are in selecting B cells based on their receptor affinity. Higher values of $\tau^\delta$ indicates the immune system is exerting stronger selection — allowing B cells with higher affinity for antigen receive sufficient help to survive and proliferate–, leading to faster increases in BCR affinity per P-SHM-S cycle.<br><br>This parameter can vary based on factors affecting costimulatory molecule expression levels, T-B cell interaction quality, and the architecture of germinal center microenvironments. This parameter can be inferred from experiments that track the distribution of BCR affinities over multiple time points in controlled germinal center reactions where all other variables (antigen availability, inflammatory cytokines, initial BCR affinity distribution, and Tfh:Tfr ratio) are either also measured or standardized, potentially through immunization experiments. Single-cell technologies tracking the fate of individual B cell clones combined with mathematical modeling could estimate this parameter by fitting observed affinity trajectories. Note that this parameter represents an intrinsic selection rate of the GC reactions, meaning that it is not taxon-specific. |
| $C_n$ | Amplitude of the exponential function describing the diminishing pool of naïve T and B cells. | This parameter is a proxy for the initial size of the naïve T and B cell pool. As $C_n$ increases (decreases), there will be more (less) naïve B cells migrating from the bone marrow to the GALT inductive site, being activated by bacterial antigens, and turning into circulating GC B cells at each round of proliferation, somatic hypermutation, and selection (P-SHM-S) cycle. A higher influx of newly activated cells broadens the BCR affinity distribution of the circulating GC B cells – effectively introducing lower-affinity clones into the pool – which reduces the average BCR affinity falling within the selection thresholds, thus lowering the average endogenous SIgA (eSIgA) affinity. Conversely, if $C_n$ becomes too low (approaching 0), the number of cells available to differentiate into circulating or plasma cells may be insufficient, leading to negligible eSIgA secreting plasma cells. However, such extreme cases were excluded from our sensitivity analysis; the range for $C_n$ was chosen to ensure that the lower bound still supports circulating and plasma cell differentiation. Since the abundance of bacteria is regulated by eSIgA concentration and affinities, which in turn regulates the immunostimulatory tone of the microenvironment, value of $C_n$ influences the affinity maturation process through ecological and inflammatory feedback loops. |
| $c_n$ | Decay rate of the | This parameter determines how fast the naïve T and B cells differentiate into |

| | | |
|---|---|---|
| | exponential function describing the diminishing pool of naïve T and B cells. | effector cells. As $c_n$ increases (decreases), naïve cells will be depleted faster (slower), leading to a faster (slower) convergence of the BCR affinity values of the endogenous plasma cells. A faster convergence means that the host will exhaust its supply of naïve cells earlier, prematurely ending GC reactions and affecting the final BCR affinity values of endogenous plasma cells. With slower convergence, naïve cells remain available longer, allowing GC reactions to continue and potentially achieve different BCR affinity outcomes. |
| $th_{range}$ | The plasma cell differentiation range, where $th_{high} = 1 - th_{range}$ and $th_{ang} = 1 + th_{range}$. | This parameter defines the relative width of the affinity window around the selection threshold ($\delta_i$) that leads to plasma cell differentiation (Fig. 1C). When set to a value of 0.2 (20%), for example, cells with BCR affinities between 80% and 120% of the threshold value will differentiate into plasma cells. Larger values create a more permissive selection process, while smaller values enforce stricter selection pressure on the circulating B cells. Therefore, this parameter affects both the range of BCR affinity and the number of circulating B cells turning to plasma cells at each round of proliferation, somatic hypermutation, and selection (P-SHM-S) cycle. |
| $\alpha_i$ | Relative invasiveness of taxon $i$. | This parameter defines the taxon's ability to penetrate intestinal epithelial cells. As this value increases, the SIgA-**uncoated** bacterial load in the GALT inductive sites increases, which in turn increases the converged endogenous affinity level against the taxon. To reduce the impact of identifiability problems during inference, we fixed the invasiveness of *Enterobacteriaceae* to 1 ($\alpha_E = 1$) and inferred the invasiveness of the symbiotic commensals *Bifidobacteriaceae*, *Bacteroidaceae*, and *Clostridiales* relative to the pathogenic taxon *Enterobacteriaceae*. Value and effects of $\alpha_i$ are taxon-specific, meaning that modulating $\alpha_i$ directly impacts the affinity maturation against taxon $i$ via increasing the cumulative uncoated to total bacterial antigen ratio in Eqn. S1.2.8. However, since altering the endogenous affinity against a specific taxonomic group affects not only its own abundance but also the abundance of other taxa through community interactions, it can produce broader indirect effects. |
| $\tau^c$ | Multiplier to calculate the additional standard deviation in BCR affinity distribution after somatic hypermutation. | This parameter adjusts the standard deviation of the BCR affinity distribution of circulating B cells after each round of somatic hypermutation (Fig. 1, step 5) .<br><br>If $\tau^c$ is too high: The affinity distribution of B cells flattens, increasing the probability of extremely high or low affinity B cells. This reduces the proportion of cells falling within the selection thresholds, resulting in fewer B cells differentiating into plasma cells or continuing circulation.<br><br>If $\tau^c$ is too low: The B cell population may lack any clones within the plasma cell differentiation range (between $th_{high}\delta_i$ and $th_{ang}\delta_i$, Fig. 1, step 4) or circulation range (between $th_{apop}\delta_i$ and $th_{high}\delta_i$, Fig. 1). This outcome is biologically implausible since the selection pressure imposed by the T cells in the GCs is competitive and operates on a relative basis, resembling a rank-based selection process.<br><br>Therefore, this parameter requires careful calibration based on either:<br><br>1. Known plasma cell numbers and affinities from experimental data, or<br>2. Known taxonomic abundances, by observing the feedback loop between IgA and microbial populations (as done in our calibration process). |

***Local Sensitivity Analyses***

**Sensitivity to sampling rates for SIgA-antigen immune complexes and SIgA-free antigens by M cells (Eqns. S1.1.6-S1.1.7, parameters $\epsilon^{m}, \epsilon^{uc}$):** $\epsilon^{m}$ and $\epsilon^{uc}$ represent the antigenic-sampling rate of SIgA-bound (masked, ${y_i}^{L,m}$) and SIgA-free (uncoated, ${y_i}^{L,uc}$) antigens by M cells for each taxon $i$, respectively (Table S3). $\epsilon^{m}/\epsilon^{uc}$ is identified as the most influential parameter by the global sensitivity analysis. The $\epsilon^{m} > \epsilon^{uc}$constraint in our baseline scenario reflects the well-characterized role of M cells in creating a positive feedback loop for the sampling of SIgA-antigen immune complexes [18,101,102]. Varying the ratio of these rates ($\epsilon^{m}/\epsilon^{uc}$) allows us to explore the importance of this M cell mediated feedback loop in impacting how 'immune education' unfolds (Fig. 6). When $\epsilon^{m}/\epsilon^{uc} = 1$, i.e., when M cells do not distinguish between the SIgA-bound and SIgA-free antigens and sample SIgA-bound antigens with a rate **as low as** the SIgA-free ones, affinity levels against symbiotic commensals (*Bifidobacteriaceae*, *Bacteroidaceae* and *Clostridiales*) increase compared to the baseline case $\epsilon^{m}/\epsilon^{uc} = 10$, approaching values close to 1 for *Bacteroidaceae* and *Clostridiales*, reflecting a bias toward predominantly neutralizing rather than non-neutralizing masking behavior (compare  the type of SIgA function relative to the level of antibody affinity, Fig. 2B). The selective bias of M cells for sampling IgA-bacteria complexes promotes the production of non-neutralizing masking endogenous SIgA for symbiotic commensals, which are consequently presented as SIgA-immune complexes to the immune system, resulting in a feedback loop favoring tolerogenic imprinting.

**Sensitivity to the invasiveness of symbiotic commensals (parameters $\alpha_{\boldsymbol{B}}, \alpha_{BC}, \alpha_{C}$**): $\boldsymbol{\alpha_i}$ is the invasiveness parameter, representing the bacteria's ability to penetrate intestinal epithelial cells. As invasiveness increases, a larger proportion of the bacterial population reaches the GALT inductive sites without being coated by SIgA. A higher uncoated bacterial load activates more dendritic cells and skews the local immune environment toward stronger Tfh cell imprinting. Consequently, the Tfh:Tfr ratio increases, promoting greater endogenous SIgA affinity maturation against the respective taxon (Fig. S7). Among the invasiveness parameters, $\alpha_{BC}$ and $\alpha_{C}$ are identified as the 6th and 7th most influential parameters in the global sensitivity analysis, with importance values modestly exceeding the median across all parameters (Fig. S6).

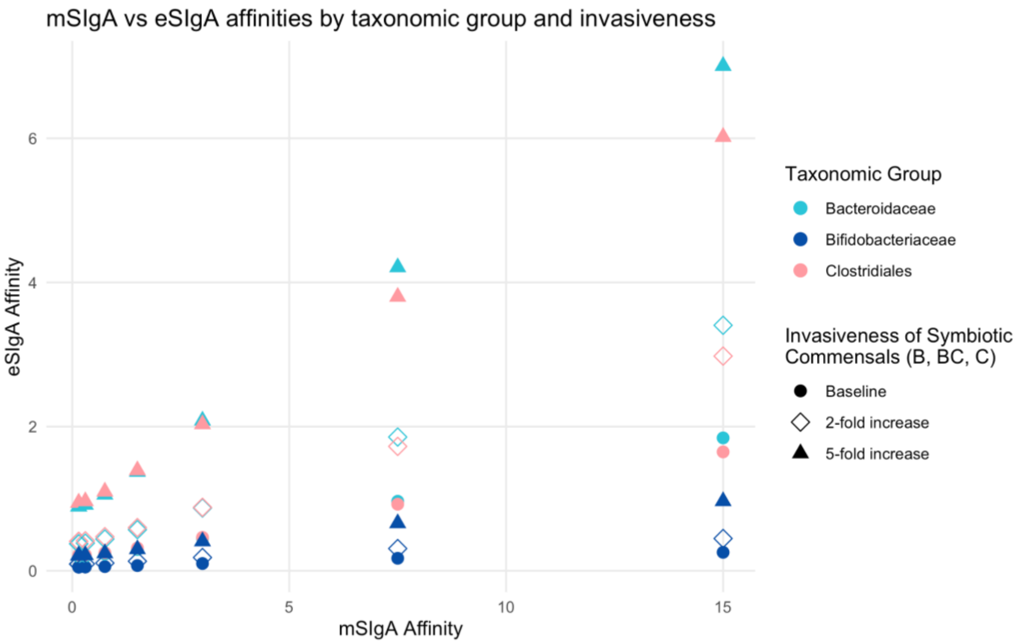

**Fig S7. Sensitivity analysis to the invasiveness parameter.** Endogenous versus maternal affinity levels (eSIgA vs. mSIgA) across taxonomic groups and varying levels of invasiveness. Each point represents the converged eSIgA affinity level for a given taxon under different levels of invasiveness, with shape denoting the degree of invasiveness (baseline, 2-fold, or 5-fold increase) and color indicating taxonomic group (*Bacteroidaceae*, *Bifidobacteriaceae*, *Clostridiales*). As invasiveness increases, uncoated bacteria more readily access GALT inductive sites, resulting in enhanced dendritic cell activation and a higher Tfh:Tfr ratio, which promotes increased eSIgA affinity.

**Sensitivity to the ranges that determine the T cell help for different B cell fates (*Modeling the role of Germinal Centers*, parameters $th_{apop}, th_{high}, th_{ang}$):** Parameters $th_{apop}, th_{high}$, and $th_{ang}$ determine the ranges for B cell apoptosis, circulation, and differentiation to plasma cells, respectively; and together they reflect the selection pressure imposed by the T cells in the germinal centers (reflected as a rank-based selection process). A sensitivity analysis on these ranges keeping all else calibrated for the baseline scenario illustrates the range of potential outcomes resulting from variations in the feedback loops between Tfh and Tfr cells, either leading to a more generous (wider BCR affinity range) or stringent (narrower BCR affinity range) selection. To do so, we first define the plasma cell differentiation range ($[th_{high}\delta_i, th_{ang}\delta_i]$) symmetrically around $\delta_i$ ($\delta_i$ denotes the antigen-specific selection threshold parameter representing the cumulative T cell help that determines various B cell fates, including apoptosis, circulation, or plasma cell differentiation; illustrated in Fig. 2C.) using only one parameter $th_{range}$, where $th_{high} = 1 - th_{range}$ and $th_{ang} = 1 + th_{range}$ (note that in Table S3 $th_{high} = 0.75$ and $th_{ang} = 1.25$, leading to $th_{range} = 0.25$). We then run our model for $0.05 \le th_{range} \le 0.75$ and $0.05 \le th_{apop} \le 0.75$ where cells with BCR affinity $< th_{apop}\delta_i$ fail to compete for T cell help, and die through apoptosis (note that in Table S3 $th_{apop} = 0.25$). We apply the constraint $1 - th_{range} \ge th_{apop}$ to ensure non-overlapping ranges defining the probability of continued circulating and probability of terminal differentiation to plasma cells.

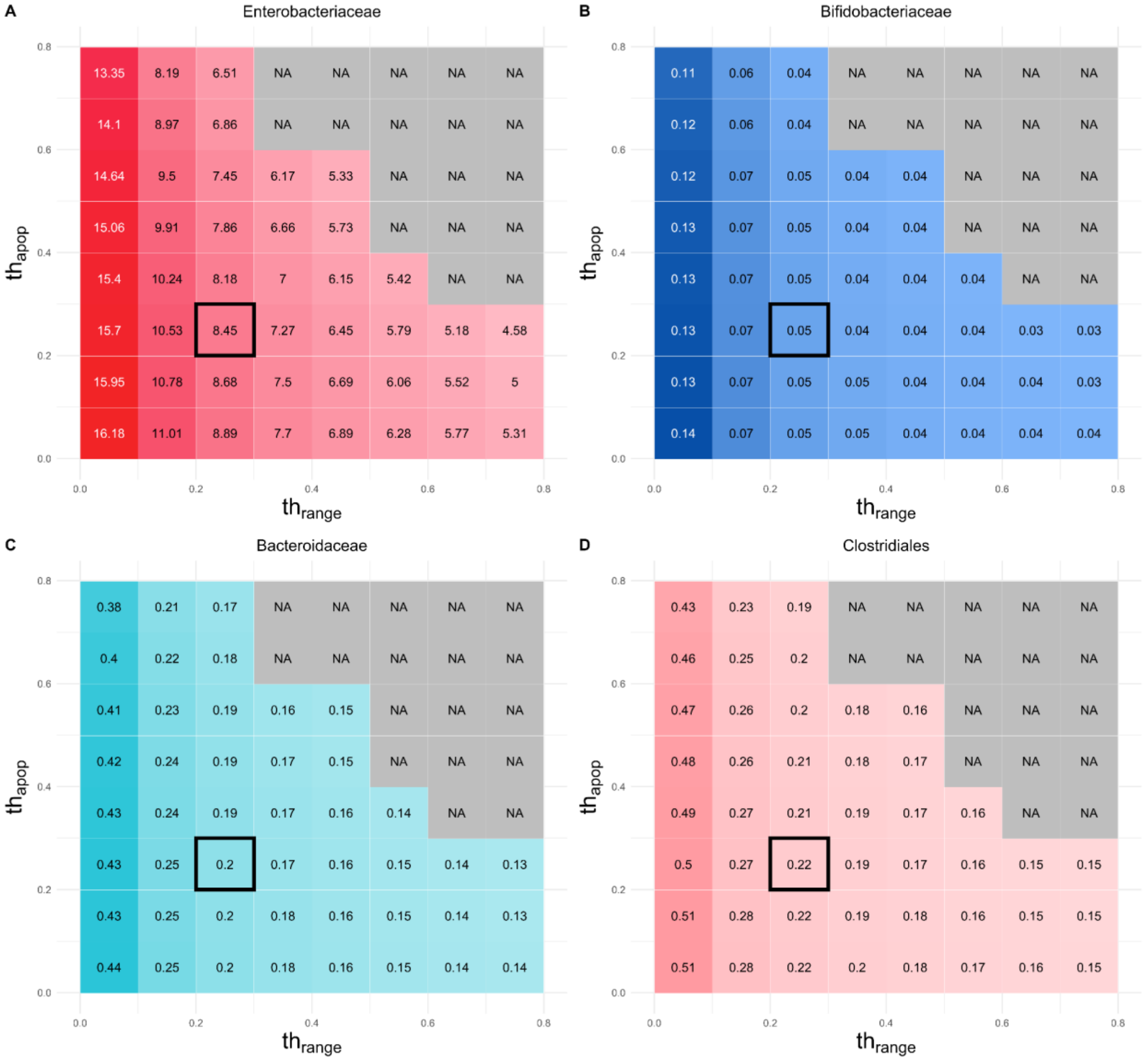

**Fig S8. Sensitivity analysis to the ranges that determine the T cell help for different B cell fates.** Heatmaps of average endogenous SIgA (eSIgA) affinity values at the end of 735 days (2 years) for $0.05 \leq th_{range} \leq 0.75$ and $0.05 \leq th_{apop} \leq 0.75$ for **A)** *Enterobacteriaceae,* **B)** *Bifidobacteriaceae*, **C)** *Bacteroidaceae* and **D)** *Clostridiales*. Colors represent the magnitude of the eSIgA affinity values, with darker colors indicating larger values. Numerical values are indicated in each box. NA represents $\{th_{range}, th_{apop}\}$ combinations with $1 - th_{range} \geq th_{apop}$ constraint. Baseline values ($th_{range} = 0.25$ and $th_{apop} = 0.25$) are indicated with the bold black boxes. Note that all affinity values targeting the symbiotic commensals (*Bifidobacteriaceae, Bacteroidaceae, Clostridiales)* are below 1, reflecting their predominantly masking behavior. This figure demonstrates the robustness of our affinity maturation model, showing that the exact numerical values of $th_{apop}$ and $th_{range}$ do not affect the functional properties of the endogenous antibodies.

**Sensitivity to the time point of M cell maturation relative to the steroid levels in the breastmilk (parameter $t^m$):** Timing of M cell maturation (parameter $t^m$) was calibrated based on qualitative information in the literature (see main text). However, it is still not clear when exactly mature M cells emerge in human infants. To assess the sensitivity of our model to parameter $t^m$, we varied $t^m$ from 30 to 154 days (time of weaning), all else being kept as in the baseline scenario (Fig. S9).

Early opening of M cells, when the *Enterobacteriaceae* load is still high in the gut lumen, leads to a higher bacterial antigen burden being recovered from the GALT inductive sites (Fig. S9C), making the infant more susceptible to enteric infections. However, this high antigenic load triggers a more aggressive affinity maturation process against *Enterobacteriaceae*, increasing the eSIgA affinity levels targeting it (Fig. S9A), and thereby reducing the cumulative *Enterobacteriaceae* burden in the gut lumen over 2 years (735 days). Although this might initially seem more favorable for the host, our model does not incorporate the detrimental downstream systemic effects of enteric infections that might result from the increased *Enterobacteriaceae* burden recovered from the GALT inductive sites (Fig. S9C). Moreover, our model assumes that M cell maturation coincides with the ontogenic timeline of other immune cells capable of initiating an endogenous immune response and producing high-affinity antibodies. In the event of a desynchronization between M cells and other immune components, the immune system would not be ready to initiate such a response, and the cumulative *Enterobacteriaceae* burden in the gut lumen would not decrease as much as shown in Fig. S9B. Immune responses against symbiotic commensals remain largely unaffected (Fig S9A), which aligns with our global sensitivity analysis that identified $t^m$ as the second least influential parameter. However, it is important to note that this analysis only considers the average eSIgA affinity against symbiotic commensals as its output metric, and does not capture the potential downstream effects of $t^m$ on high pathogenic load.

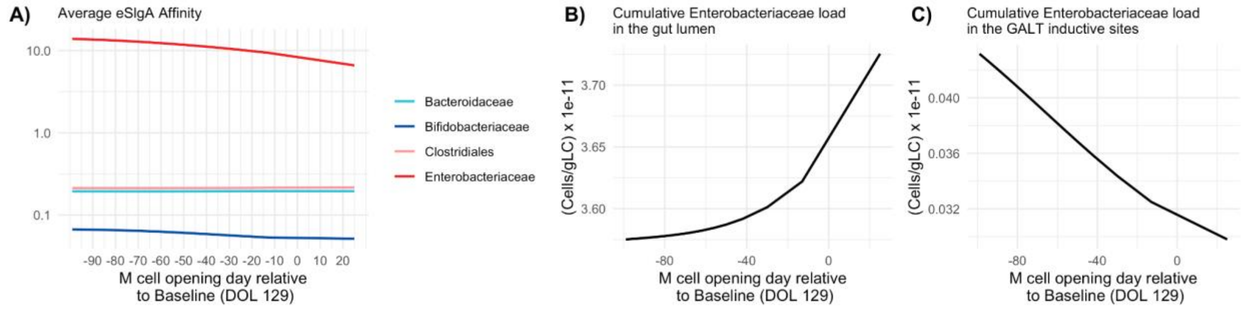

**Fig S9. Impact of early and delayed M cell opening on endogenous SIgA affinity maturation and *Enterobacteriaceae* load. A)** Average eSIgA Affinity values, **B)** Cumulative *Enterobacteriaceae* load in the gut lumen, and **C)** Cumulative *Enterobacteriaceae* load in the GALT inductive sites over the course of 2 years (735 days) for different delay durations of M cell opening relative to the baseline M cell opening time (DOL 129). DOL: Day of life. Early M cell opening increases *Enterobacteriaceae* antigen recovery in GALT inductive sites, potentially heightening susceptibility to enteric infections, yet triggers a more aggressive affinity maturation process against

*Enterobacteriaceae*, reducing their cumulative burden in the gut lumen over time. Immune responses against symbiotic commensals remain largely unaffected

**Sensitivity to the multiplier to adjust the incremental increase in the selection threshold calculation (parameter $\tau^{\delta}$):** Parameter $\tau^{\delta}$ is used in Eqn. S1.2.8 to adjust the incremental increase in the selection threshold $\delta_i$ in response to sampled antigens in GALT inductive sites. Biologically, it represents how "stringent" the immunological signals in the germinal center environment are in selecting B cells based on their receptor affinity (see Table S6 for detailed explanation). It is identified as the second most influential parameter by the global sensitivity analysis, supported by its impact on the eSIgA affinities against symbiotic commensals (*Bifidobacteriaceae*, *Bacteroidaceae* and *Clostridiales*) shown in Fig. S10A.

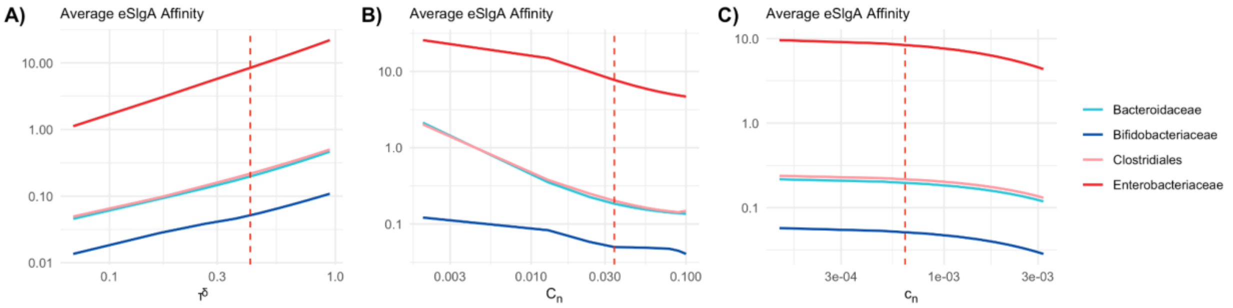

**Fig S10. Impact of parameters $\tau^{\delta}$, $C_n$, and $c_n$ on endogenous SIgA (eSIgA) affinities.** Sensitivity of the average endogenous SIgA (eSIgA) affinity values at the end of 735 days (2 years) in response to **A)** the multiplier to adjust the incremental increase in the selection threshold during GC reactions ($\tau^{\delta}$), and **B)** the amplitude ($C_n$) and **C)** the decay rate ($c_n$) of the exponential function describing the diminishing pool of naïve T and B cells. Red dashed line marks the baseline values. Note that both axes are shown on logarithmic (base 10) scales.

**Sensitivity to the decay rate and the amplitude of the exponential function describing the diminishing pool of naïve T and B cells (parameters $C_n$ and $c_n$):** $C_n$ denotes the amplitude, and $c_n$ denotes the rate of decay of the pool of naïve T and B cells as the host ages and as these cells undergo imprinting. This process is modeled as an exponentially decreasing function of time $t$, $C_n e^{-c_n t}$. $C_n$ and $c_n$ are identified as the third and fourth most influential parameters by the global sensitivity analysis, respectively (Fig. S6). While $C_n$ represents the initial size of the naïve T and B cell pool – determining how **many** naïve B cells migrate from the bone marrow to join the circulating B cell pool and thus the GC reactions at each time point –, $c_n$ governs how **fast** the naïve B and T cells become activated and differentiate: naïve B cells joining the GC reactions and naïve T cells differentiating into effector subsets such as Tfh or Tfr cells. Together, they shape the affinity maturation process (see Table S6 for detailed explanation). A high $C_n$ introduces a higher number of newly activated B cells into the light zones of the germinal centers at each time step, broadening the BCR affinity distribution—effectively lowering the average affinity due to the high influx of lower-affinity clones (Fig S10B). A high $c_n$ depletes the naïve pool more rapidly, accelerating convergence and prematurely terminating GC reactions, resulting in reduced BCR affinity values of endogenous plasma cells (Fig S10C).

***Illustrative validation using an external dataset***

To assess the validity of our model outputs, we compared them against an independent, external dataset. This analysis serves as an illustrative example of how model predictions in Fig. 3E can be externally evaluated by aligning them with real-world data trends. We examined the predictive power of microbial phyla and calprotectin levels in distinguishing celiac disease (CD, 2 subjects) from healthy controls (HC, 4 subjects) in infant gut microbiomes based on the data published by Olivares *et al.* [64]. We filtered subjects that were delivered vaginally, had no antibiotic exposure, and were breastfed until month 4, and used their samples collected at 6 months of age as predictors of disease state. Logistic regression models were used for each individual predictor (Abundances of *Proteobacteria*, *Actinobacteria*, *Bacteroidetes*, *Firmicutes*, and levels of Calprotectin), employing a cross-validation strategy with 3 folds and 100 repeats, while using upsampling to address class imbalance. Model performance was evaluated using Area Under the ROC Curve (AUC) with bootstrap-derived confidence intervals. Based on our model, we classified immunological outcomes as tolerant vs. not tolerant (affinities against all symbiotic commensals all below 1 simultaneously or not, respectively) and used this as a proxy for HC vs CD, respectively. We used 100 simulations where the breastfeeding duration randomly varied between 120 and 180 days to mimic the feeding patterns of the subjects. To represent a measurement taken within month 6, we sampled the predictor values at random days between day 150 and day 180. To calculate the AUC values at each iteration, we subsampled 6 subjects with 2 subjects being hyperreactive and 4 subjects being tolerant to resemble the data distribution. To prevent overfitting, we excluded any bootstrap iterations that produced AUC values of 1 during model performance evaluation. We then used the summary statistics of these AUC values. Our model showed good agreement with experimental data in terms of taxonomic rankings (Fig. S11A), with no rank differences across all five predictors. Both model and data (Fig. S11B) consistently identified Calprotectin as the strongest predictor (AUC of 1.00 in data and 0.73 on average in the model), which is expected given CD's inflammatory nature. More significantly, our model accurately reproduced the relative predictor importance patterns observed in the data across all taxonomic groups, providing a source of external validation for our model outcomes.

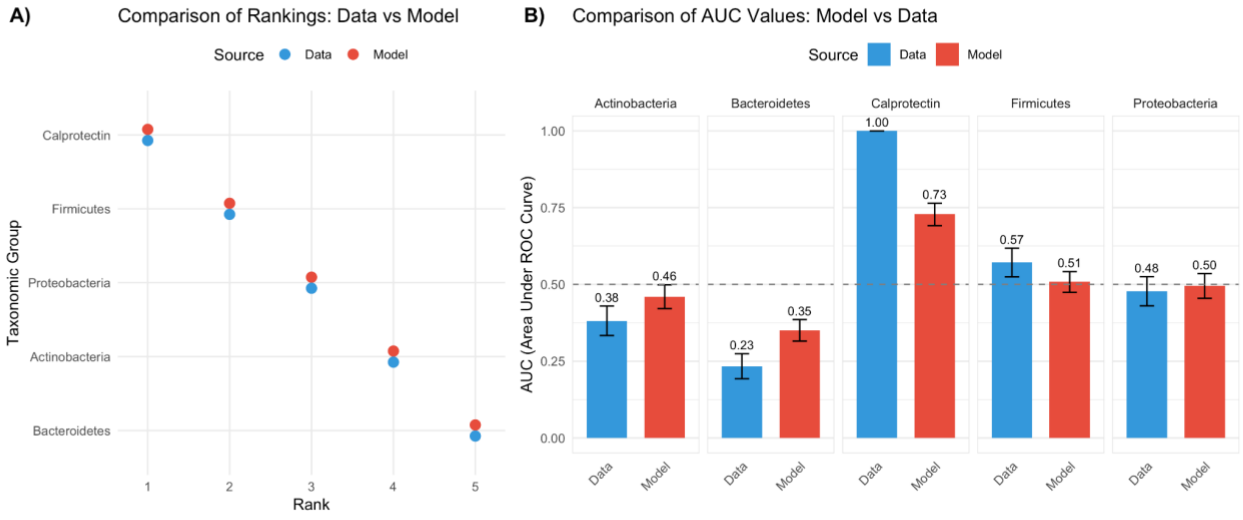

**Fig S11. Comparison of predictive performance between cohort data and computational model. A)** Comparison of rankings for predictors between experimental data (blue) and model predictions (red). Connected points indicate the same predictor, with horizontal position showing rank order (1-5) in predictive importance. Identical ranking of paired points demonstrates the strong agreement between the model and the cohort data. **B)** Comparison of AUC (Area Under ROC Curve) values for cohort data (blue) and model predictions (red). Error bars represent the standard error. Calprotectin shows the highest predictive power in both datasets, while taxonomic groups demonstrate a moderate predictive performance. The dashed horizontal line at 0.5 represents the threshold for random prediction.

***Model Limitations***

**Model Complexity:** Our framework addresses a high-dimensional question related to early life immunity. In developing the model, we inevitably made decisions about its complexity, constrained by the qualitative and quantitative data available in the literature. Certain mechanisms, such as the detailed dynamics of T-cell-B-cell interactions in the gut lymphoid tissues, the regulation of commensal microbiota by IgA, and the influence of the multiple active components in breastmilk on early immune development remain open questions in the literature. Consequently, we adopted the most widely accepted explanations, which are detailed alongside underlying evidence in the main text and methods. To enable feasible parameter inference, we simplified some biological processes, such as excluding explicit representations of certain germinal center (GC) reactions (e.g., interactions with antigen-presenting cells and B cell antigen capture for T-cell presentation), abstracting the feedback mechanisms between Tfh, Tfr, and B cells, and generalizing interactions between IgA and microbial taxa to focus on overall effects rather than detailed microbe-antibody dynamics. Capturing early life dynamics requires these abstractions to be integrated with explicit exogenous and endogenous processes that occur at specific points during development, leading to a high dimensional model. The resulting model frame – which may appear both overly simplistic and overly complex - reflects a balance between minimizing identifiability issues during parameter inference while ensuring that the essential processes are adequately represented.

**Memory B cell compartment:** Our model does not include a distinct memory B cell compartment due to insufficient quantitative data for parametrizing B cell fate decisions between memory and plasma cell formation. Memory B cells are particularly important given the preferential differentiation of naive B cells to memory B cells during early life [124] and the contribution of neonatally induced circulating memory B cells to continuously shape the gut mucosal immune response [151]. However, persistent humoral memory to gut pathogens and commensals can be maintained either by long-lived memory B cells replenishing dying plasma cells or by long-lived plasma cells themselves [155]. Given that the composition of the microbiome remains relatively stable, and the nature of antigenic stimuli is consistent throughout adulthood (the Steady Phase in our model), both mechanisms would produce mathematically similar outcomes. Since our model assumes a stable microbiome composition as the Steady Phase, we effectively represented the continuous replenishment of plasma cells by memory cells by assigning a plasma cell death rate of zero at steady state. This approach maintains a stable plasma cell population even in the absence of the stimulating bacterial antigen, indirectly reflecting the role of the memory compartment in sustaining long-term humoral immunity. While this simplification captures the overall effect of memory B cells on sustained antibody production, it does not explicitly model the dynamics of the memory B cell population, which remains a limitation of our current framework.

**B cell selection and antigen diversity:** We assume that B cell interactions with different antigens occur in isolation. While in reality competition for multiple antigens simultaneously could influence B cell selection, our model does not account for such interactions. By treating each antigen independently, we simplify the dynamics of affinity maturation, which may overlook potential synergistic or competitive effects in response to multiple antigens. However, quantitative data to sufficiently parametrize such interactions is currently lacking in the field.

**Role of M cells during early life in humans:** Our assumptions of the timing of M cell maturation and its significant role in antigenic sampling heavily relies on mouse models, and evidence for M-cell based antigenic sampling for humans during early-life is almost non-existent. However, there is substantial evidence for the presence and function of M cells in adult humans. Gullberg *et al.* [156] characterized M cells in human Peyer's patches, identifying specific cell adhesion molecules and demonstrating their role in facilitating antigen uptake. Furthermore, Hase *et al.* [157] identified glycoprotein 2 (GP2) as a specific transcytotic receptor on M cells in both mice and humans, suggesting a conserved mechanism across species. These studies provide strong evidence for the existence and functional importance of M cells in human intestinal tissue. Our sensitivity analysis on the timing of M cell maturation quantitatively explores the possibility that M cell development in human infants may not significantly limit the efficacy of antigen sampling, simulating a scenario where M cells begin antigenic sampling as early as day 30.

**Additional effects of SIgA on community composition and antigenic sampling:** In addition to its masking role in our model, SIgA plays a crucial role in promoting niche colonization and stabilizing the gut microbiota. By creating temporal and spatial niches along the gastrointestinal tract, SIgA enhances microbial diversity and fosters beneficial host–microbiota interactions [18]. Additionally, Rollenske *et al.* showed that SIgA coating protects the microbiome from environmental stressors like bile toxicity and bacteriophages [158]. In our model, SIgA's colonization-promoting effects could theoretically increase the net growth rate of commensals, represented by a scaling coefficient. This adjustment would weaken colonization resistance in scenarios with insufficient IgA (either maternal or endogenous), as commensals would experience reduced growth. To maintain the same model fit in cases of no breastfeeding (and thus lack of maternal SIgA), re-calibrating the competition terms—likely by enhancing the neutralization of pathogens by commensals—would be necessary to preserve the longitudinal microbial abundances in the gut lumen. Additionally, SIgA likely influences antigenic sampling by modulating microbial proximity to the gut epithelium, beyond its masking role and M cell adhesion bias (which are incorporated in our model). In our model, this effect would modify the sampling rate based on the change in a particular taxon's proximity to the epithelium. Unfortunately, the quantitative data required to accurately capture these effects remain unavailable in the current literature.

**Innate immune mechanisms:** Our model focuses on adaptive immune responses and necessarily omits several critical processes that change dynamically over early-life. In particular, exclusion of innate immune mechanisms means leaving out the tolerogenic bias of the Pattern Recognition Receptors (PPRs) and Dendritic Cells during infancy, known to direct T-cells towards a largely 'anti-inflammatory' response [159]. This phenomenon will influence the fate of commensal-specific T cells, eventually affecting the endogenous antibody response. In our model, the absence of this temporal tolerogenic bias is likely to be numerically (though not mechanistically) compensated by the combination of anti-inflammatory effects of maternal antibody masking and the inherent anti-inflammatory properties of symbiotic commensals. While the potential overestimation of these anti-inflammatory properties may affect the binary immune outcome of a tolerogenic or hyperreactive state given the duration of different feeding practices, the remainder of our analysis and qualitative results and conclusions should remain robust against such biases.

**Cross-reactivity of SIgA:** While we model antigen-specific SIgA responses independently across taxonomic groups, this does not account for evidence that SIgA can exhibit cross-species and even cross-phyla reactivity. Such cross-reactivity may arise from shared epitopes between species, similar glycan motifs present on taxonomically distant bacteria, or a combination of canonical and noncanonical binding [17]. These mechanisms allow individual IgA clones to bind to diverse microbial taxa, potentially blurring the boundaries of taxon-specific immune targeting. However, the mechanistic basis of this "cross-species reactivity" remains unclear [17]. Quantitative data on the range and specificity of SIgA binding across microbial communities are still limited, making it difficult to parameterize cross-reactive binding events in a mechanistically grounded way. While our antigen-specific modeling approach preserves model tractability and captures key dynamics of interest, it likely underestimates the masking effect of low-affinity SIgA binding across symbiotic commensals, representing an important avenue for future model refinement.

**Impact of LPS structures on immune education:** Our model treats LPS as a generic inflammatory signal, primarily focusing on the immunostimulatory effects of hexa-acylated lipid A derived from Escherichia-Shigella. However, experimental evidence, notably from Vatanen *et al.* [78], demonstrates that LPS molecules vary significantly in their immunogenic properties depending on their acylation pattern and bacterial origin. Penta- and tetra-acylated LPS produced by Bacteroides dorei, for example, are antagonistic to TLR4 signaling and actively suppress immune activation. These immunoinhibitory LPS types not only fail to induce endotoxin tolerance, but can also block the immune-educational effects of stimulatory LPS, as shown in both in vitro assays and mouse models of autoimmunity. Because our current model does not distinguish between LPS subtypes, it cannot fully capture scenarios in which tolerance is impaired due to insufficient immune stimulation. Future iterations of the model could incorporate functionally distinct LPS categories to explore both ends of the immune education spectrum—from hyperactivation-induced breakdown to under-stimulation-induced failure.

**Alternative Pathways for Immune Imprinting:** Our model focuses on the induction of microbiota-specific IgA via germinal center dynamics within organized mucosal inductive sites, such as Peyer's patches and colonic patches. These structures directly sample luminal antigens through M cells, are known to support affinity maturation and class switching in response to microbial colonization, and have been shown to be indispensable for generating taxon-specific IgA against commensal bacteria [160,161]. We do not explicitly incorporate parallel routes of immune activation, such as the goblet cell-associated antigen passage (GAP)–mesenteric lymph node (MLN) axis. This pathway is particularly relevant during pre-weaning, when GAPs facilitate the delivery of luminal antigens to tolerogenic dendritic cells in the lamina propria, which then migrate to MLNs to support regulatory T cell expansion and systemic immune imprinting [7]. While MLNs are capable of supporting germinal center formation [162], their role in microbiota-specific SIgA generation appears to be primarily modulatory. Both PPs and MLNs interact with dendritic cells and B cells, but PPs and colonic patches are more directly involved in the initiation of taxon-specific responses, whereas MLNs contribute to the broader regulation and dissemination of IgA-producing cells [162–164]. These mechanisms, while essential for establishing immune homeostasis, lie outside the specific scope of our current model.

**Gut permeability:** Temporal changes in the tight junctions and permeability of the gut epithelium are also elided in our framing. These factors influence the translocation of antigens, pathogens, and toxins from the gut lumen into the underlying tissues, increasing the risk of pathogen infiltration. The scenario of higher permeability of the gut epithelium during very early stages of ontogeny is partially recapitulated during our sensitivity analysis for the timing of M cell maturation and the initiation of antigenic sampling. Early opening of M cells — akin to compromised tight junctions and increased gut permeability during early ontogeny — allows a higher bacterial antigen burden to be recovered from GALT inductive sites. This elevated antigenic load initiates a more aggressive affinity maturation process against *Enterobacteriaceae*, thus recapitulating an exacerbated immune response against this taxon when gut permeability is compromised. The affinity maturation processes for the symbiotic commensals (*Bifidobacteriaceae*, *Bacteroidaceae*, and *Clostridiales*) are mostly unaffected, demonstrating the strong anti-inflammatory influence of the maternal antibody masking of these bacterial taxa. Similar to these results, implementing the temporal changes in gut permeability into our model is likely to increase the antigenic load recovered from the GALT inductive sites during the very early stages of ontogeny. However, the re-calibration of antigenic sampling rates and affinity maturation parameters such that the endogenous affinity levels still converge to maternal ones will counterweight this temporal bias. This adjustment would likely result in a steeper convergence of the affinity levels without affecting any other outcomes regarding the local antibody response.

**Functional shifts in gut microbiome:** Lastly, strain-specific shifts in the gut microbiome are not represented in our model, yet these shifts are known to affect metabolic and functional development. Our model represents bacterial influences on the microenvironment in a binary fashion (inflammatory or anti-inflammatory), a simplification that yields model tractability, but comes at the expense of generalizing the maturation of the antibody response towards a taxonomic group rather than specific strains. However, formally addressing the known distinct roles of certain bacterial strains in gut immune system development (*e.g.* [135]) and modeling their load and timing of inoculation would enhance our model's ability to address questions specific to the impact of hygiene practices, nutrition, and therapeutic use of probiotics on immune education.

**Impact of unconsidered exogenous variables:** While our model captures primary exogenous inputs that are well-characterized in early immune development, it does not account for other potentially influential factors, such as milk-derived EGF, lipids, viral exposures, or environmental stressors. These variables, though not included in our current model due to lack of robust quantitative data, could affect the immune landscape and modify the predicted outcomes. We suggest that future iterations of the model incorporate these variables, as data becomes available, to provide a more comprehensive analysis of early-life immune dynamics.

***Experimental Model Validation***

Our model is rooted in a mathematical formalization of well-described mechanisms, with the goal of predicting hard-to-intuit outcomes of emergent properties of mechanistic interactions. However, how the different mechanisms are combined in the model may lead to inappropriate weight to different processes, leading to mismatch between model predictions, and empirical outcomes, and this could be interrogated with simple experiments or cohort studies. We provide some plausible experiments below to test the predictions of our model.

#### 1. Longitudinal cohort study to validate predictive power of fecal samples

**Prediction:** Our model predicts that the predictive power of SIgA-bound Enterobacteriaceae abundance for downstream immune phenotypes follows a non-monotonic pattern over the course of ontogeny (Fig. 3E).

**Approach:** This can be tested in a longitudinal human birth cohort by collecting monthly fecal samples for 16S rRNA sequencing and IgA-seq from birth to two years of age, coupled with clinical follow-up for allergic or inflammatory outcomes. Machine learning models trained on different timepoints could assess how well microbial and SIgA-binding features at each stage predict immune outcomes. This would validate the model's proposed temporal window of maximal diagnostic value and inform optimal sampling strategies in clinical research.

#### 2. Probiotic supplementation as an intervention when breastfeeding is not possible

**Prediction:** Our simulations indicate that in the absence of maternal antibodies, the early introduction of key symbiotic commensals belonging to Bacteroidaceae and Clostridiales can reduce the pathogenic-to-commensal ratio up to 30 fold in the gut lumen (Fig. 5A). This finding supports the use of targeted probiotic supplementation as a practical intervention when breastfeeding is not possible.

**Approach:** In mouse models, neonates receiving exclusive complementary feeding could be administered defined probiotic cocktails, with outcome measures including Enterobacteriaceae abundance in fecal samples and inflammatory biomarkers such as calprotectin. Based on our model predictions (Fig. 5A), we would expect a marked reduction in the pathogenic-to-commensal ratio — potentially up to 30-fold — along with a steadily decreasing trend during the first 30 days of life. While the precise magnitude may vary, reproducing this directional shift would lend support to the proposed mechanism and provide a foundation for future translational work. Although further validation in human settings is needed, this experimental design offers a feasible approach to test whether targeted early-life supplementation can reshape gut ecology in accordance with model predictions.

#### 3. Attenuation of hyperreactivity in offspring of mothers with IBD or allergies

**Prediction:** Our model predicts that even when maternal SIgA displays hyperreactivity toward commensals—as may occur in mothers with IBD or allergic disorders—the immune system of the offspring does not simply replicate this reactivity, but exhibits an attenuated phenotype (Table 1, Fig. S4).

**Approach:** Murine models of maternal gut inflammation provide a tractable system to test this prediction. Dams with DSS-induced colitis—a condition known to alter intestinal SIgA reactivity—can be used to generate offspring exposed to hyperreactive maternal antibodies via breastfeeding. Fecal IgA-seq can be performed on both dams and their offspring across developmental timepoints to quantify the similarity of SIgA-binding profiles. Comparing the magnitude of hyperreactivity in maternal versus offspring SIgA responses will allow quantification of attenuation, which can be directly compared with model-predictions in Table 1. Complementary measurements of tolerogenic markers in the gut, such as regulatory T cell abundance and/or cytokine levels in the lamina propria, can further assess whether immune programming remains biased toward tolerance. Experimental confirmation of such attenuation would support the model's prediction that breastmilk, even from an immunologically dysregulated mother, retains regulatory properties that buffer against the vertical transmission of inflammatory phenotypes.

**4. Knockout experiments to test the significance of M cell sampling bias**

**Prediction:** Our model predicts that the bias in M cell-mediated antigen sampling — the preferential uptake of SIgA-coated bacteria over uncoated ones — is critical for driving the development of immune tolerance (Figs S6 and 6). This bias results in an overrepresentation of masked commensals in antigen presentation (since the neutralized ones will be expulsed from the lumen, they will not be sampled), promoting tolerogenic programming of dendritic cells, particularly via the C-type lectin receptor SIGNR1 [75] and regulatory follicular T cell responses. If this sampling preference is lost, the tolerogenic bias of the GC reactions will also be lost, increasing the risk of hyperreactivity.

**Approach:** While the selective adherence of M cells to SIgA is functionally well-established [73,74], the molecular identity of the responsible receptor remains unknown, making direct genetic knockout experiments unfeasible. However, it is not the sampling bias per se but its downstream effect — the conditioning of dendritic cells in the subepithelial dome — that ultimately drives tolerance. Therefore, our prediction can be tested using SIGNR1 knockout (Signr1–/–) mice, where DCs lack the receptor necessary for tolerogenic response to SIgA [165]. Neonatal mice can be colonized with a defined microbial consortium and provided maternal SIgA via breastfeeding. Comparing fecal IgA-seq profiles from Signr1–/– mice and wild-type controls, one can assess whether the absence of SIGNR1 disrupts the expected development of masking SIgA profiles and tolerance.

***Experimental Model Parametrization***

1. **Characterizing how antibody affinity levels map to bacteria coating rates (Eqns. S1.1.1-S1.1.2):** Differential coating rates of bacteria can be quantified by using IgA-Seq. To mathematically map coating rates to antibody affinity levels (as reflected in the model, Fig. 1B), Surface Plasmon Resonance (SPR) [166,167] or Bio-Layer Interferometry (BLI) [168] can be used to quantify the binding affinity of SIgA for bacterial antigens. Using this data for multiple symbiotic and pathogenic strains, one can quantify the function to map affinity levels to masking/neutralizing rates.

2. **Characterizing how the selection threshold (our quantification of T cell help determining the B cell fates) changes during affinity maturation:** The selection threshold in our model is a composite variable that reflects the outcome of multiple selection processes occurring within the germinal centers. This approach aligns with the common practice in

mathematical modeling of pooling and abstracting detailed mechanisms into higher-level parameters. A selection bias toward an increased BCR affinity range when Tfh:Tfr ratio [169,170] and/or the inflammatory tone of the microenvironment [171] is increasing is well established in the literature. However, the mathematical nature of this relationship is assumed to be multiplicative in our model (Eqn. S1.2.8), with the addition of a factor to account for their differences in magnitude (Supplementary Material, *Immune Dynamics)*. Quantifying the nature of this relationship (additive, multiplicative, non-linear) is challenging, but several experimental approaches can be designed to investigate this hypothesis:

a. ***In vitro* germinal center reactions:** Organoid cultures or germinal center-like structures *in vitro* can be used, where one can control the levels of cytokines, chemokines, and T-cell interactions. With established protocols for *in vitro* germinal center B cell culture, we can elucidate the molecular mechanisms of GC B cell differentiation [172,173]. By varying levels of cytokines (e.g., IL-4, IL-21) and T-cell help (e.g., by adding Tfh cells or anti-CD40 antibodies), isolating B cells and using Surface Plasmon Resonance (SPR) to quantify their binding affinity at different time points, one can curate a dataset where the longitudinal relationship between the affinity levels, T cell help, and microenvironmental effects can be quantified.

b. **Animal models with conditional knockouts:** Similar to *in vitro* methods, we can apply conditional knockout mouse models of cytokine receptors (e.g. IL-21R) [174] or T-cell help (e.g. CD40-CD40L interactions) [175,176], isolate plasma cells from lamina propria at different time points during ontogeny, quantify the B-cell binding affinity as described in point a., and quantify the relationship.

c. ***Ex Vivo* Lymphoid Tissue Cultures:** *Ex vivo* engineered organotypic cultures have enabled the real-time study and control of biological functioning of mammalian tissues. A B cell follicle organoid made of nanocomposite biomaterials recapitulating the anatomical microenvironment of a lymphoid tissue that provides the basis to induce accelerated germinal center reactions has been already demonstrated in the literature [177,178]. Thus, another option is to culture explanted lymphoid tissues and manipulate the levels of microenvironmental factors (e.g., through cytokine treatments) and T-cell help (e.g., through anti-CD40 or anti-ICOS treatments), isolate the plasma cells at different time points, quantify the affinity as described in point a., and quantify the relationship.

3. **Quantifying the proliferation, somatic hypermutation, and selection (P-SHM-S) cycle:** To measure which B cells with specific BCR affinities are selected, and which have died during proliferation, somatic hypermutation (SHM), and selection in germinal centers (GCs), several methods can be employed.

   a. One can utilize single-cell RNA sequencing (scRNA-seq) as demonstrated by Corinaldesi *et al.* [179]. Their study used scRNA-seq to track immunoglobulin repertoire and transcriptomic changes in germinal center B cells, providing detailed insights into the selection and differentiation of B cells based on affinity maturation. This method allows for the precise characterization of B cell populations and their affinity profiles, which can be used to quantify how the distribution of BCR affinities change over the course of P-SHM-S cycles in our model.

b. Another alternative is to use EdU incorporation assays, as described by Biram and Shulman [180]. Their study demonstrates the use of EdU labeling to measure B cell proliferation *in vivo*, providing a detailed methodology for tracking B cell proliferation and selection within germinal centers using flow cytometry. This technique can quantify the impact of BCR affinity on cell survival and proliferation, which can be used to quantify the selection range for different B cell fates (points 1-7 under *Modeling the role of Germinal Centers*, Materials and Methods) that is used in our model.
